# Mapping forests with different levels of naturalness using machine learning and landscape data mining

**DOI:** 10.1101/2023.07.30.551142

**Authors:** Jakub W. Bubnicki, Per Angelstam, Grzegorz Mikusiński, Johan Svensson, Bengt Gunnar Jonsson

## Abstract

To conserve biodiversity, it is imperative to maintain and restore sufficient amounts of functional habitat networks. Hence, locating remaining forests with natural structures and processes over landscapes and large regions is a key task. We integrated machine learning (Random Forest) and open landscape data to scan all forest landscapes in Sweden with a 1 ha spatial resolution with respect to the relative likelihood of hosting High Conservation Value Forests (HCVF). Using independent spatial stand-and plot-level validation data we confirmed that our predictions (ROC AUC in the range of 0.89 - 0.90) correctly represent forests with different levels of naturalness, from deteriorated to those with high and associated biodiversity conservation values. Given ambitious national and international conservation objectives, and increasingly intensive forestry, our model and the resulting wall-to-wall mapping fills an urgent gap for assessing fulfilment of evidence-based conservation targets, spatial planning, and designing forest landscape restoration.

## Introduction

Remnants of naturally dynamic forest landscapes are key biodiversity hotspots providing habitats to a large number of species, forming intact and resilient ecosystems, and provisioning of multiple ecosystem-level services^1,2^. The scarcity and continuing loss of such areas^3^ has raised awareness about the need for forest conservation and forest landscape restoration^4^. In particular, primary and old-growth forests with low human impact form important local biodiversity hotspots^5–8^. Hence, there is a growing need for effective mapping of such areas to safeguard their existence through conservation and restoration, and as a foundation for nature restoration and spatial planning to ensure habitat network functionality^9–13^.

Forest species have adapted to a diversity of naturally occurring disturbance regimes^14^, which create structural patterns at multiple scales^15^ and form diverse habitats. The level of forest naturalness^16,17^ reflects a transition from naturally dynamic forests as complex ecosystems to simplified cropping systems managed for wood and biomass production^18^. Globally, such transitions are creating expanding frontiers of naturalness loss^19,20^.

Green infrastructure (GI) is an established concept addressing the urgency of conserving and restoring sufficient areas of functional and representative habitat networks^21–23^ in a multiple-scale approach to defining and mapping areas that need to be strictly protected or managed to favour ecosystem resilience, biodiversity and ecosystem services^24^. Planning and maintaining functional GI networks requires knowledge on the existence and location of necessary amounts of high conservation value areas able to maintain structural and functional connectivity across landscapes and regions^23–25^. In areas dominated by intensive cropping systems for production of wood biomass, both the provisioning of other ecosystem services and biodiversity conservation thus become challenging. The need to identify remnant natural forest patches triggered the use of the term forest naturalness^16^, and the establishment and mapping of High Conservation Value Forests (HCVF)^26^, generically capturing forests with high levels of naturalness ^27^. Typically, HCVF harbour native tree species, has a long history of forest continuity, vertical and horizontal structural complexity and low levels of anthropogenic influence^7,28,29^. Such HCVF mapping data can be used for assessing fulfilment of evidence-based conservation targets^23,30^, spatial planning^25,28^, and designing forest landscape and nature restoration^31^.

With a long history of intensive wood biomass production, Sweden has become a globally important producer of wood products. However, this has caused an overall transformation from naturally dynamic forests into effective wood production cropping systems^32,33^. Outside the Scandinavian mountain range foothill forests, only fragments of such naturally dynamic forests remain^12^. In Sweden, the first national-scale HCVF dataset was compiled in 2016, based on known and registered (at that time) forest biodiversity hotspots within and outside formally protected areas. The HCVF were mapped by field inventories during several decades and without a predefined sampling scheme^34^. Thus, the background information, based on costly and time-consuming field surveys, is neither up to date, nor comprehensive, and is limited in terms of its spatial and habitat representativeness.

The increasing availability of wall-to-wall spatial datasets describing multiple dimensions of landscapes with unprecedented thematic resolution and spatial precision creates an opportunity to overcome these limitations^35,36^. Examples of such datasets range from raw remote sensing data collected globally by modern, civil satellite missions such as Landsat 8^37^ and Sentinel-2^38^; sophisticated data products such as the high-resolution maps of global forest cover change^39^ and forest canopy height^40^; to LIDAR-based forest structure measurements^41^ and high-spatial and thematic resolution land cover/land-use maps^42^. Additionally, rapid developments in applying machine learning to ecology and conservation^43–45^ provide an opportunity to map the naturalness and thus conservation values of forests^46^. It is challenging to provide direct mechanical meanings to multi-scale spatial proxy variables describing forest naturalness, as they interact in complex and highly non-linear ways. However, machine learning, with its ability to flexibly and automatically find the best predictive patterns explaining the data, is promising for providing a robust solution to this problem. Big spatial data can be processed with machine learning to develop a contiguous and complex measure of the naturalness of forest ecosystems, accounting not only for simple tree cover dynamics but also for forest structural properties and composition of surrounding landscapes at different spatial scales^46^.

Given the high pressure on forest ecosystems, and the urgent need for spatially explicit information on remaining HCVF, the aim of this study was to provide validated mapping of forest areas crucial for maintaining biodiversity at a landscape level. Implemented and tested on Sweden’s forests, the methods we developed are potentially applicable to other forest regions.

We used Sweden as a case study for the following reasons: (1) Sweden’s forest area is the largest in the European Union (c. 280 000 km^2^), ranges from temperate deciduous, through boreal to subalpine ecoregions, and covers wide gradients in forest type and forest landscape configurations and ownership patterns; (2) Its natural forests and forest landscapes have been transformed into wood biomass cropping systems to a high degree, which makes identifying HCVF remnants critical^12,47,48^; (3) There is strong political pressure to further intensify wood biomass oriented forestry^49^, and (4); The systematic public HCVF field surveys have been terminated^50^.

We present a comprehensive framework for predicting the local occurrence of forests with different levels of naturalness using a data mining and predictive modelling approach. More specifically, we used the Random Forest (RF) machine learning algorithm and publicly available high-resolution spatial datasets describing landscape configuration, topography, forest structural properties and various socio-economic factors affecting landscape patterns and processes at multiple scales. We trained our models and tested their performance using the only existing, yet incomplete, national scale HCVF database. Finally, using a comprehensive set of independent spatial datasets, we validated the extent to which the predicted relative likelihoods of HCVF occurrence actually represent a gradient of forest naturalness and conservation values.

## Results

### Predicting naturalness as conservation value

We used data mining and predictive modelling^51^ to scan all forest landscapes in Sweden for the occurrence of High Conservation Value Forests (HCVF). This modelling approach was motivated by the inherent properties of HCVF, which are complex and high-dimensional with respect to, among other factors, the physical landscape, biodiversity, socio-economics, management and history of use. This complexity means that a complete set of wall-to-wall spatial data describing all relevant dimensions is unavailable and one has to rely on available spatial proxy variables. Focusing on the level of naturalness as a proxy for HCVF, our approach was to integrate many different data sources describing multiple dimensions of forest landscapes at multiple spatial scales, including the physical landscape, tree stands’ bio-physical structure, and socio-economic factors related to current and historical anthropogenic pressure on forests across the whole of Sweden (Table 2). Expecting highly non-linear and complex interactions among proxy variables (especially when exploring them at multiple scales), we used the machine learning classifier Random Forest (RF)^52^ to train our models and generate predictions. We predicted the relative likelihood of HCVF occurrence in c. 21.85 million 1-ha pixels which are dominated by forest (i.e. forest cover > 0.5) in all four study regions, representing c. 78 % of the total forest area in Sweden (Table 1). The results based on a 10-fold spatial cross-validation (SCV)^53–55^ show that all four RF models, trained and validated independently for each of the four study regions, fit the data well and have high predictive capabilities. The model performance metrics fall into the following ranges (metric name [range]): ROC AUC [0.89-0.90], PR AUC [0.84-0.89], Pearson R [0.66-0.68], Brier’s score [0.14-0.15] and MCC [0.61-0.62] (see Figure 2 and Table 3 for detailed statistics for each model).

**Figure 1.**
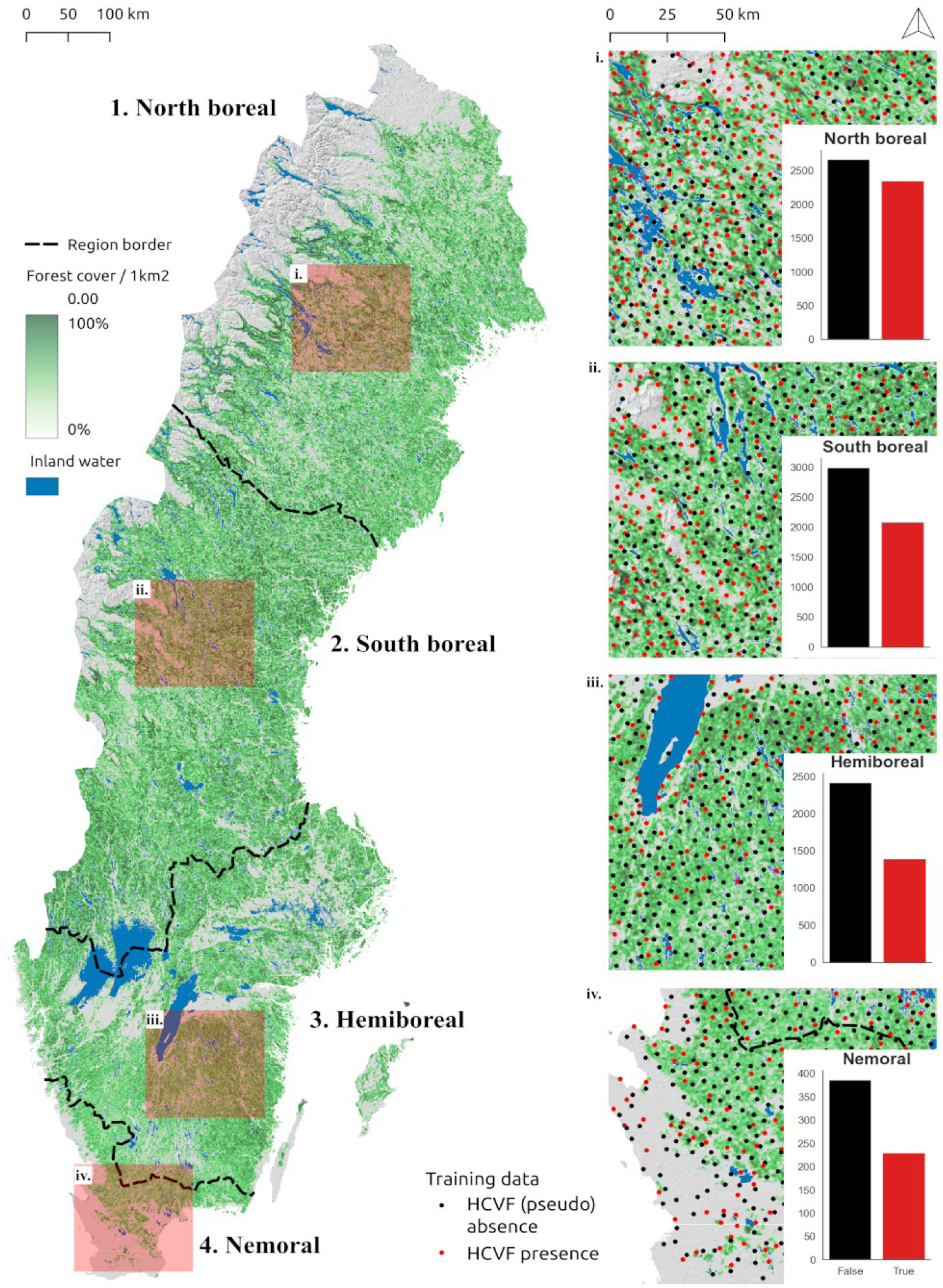
The study area (Sweden) divided into 4 regions (1: North boreal, 2: South boreal, 3: Hemiboreal, 4: Nemoral) and the illustration of the 1 ha forest pixel sampling density used for Random Forest models training and validation (red: HCVF presence, black: HCVF pseudo-absence). The barplots show the sampling distribution of presences (True) and pseudo-absences (False) for each region. The minimum distance between all sampling locations from the same category was set to 5 km, while the minimum distance between categories was set to 1 km. The forest cover is based on the Swedish national land cover data (NMD), but excluding temporarily non-forest areas (i.e. recently logged forest or young plantations).

**Figure 2.**
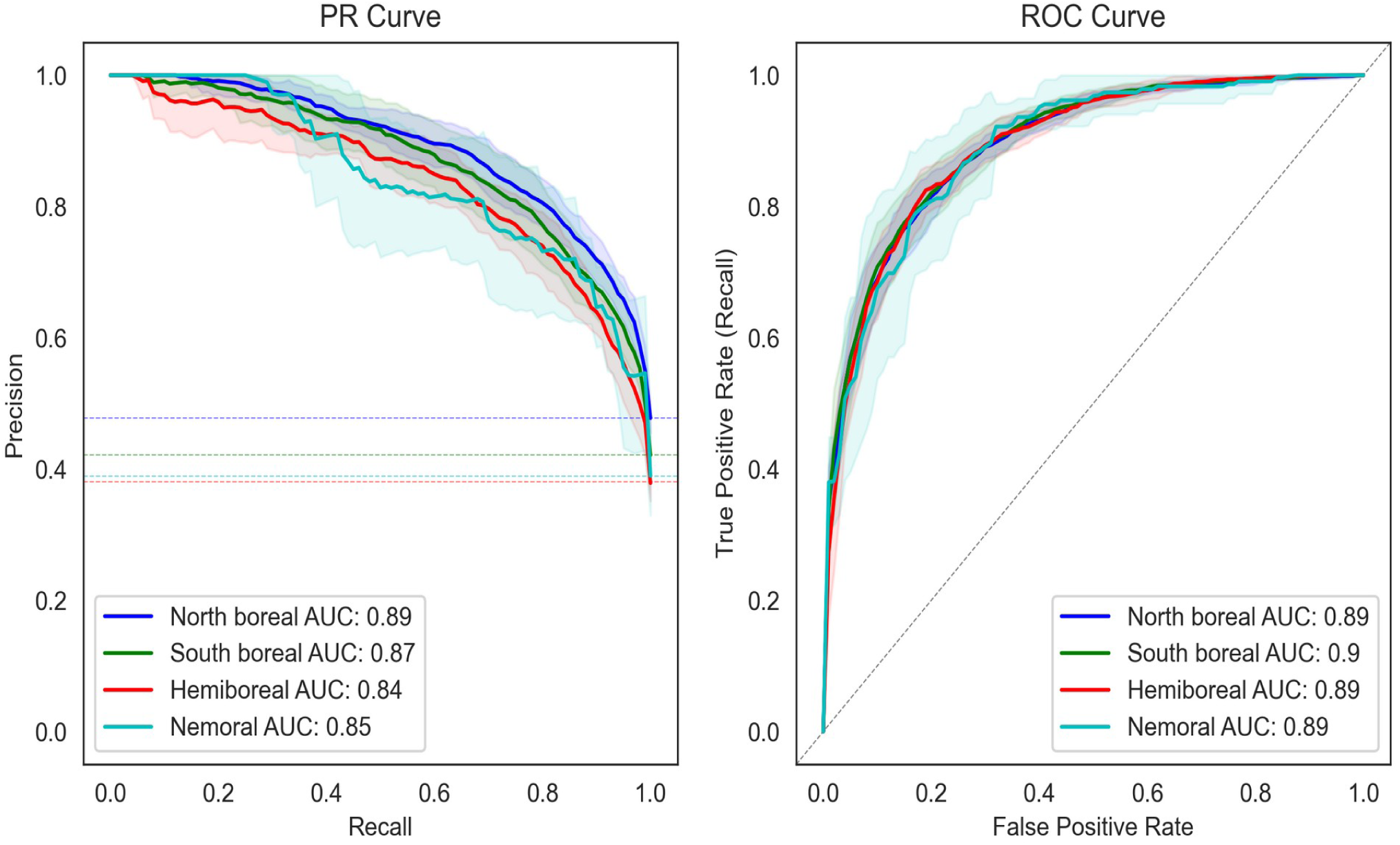
The performance of four Random Forest models trained independently for each study region as resulted from the 10-fold spatial cross-validation (SCV)^53–55^, and visualized as ROC Curve^97^ (right) and Precision-Recall Curve^97^ (left). For SCV we overlaid on the study area a grid of 20×20 km^2^ and randomly assigned all grid cells to different spatial subsets (i.e. folds) for the cross-validation procedure. The bold lines are based on the mean values of model performance metrics (calculated for each k-fold), while shaded areas indicate ± 1 SD. AUC is the area under the curve. The dashed-lines indicate a reference to no-skilled classifiers (for the PR Curve they correspond to the proportion of HCVF presence samples in each region).

**Table 1.** The basic land cover areal statistics (in kha) describing the study area (Sweden) divided into 4 regions (North boreal, South boreal, Hemiboreal and Nemoral). The “Total forest” area is the sum of “Current forest” (tree height > 5m), “Temporary non-forest” (recently logged or burned forest; tree height < 5m) and “Mountain woodland”. The “Mountain woodland” is the subalpine mountain birch (Betula pubescens ssp. czerepanovii) tree line forest^80^. The sources of data are the Swedish national land cover dataset (NMD) and High Conservation Value Forests database (HCVF).

|  | North boreal | South boreal | Hemiboreal | Nemoral | Total for Sweden |
| --- | --- | --- | --- | --- | --- |
| **Terrestrial land** | **15279.6** | **13465.7** | **10206.3** | **1921.1** | **40872.7** |
| Buildup | 7.3 | 17.5 | 54.8 | 18.5 | 98.1 |
| Roads | 130.9 | 231.8 | 330.6 | 80.5 | 773.8 |
| Wetlands | 2337.5 | 1191.6 | 349.7 | 48.8 | 3927.6 |
| Agriculture | 106.3 | 326.7 | 1572.1 | 577.1 | 2582.2 |
| Total forest | 9481.3 | 10500.6 | 6896.0 | 945.0 | 27822.9 |
| **Current forest** | **7157.6** | **8010.5** | **5673.8** | **767.1** | **21609** |
| Temporary non-forest | 1579.5 | 2279.7 | 1222.2 | 177.9 | 5259.3 |
| Mountain woodland | 744.2 | 210.4 | 0.0 | 0.0 | 954.6 |
| Formally protected forest | 1522.5 | 484.6 | 255.6 | 38.5 | 2301.2 |
| **HCVF** | **1960.4** | **1072.7** | **351.9** | **48.8** | **3433.8** |

**Table 2.**
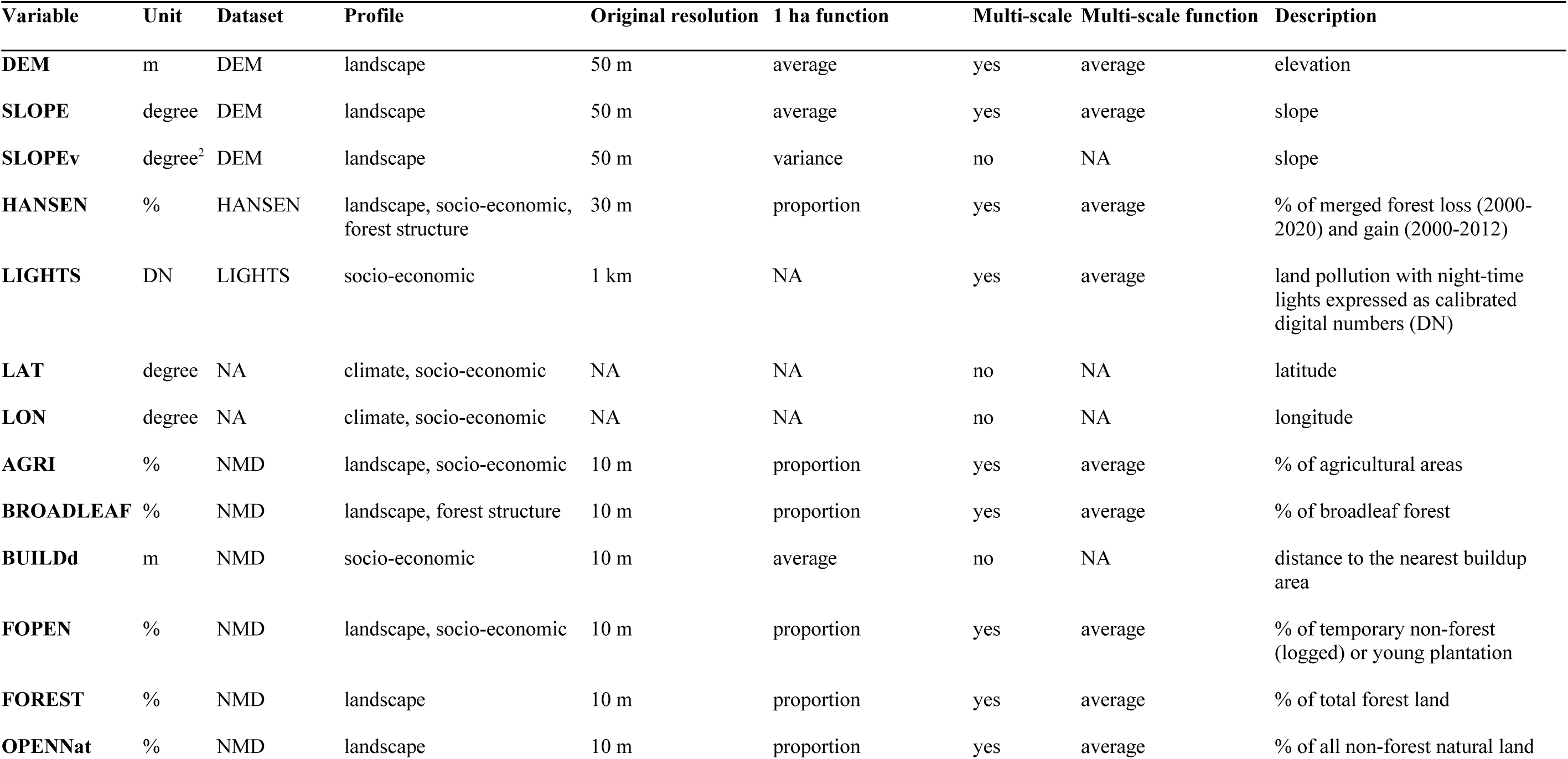
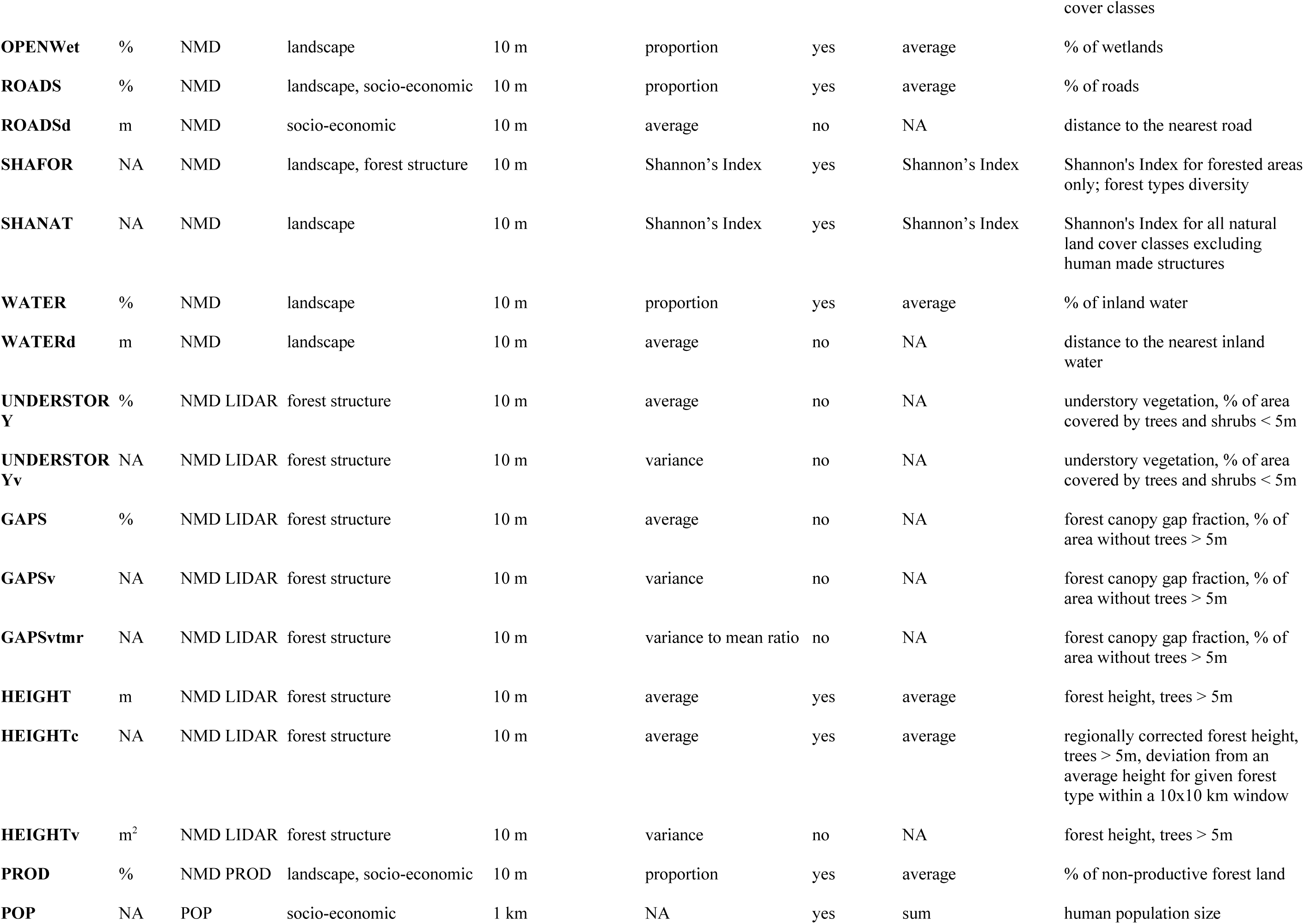
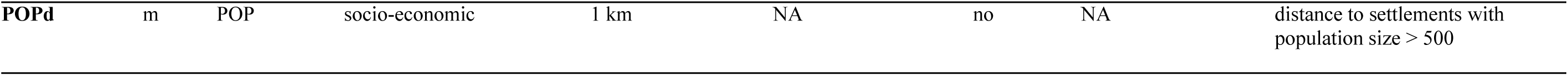
The list of 31 spatial predictors used as explanatory variables in this study together with their basic descriptions, data sources and pre-processing (re-sampling) strategies. The multi-scale variables (value “yes” in the “Multi-scale” column) were coded for further analysis using the following template: {variable_acronym}{spatial_scale_code}; the codes for spatial scales are as follows: 003: 0.3 km, 005: 0.5 km, 011: 1.1 km, 051: 5.1 km and 101: 10.1 km. The “1 ha function” is the function used to re-sample rasters from the original variable’s resolution to a target resolution of 1ha, when applicable. The “Multi-scale function” is the function used to aggregate spatial information within (multi-scale) moving windows of different sizes. In total, the initial list of predictors, including all derived multi-scale variables, contained 128 variables.

**Table 3.** The results of 10-fold spatial cross-validation^53–55^ for each study region (mean ± standard deviation). Next to the classification accuracy and TSS^100^ (True Skill Statistic) metrics (threshold set to 0.5), multiple threshold-independent model performance metrics were evaluated: ROC AUC^97^ (the area under the ROC Curve), PR AUC^97^ (the area under the Precision-Recall Curve), Pearson correlation coefficient, Briers score^98^ and MCC^99^ (Matthews Correlation Coefficient).

| Region | Accuracy | TSS | ROC AUC | PR AUC | Pearson R | Briers score | MCC |
| --- | --- | --- | --- | --- | --- | --- | --- |
| North boreal | 0.81 ± 0.02 | 0.61 ± 0.05 | 0.89 ± 0.01 | 0.89 ± 0.02 | 0.68 ± 0.03 | 0.14 ± 0.01 | 0.62 ± 0.04 |
| South boreal | 0.81 ± 0.02 | 0.61 ± 0.05 | 0.90 ± 0.02 | 0.87 ± 0.04 | 0.69 ± 0.04 | 0.14 ± 0.01 | 0.62 ± 0.05 |
| Hemiboreal | 0.81 ± 0.02 | 0.60 ± 0.06 | 0.89 ± 0.02 | 0.84 ± 0.02 | 0.67 ± 0.03 | 0.14 ± 0.01 | 0.61 ± 0.04 |
| Nemoral | 0.81 ± 0.05 | 0.57 ± 0.16 | 0.89 ± 0.03 | 0.85 ± 0.05 | 0.66 ± 0.06 | 0.15 ± 0.02 | 0.61 ± 0.09 |

The variable selection procedure to decrease the level of co-linearity between the initial 128 explanatory variables, applied before the 10-fold SCV and training the final models, reduced the number of variables to 49, 50, 53 and 48 for the North boreal, South boreal, Hemiboreal and Nemoral regions, respectively (Supplementary Figures 2-5). Interestingly, although a unique combination of variables and spatial scales was selected in each model, we observed the same variables (and similar spatial scales) amongst the most influential ones for each model (Figure 3; Supplementary Figures 2-5; see Table 2 for all variable acronyms). Moreover, the Partial Dependencies Plots (PDPs)^43,56^ clearly indicated consistent patterns of the variables’, often strongly non-linear, relationships with the relative likelihood of HCVF occurrence across all independently trained models. This included 1) the variables describing forest structural properties, such as the uncorrected and regionally-corrected forest height calculated at a target 1 ha scale (HEIGHT, HEIGHTc), as well as at 0.3 (HEIGHTc003), 0.5 (HEIGHT, HEIGHTc005) and 0.11 (HEIGHTc011) spatial scales (expressed in km as the length of a squared window placed at the centre of a target 1 ha pixel), which were some of the most important variables for all regions, and the variation in forest height within each 1 ha pixel (HEIGHTv), a highly important variable for the North boreal and South boreal regions; 2) the variables describing the multi-scale landscape patterns of forest management intensity based on Hansen’s data^39^ (GFC; combined layers of forest loss and gain, HANSEN003, HANSEN005 and HANSEN011) and the Swedish national land cover data (NMD; logged forest, FOPEN003 and FOPEN011), which were the most influential variables for all regions; and 3) the variables describing the physical structure of a landscape (also a proxy for its accessibility) as DEM011 and SLOPE011 (dominant drivers for North boreal, South boreal and Nemoral regions).

**Figure 3.**
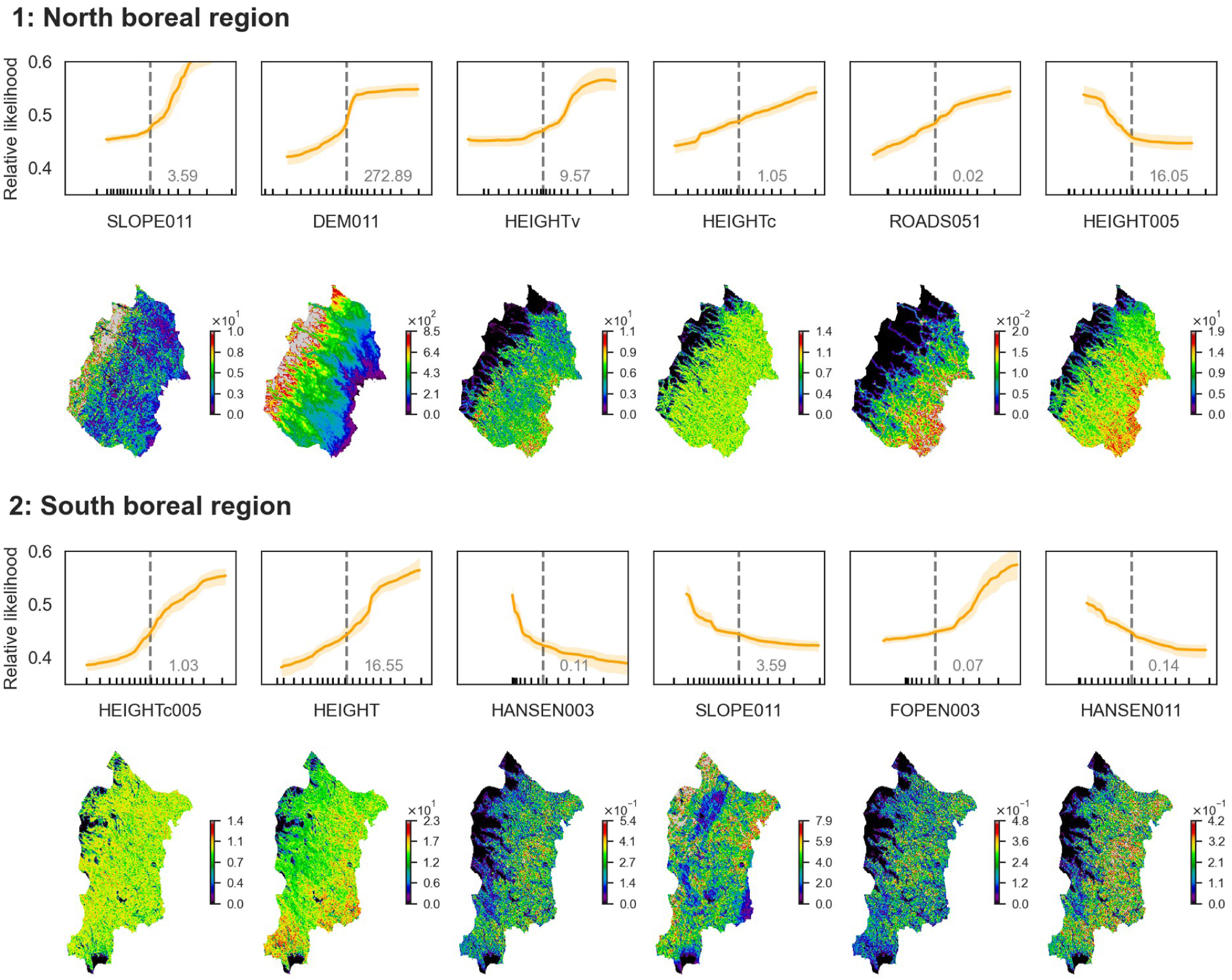
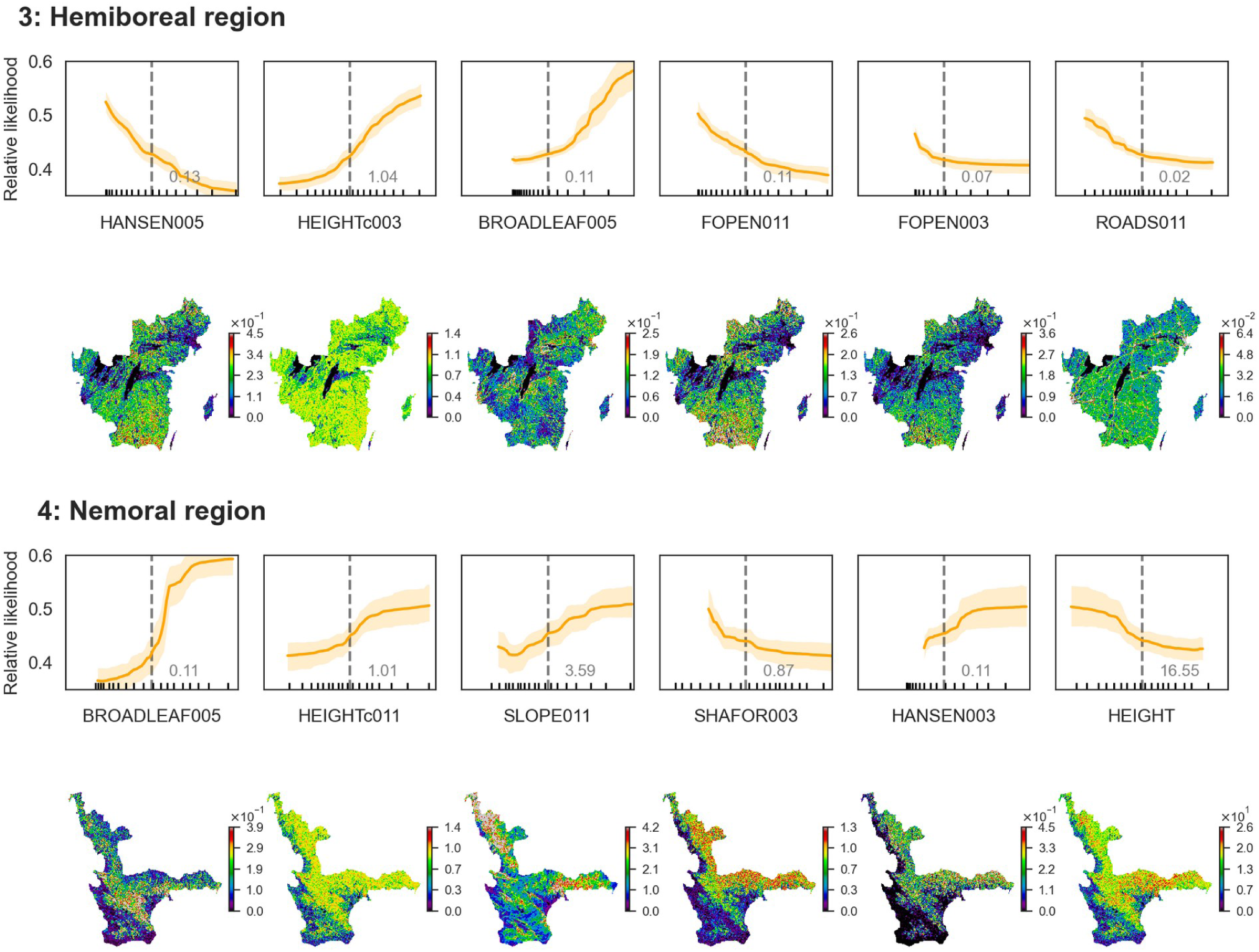
The partial dependence plots (PDPs)^43,56^ for each study region of the six most influential variables in the Random Forest models (top row) and the corresponding maps (bottom row). A partial dependence is the dependence of the relative likelihood of HCVF presence on one predictor variable after averaging out the effects of the other predictor variables in the model. PDPs show the mean response with 95% confidence intervals obtained from 500 bootstrap replicates. All variables were scaled to 0 mean before the model training. The vertical dashed-line, together with the corresponding number in grey, indicates the mean value in the original variable units. The maps are visualized in the original variable units. See Table 2 for the full list and detailed descriptions of all explanatory variables used in this study. The multi-scale variables are coded using the following template: {variable_acronym}{spatial_scale_code}; the codes for spatial scales are as follows: 003: 0.3 km, 005: 0.5 km, 011: 1.1 km, 051: 5.1 km and 101: 10.1 km.

Across all regions, HCVF were more likely to be found in areas with more complex topography (SLOPE, SLOPEv), higher elevation (DEM), in tree stands that are higher than the regional average (HEIGHTc) and structurally more diverse (HEIGHTv). All the above variables are consistent with higher levels of forest naturalness, and showed monotonically increasing non-linear relationships with the relative likelihood of HCVF occurrence, in most cases with a form of sigmoid-shaped function (Figure 3). In the Hemiboreal and Nemoral regions, the proportion of broadleaf tree stands also followed the same pattern. The opposite, i.e. monotonically decreasing relationships, were observed for all variables representing different aspects of a priori negative human impact on landscapes surrounding the target 1 ha forest pixels. These were multi-scale and distance-based variables related to both direct forest management practices and indirect measures of landscape accessibility for forestry (e.g. HANSEN, FOPEN, ROADS). These patterns were consistent among regions.

The multi-scale variables, acting as proxies for forest management intensity (e.g. HANSEN or FOPEN), were important predictors not only in forest dominated landscapes, but also captured isolated HCVF patches surrounded by a non-forest matrix (e.g. forest patches on islands, along the sea coast, or surrounded by agricultural fields). Here, a low value for forest management intensity in the immediately surrounding pixels increased the relative likelihood of HCVF presence.

### Validation of predictions

Two independent sources of relevant spatially explicit information were used to validate the model. The stand-level validation of predicted relative likelihood of HCVF occurrence, based on Sveaskog forest company data (n=57548 tree stand polygons), showed consistent patterns across all four study regions (Figure 4, Supplementary Tables 1-4). The mean predictions per category clearly followed the ordinal scale of forest management types. The highest values were associated with tree stands set-aside for conservation, i.e. without any management NF (mean values for regions 1-4: 0.64, 0.67, 0.59, 0.72) and NF_NM (0.72, 0.70, 0.67, NA), followed by tree stands with conservation-oriented management NM (0.57, 0.53, 0.58, 0.63), and production forests with enhanced conservation concern PE (0.42, 0.40, 0.43, 0.59). The lowest values were associated with general concern production forests PG (0.29, 0.28, 0.32, 0.37). The pairwise numerical differences between conservation-oriented (NF, NF_NM and NM) and production-oriented (PE, PG) management objectives were highly statistically significant (Tukey’s posthoc test, p-values < 0.001) in all regions except the Nemoral region where there was no significant difference between types PE, NM and NF. Similarly, for the forest naturalness variable in all regions, tree stands labelled as natural had significantly higher mean predicted values (0.65, 0.63, 0.58, 0.65) than tree stands without that label (0.30, 0.30, 0.33, 0.39).

**Figure 4.**
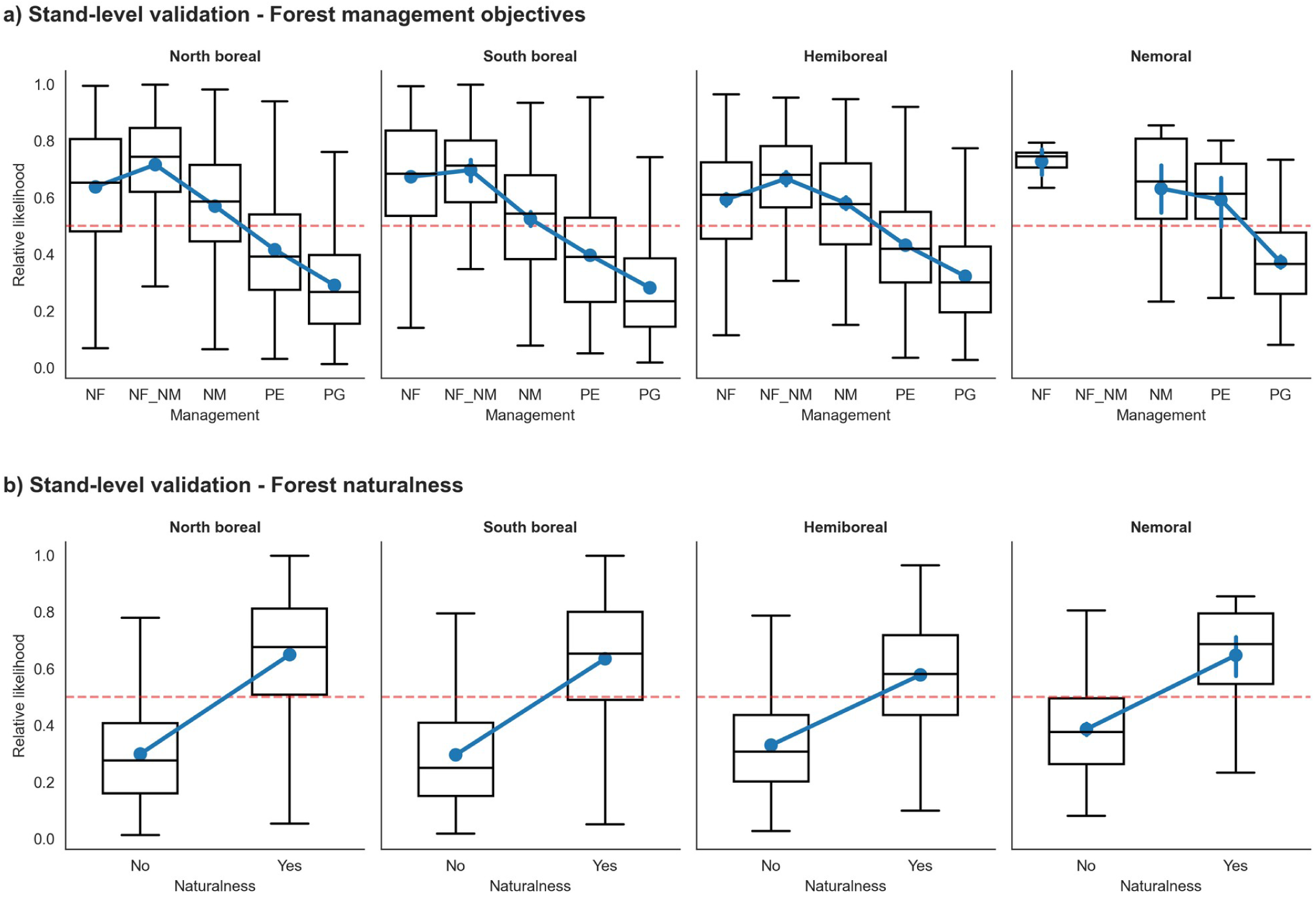
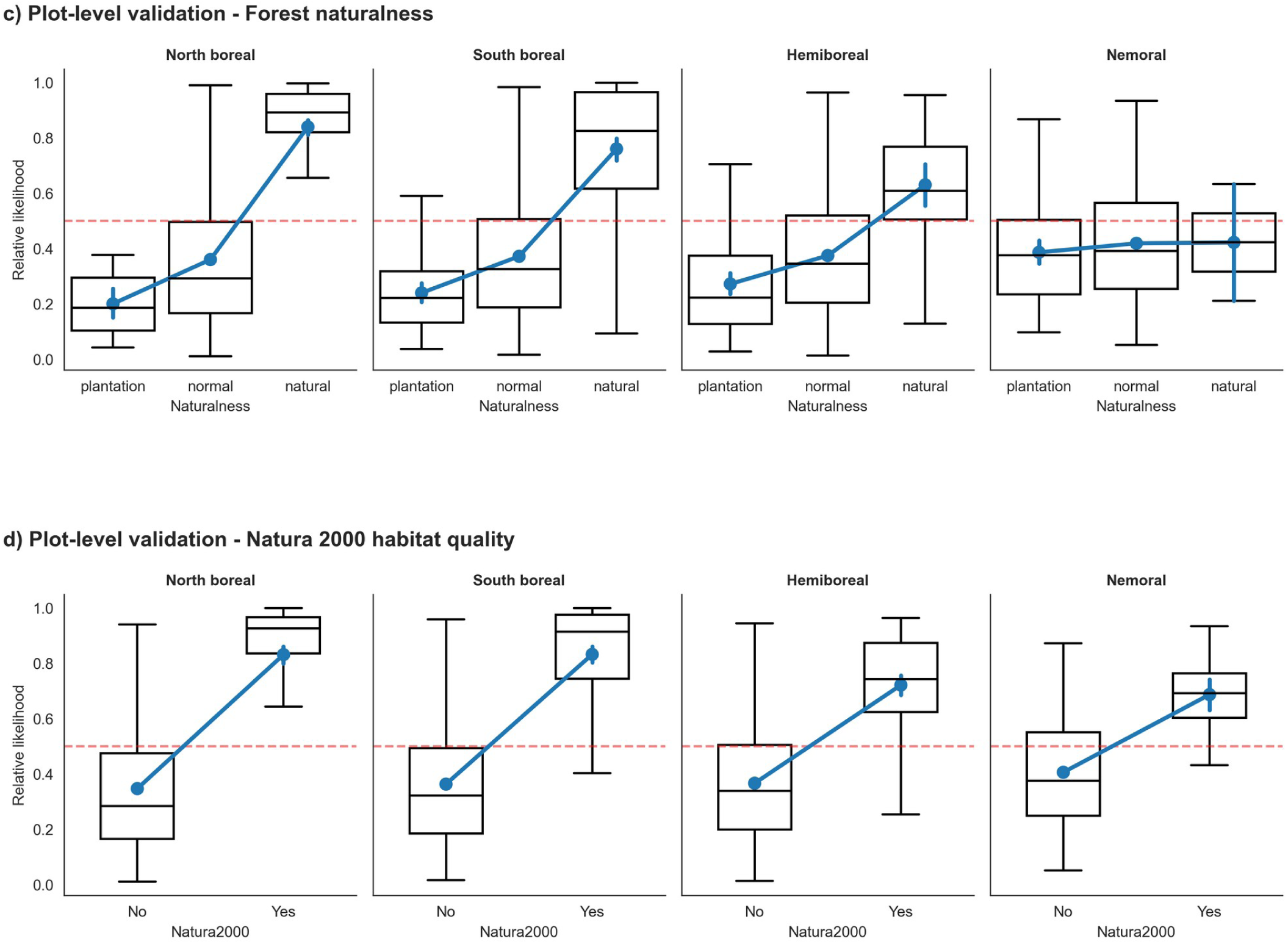
The results of external validation with the Sveaskog forest management compartment dataset (stand-level; n=57548 polygons), and the NFI (plot-level; n=13775 plots) datasets. The boxplots show the median and the quartiles of the dataset, while the whiskers extend to show the rest of the distribution, except for points that are determined to be “outliers” using a method that is a function of the inter-quartile range. Additionally, the boxplots are overlaid with the mean values (blue dots) with 95% confidence intervals. Four categorical variables related to forest naturalness and conservation values verified in the field were used for validation: a) Forest management objectives for individual compartments (Sveaskog): "NF" (Nature conservation, non-intervention), "NF_NM" (Nature conservation, not yet specified), "NM" (Nature conservation-oriented active management), "PF" (Production with enhanced conservation concern), and "PG" (Production with general conservation concern); b) Forest naturalness (Sveaskog, binary); c) Forest naturalness (NFI): “plantation”, “normal”, “natural”; d) Natura 2000 habitat qualities (NFI, binary). The horizontal red dashed-line indicates the HCVF relative likelihood threshold at 0.5.

The results from the second validation, based on Swedish National Forest Inventory (NFI) plot-level data (n=13775 plots) were in accordance with the Sveaskog stand level validation and were also consistent among regions (Figure 4, Supplementary Tables 5-8). In the North boreal, South boreal and Hemiboreal regions, NFI plots that had already been recognized as having a high level of forest naturalness (category “natural”; mean values for regions 1-3: 0.84, 0.76, 0.63) had significantly higher values of predicted relative likelihood of HCVF occurrence than average (category “normal”; 0.36, 0.37, 0.38) and forest plots labelled as “plantation” (0.20, 0.24, 0.27). In the Nemoral region there was no difference between the naturalness categories; however, there were only two data points available for the “natural” category. The comparison of predicted values between NFI plots classified as having Natura 2000 habitat qualities showed clear and highly statistically significant differences in all regions. Those plots had much higher values of predicted HCVF relative likelihood (mean values for regions 1-4: 0.83, 0.83, 0.72, 0.69) compared to the areas without such qualities (0.35, 0.36, 0.37, 0.41).

## Discussion

This study explored integrating machine learning and landscape data mining to scan forest landscapes across all forestland in Sweden with respect to the relative likelihood of hosting biodiversity hotspots in the form of high conservation value forests (HCVF). Applying the Random Forest model generated high-accuracy predictions, resulting in a HCVF relative likelihood surface in the form of a thematic map for ranking forests and landscapes with respect to their levels of naturalness (Figure 5). We validated the models’ results against different independent data representing the naturalness concept at both the forest stand and plot scales. This confirmed that the predicted relative likelihoods of HCVF occurrence actually represent forests with different levels of naturalness and conservation values. We thus demonstrated that publicly available spatial datasets and current machine learning-based predictive modelling can generate the urgently needed mapping of forests with high conservation values, as well as to identify forests with low risks for conflicts between intensive forestry and biodiversity conservation^57^. Our predictions are meant to be used as the first step in the process of making informed strategic conservation decisions and forest management planning. Obviously, before making any final tactical and operational decisions concerning area, field validation should always be made.

**Figure 5.**
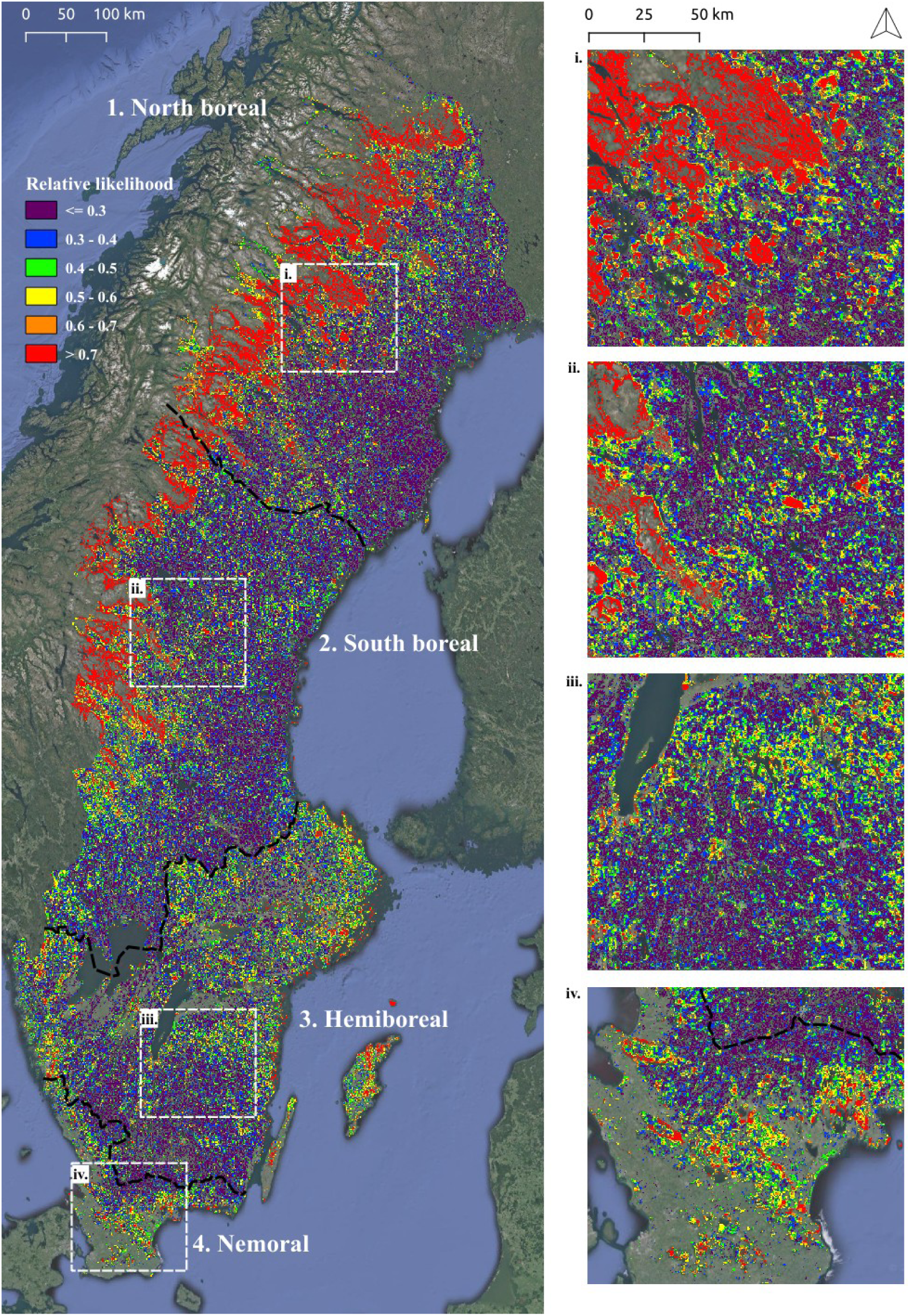
The final prediction map visualizing the relative likelihood of HCVF occurrence for the entire Sweden with 1 ha spatial resolution, and which can be used for ranking landscapes with respect to forest naturalness. An independent Random Forest model was trained for each study region, and predictions for all regions were compiled into the final map. The background map is the Google Satellite layer. The online interactive version of this map is available at: https://bubnicki.users.earthengine.app/view/swedentest.

### Comparison with other studies

Attempts to map forests with high levels of naturalness and conservation value encompass many different approaches ranging from systematic field inventories (e.g. woodland key habitats schemes in northern Europe)^58^, analyses of historical and contemporary databases^29,30,59^, and more recently, remote sensing based analyses using multiple sensors and increasingly advanced mapping methods, including the application of machine learning^3,11,28,60,61^. However, in contrast to our work, most other recent studies using high-resolution and wall-to-wall spatial datasets to map HCVF either apply a global or continental perspective, as is the case with the maps of global Intact Forest Landscapes^3^ and European primary forests^11^, or encompass only local areas of general conservation interest^28,60^. In the case of global or continental models, the spatial and thematic mapping resolution is usually too coarse to be useful for strategic landscape-scale spatial planning. For example, models generating large-scale predictions are trained using limited and spatially clustered data, often predicting (extrapolating) beyond features space covered by reference data^62^. The latter usually results in weak predictive performance, especially when assessed using spatial k-fold cross-validation strategies^55,62^ (but also see^63^). In addition, mapping efforts focused exclusively on target areas of general conservation interest are obviously limited by their spatial coverage.

Consequently, these mapping approaches (i.e. global and targeting conservation areas) are not well suited for strategic landscape-scale spatial planning and prioritizing areas for conservation and restoration. This lack of “actionable” maps has been identified as a serious obstacle for developing national climate and biodiversity strategies and implementing effective actions in conservation and restoration of biodiversity^64^. Our study, presenting detailed information on the relative likelihood of HCVF occurrence for an entire country, demonstrates a clear advantage supporting informed conservation planning, identifying patches of high conservation interest, and the spatial opportunities for expanding or linking such patches through area protection or active forest restoration^13^. It can be inferred, for example, that areas with intermediate likelihood values that are spatially connected to protected areas can support their functionality through passive or active restoration that over time advances their conservation values. Moreover, our modelling approach also identifies landscapes with low level of forest naturalness and conservation value. Two alternative future management trajectories could be considered for such areas; either wood biomass oriented production forestry can be continued, or depending on local environmental priorities, forest conservation and restoration plans can be developed and implemented^13,57,65^.

The complexity of spatial, non-linear and multi-scale interactions between all dimensions describing HCVF, calls for a robust machine learning-based mapping methodology, able to account for the expected spatial variation in the importance of different factors determining the probability of HCVF occurrence in different landscapes. Global models, usually trained using limited and spatially clustered data^62^, tend to over-simplify local-scale interactions trying to generalize over large spatial domains, which often results in poor predictive performance at local (landscape) scales, even when using robust machine learning algorithms such as Random Forest or Boosted Regression Trees^11^. The potential solution would be to apply a spatially-explicit machine learning algorithm able to cope with spatial heterogeneity in explanatory variables and their interactions over large spatial domains, but these methods are currently either at a early stage of development (e.g. Spatial Random Forest)^66,67^ or rarely applied in the ecological and conservation context^68^. A practical workaround we used in this study was the “regionalization” of the study area, i.e. dividing it into four regions characterized by different climatic, anthropogenic and historical forest conditions and the fitting of an independent Random Forest model to each.

The recent studies by Munteanu et al.^28^ in the Romanian Carpathians and by Ørka et al.^61^ in Norway are conceptually and technically the most similar to our work. Munteanu et al. used Maxent software, satellite images and information on current potential anthropogenic pressure to map HCVF in the Carpathians in Romania. However, it is difficult to assess the real predictive power of this model as the training data were highly spatially clustered and the model performance assessment did not take into account spatial correlation in training data that had clearly not been generated by a random probabilistic sampling process^55,62^. Moreover, they excluded historically disturbed forests from their consideration and, contrary to our work, targeted only forests with the highest level of naturalness, thus making the output map less suitable for landscape-scale spatial planning applications. Additionally, while Munteanu et al. proposed an interesting conceptual framework for mapping HCVF, they neither explicitly included the interactions between structural and human-related HCVF dimensions, nor explored the effects of landscape variables at multiple scales as we did in our study.

Ørka et al.^61^ recently presented a framework for a remote sensing-based forest ecological base map of Norway. One of the mapped variables was forest naturalness classified as a binomial variable using generalized boosted regression modelling. Similarly to our work, Ørka et al. stratified the study area into the five regions and fitted an independent model to each region, and provided the maps with predictions for the entirety of Norway, although at a much coarser spatial resolution (10×10 km^2^) than in our study. This makes the results less suitable for landscape-scale spatial planning and conservation prioritization. The most evident difference between our study and Ørka et al’s. forest naturalness model is, however, that they focused on only LiDAR-derived structural metrics as one dimension of HCVF, but not drivers affecting the level of naturalness.

### The limitations of our approach

As training data, we had access to a comprehensive compilation of identified HCVF as included in the national Swedish database^34^ with updates in 2019 and 2020. As such it covers all of Sweden, includes all types of forests, and is representative in terms of its spatial coverage of different landscapes across the entire country. Yet, the database does not formally represent a random probabilistic sample of existing HCVF. Being identified over decades, for different purposes, using different (but comparable) protocols over many years and including formally protected and voluntarily set aside forests as well as non-protected forests, their actual level of naturalness and conservation value may differ. This variability could not be controlled for in the modelling and represents potential noise in the predicted relative likelihood of HCVF occurrence. Moreover, we could not track the cases where a field inventory failed to confirm that a candidate forest is actual HCVF, because this information was not saved in the database. We accounted for this while arranging the training data (presence-only) and interpreting the final results (relative likelihood instead of an actual probability), as described in detail in the methods section. Still, by building our models on more than 6000 a pixels (1 ha) selected out of a total area of c. 3.44 million of ha of confirmed HCVF by applying our sampling strategy (including a minimum distance of 5 km between sampled pixels and filtering out areas < 10 ha), we compiled a representative training dataset of HCVF occurrences covering different landscapes and distributed uniformly across the entire country and the four study regions.

All our models use Global Forest Change (GFC)^39^ data as one of the variables predicting the relative likelihood of HCVF occurrence. Acknowledging issues related to the use of these data in calculating the absolute level of forest loss^69^, we argue that our models use these data in a correct manner. In particular, we used the GFC as one of many potential proxies of human-related pressure on forest areas that, in most cases, correctly reflected the spatial patterns of forest management intensity at the local landscape level. By combining both GFC “loss” and “gain” layers, we provided the model with information about the continuity of forest cover (since 2001) at multiple scales. However, this information indeed does not take into account the origin of the detected change (i.e. natural disturbance, final felling or thinning).

The applicability of our approach elsewhere (in terms of its transferability and scalability) may be limited by the availability of spatially-extensive datasets as used in this study. However, as we observe a rapid technological breakthrough in “sensing” the environment generating an enormous amount of new accumulated data, we expect this will change in the near future. Even today, most of the source datasets (or their counterparts) used in this work as spatial predictors are globally or nationally available (GFC, night-time lights, high-resolution land cover maps, road networks, etc.). On the other hand, the availability of the LiDAR-derived proxies describing forest biophysical structural properties is still limited in many parts of the world. An interesting alternative to the airborne LiDAR for obtaining this information can be, at least to some extent, the use of satellite data and statistical extrapolation of point-based measurements of forest canopy height as proposed by Potapov et al.^40^

However, we argue the most limiting factor to apply our approach in other areas is still the availability of HCVF training data. Potential useful sources of national level reference data on HCVF occurrence can include, for example, databases of forest monitoring systems assessing forest naturalness in the field (as e.g. National Forest Inventories in Sweden, Norway and Finland), retrospective remote sensing analysis of forest cover dynamics^12,28,47^, or citizen-science projects inventorying high conservation value areas^64^, potentially also involving different forest landowner categories. At the international level, initiatives like the “European primary forest database v2.0” seem to be especially promising^59^. Moreover, rapidly growing information on the occurrence of species of conservation interest such as that in the Global Biodiversity Information Facility (GBIF) will increasingly allow for additional validation of identified (potential) HCVF areas.

### Making use of the prediction map

In spite of national and international policies on biodiversity conservation, the favourable conservation status of species populations, habitat network functionality and resilience of forest ecosystems is deteriorating^70^. Consistent with the quantitative conservation targets of the Convention of Biological Diversity (CBD)^71^, and the EU Biodiversity Strategy for 2030^72^ proposed to protect at least 30% of the EU land area, of which a third should be under strict legal protection. In contrast to this, the actual area of legally protected forests in Sweden is 8% if including the mountain areas, and only 4% if not. This calls for the filling of the gaps between the targeted and currently set-aside area shares, both through conservation of existing high conservation value forests^73^, management and restoration of near-natural forest remnants^13^. Our approach to mapping the relative likelihood of hosting high conservation value forests (HCVF) can contribute to addressing both these challenges, providing that field validation will precede actual conservation decisions.

In addition to quantitative conservation targets, establishing functional habitat networks also requires that qualitative targets are satisfied^23,71^. An important principle for assessing and planning functional habitat networks is the acronym BBMJ^74^, which stands for Better, Bigger, More and Joined. This approach is a key principle for ranking local landscapes with respect to where establishing protected areas should be focused, or landscape and nature restoration initiated^23^.

The framework we proposed provides a systematic and consistent first-step mapping, based on which a second-step field validation can be designed for final selection of additional areas that strengthen GI functionality. Accordingly, we provide both a national-scale filter for spatial GI-considerations, which currently does not exist but which is urgently needed, and opportunities for cost efficient and precise field surveys of high conservation value forests. Furthermore, the model provides spatially explicit identification of areas in a continuous gradient from the lowest to the highest relative likelihood of HCVF occurrence. The areas where our models predicted the intermediate values of HCVF relative likelihood (i.e. green, orange and yellow colour codes in Figure 5 and Figure 6) thus represent “crossroad” entities that, depending on governance and management choices, can be put on a trajectory to support GI or not. For supporting GI, nature conservation values can develop naturally over time if the focal forest patch is set aside for natural (free) development, or be enhanced through active restoration management^75^. Based on habitat functionality assessments, restoration management can be oriented towards improving currently unfavourable ecological status by e.g. increasing quantities of dead wood, increasing structural complexity or favouring broadleaf trees^34^. These forest attributes are insufficient in large areas of northern European boreal forests^13,76^. Additionally, the continuous gradient of predicted values of HCVF relative likelihood, provides opportunities at regional scales to enlarge previously known conservation hotspot areas, such as expanding the Scandinavian mountain intact forest landscape^77^ eastwards, or identifying previously unknown clusters of forests that in the future may develop intact forest landscape qualities and improve landscape connectivity over larger scales. Conversely, landscapes representing the other end of the naturalness gradient are suitable as focal areas for biomass production and intensified forest management^57^.

**Figure 6.**
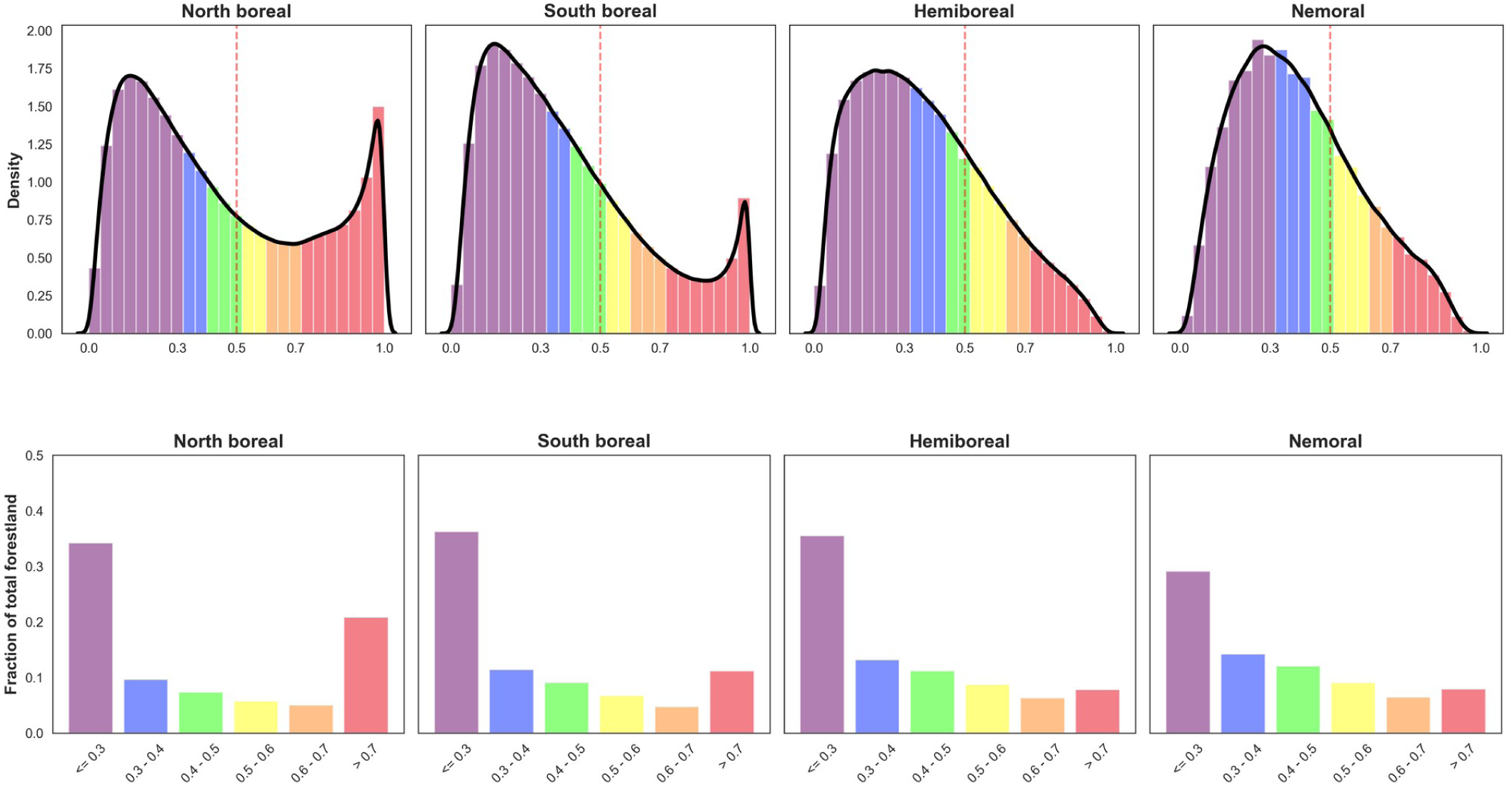
The distribution of predicted relative likelihood of HCVF occurrence (x-axis) for four Swedish regions presented as a) histograms with fitted kernel density estimates (upper row) and b) barplots of the fraction of total forestland estimated per region and including temporary non-forest areas (bottom row). The colours correspond to the six classes of HCVF relative likelihood visualised in the prediction map (Figure 5).

Potentially, our model could serve for designing a landscape-scale zoning of management as proposed by e.g. the TRIAD approach^78^. Hence, despite our focus on the conservation opportunities that are generated by the model and its outputs, we stress that low to intermediate probabilities are indicative for lower conservation ambitions. In combination with projections on climate-change induced site changes and various biophysical data, climate-smart management approaches may be outlined for continued wood biomass oriented forestry as well as on alternative closer to nature management alternatives^79^. In addition to configuring forests into efficient spatial management units, it is also important to consider factors such as proximity to road networks and industry, as well as the forestland ownership status, particularly when concerning non-industrial private and private forest company ownership.

### Conclusions

To conclude, in the era of big data and technological revolution, access to free and open evidence-based knowledge is key to democratic governance towards sustainable forest landscapes. Our approach uses open access spatial data including remote sensing and LiDAR data, open source software tools, and machine learning to assist ranking forest landscapes at different scales with respect to their level of naturalness and conservation values. We provide an “actionable” map with sufficient details for regional to local-landscape spatial planning, which fills a critical gap for implementing national, EU and international biodiversity conservation policy.

## Methods

### Study area

The study area covers the entire Sweden of which 70% is forested (Table 1). We divided Sweden into four regions (Table 1, Figure 1) based on biogeography (from nemoral to north boreal) delimited by county borders, which represent units for statistical reporting and regional spatial planning. Although forest is the main land cover in all regions, the fraction is lowest in the Nemoral (49.2%) and highest in the South boreal region (78.0%). Scots pine (*Pinus sylvestris*) forests dominate (39.8%), followed by Norway spruce (*Picea abies*) forests 27.7%, mixed coniferous forests 12.8% and mixed coniferous and deciduous forests 7.0%^48^. Broadleaf forests cover 7.4% of the entire forestland, which include 1.1 million ha of subalpine mountain birch (*Betula pubescens* ssp. *czerepanovii*) tree line forests^80^.

The Swedish forest landscape has been subject to clearing of forests for agriculture on fertile soils for millennia. Commercial harvesting of wood has occurred since Medieval times^81^, and has re-shaped the forest landscape from the 1800s onwards, with forestry expanding from the south, north-and westwards throughout the country^82^. Initially, logging targeted large diameter trees for saw timber, and regionally even-aged managed forests were exploited to provide wood and bioenergy for the mining and iron industries, which reduced growing stocks significantly up to the early 20^th^ century^83^. In the 1920s, silvicultural measures to increase wood production were introduced, and since then the growing stock has increased significantly. Clearcutting became the dominant harvesting method from the 1950s and onwards. Currently, approximately 70% of the Swedish forestland has been clear-cut at least once^84^. In the 1990s, some measures were introduced to ameliorate the impact of intensive silvicultural practices on biodiversity. Current management practices include soil scarification, planting, pre-commercial and commercial thinning, and in many cases also draining and fertilization^85^. This has created forests with high growth rates, dominated by cohorts of single tree species (mainly Picea abies or Pinus sylvestris) void of old-growth characteristics^48^. Generally, historical land clearing for agriculture has had a larger impact in the Hemiboreal and Nemoral regions than in the North and South boreal, where human expansion and settlement are less pronounced, being limited to fertile soils and favourable local climates particularly along river valleys. Nationally, the largest share of protected forests is in the Scandinavian Mountains Green Belt^12^ of the North and South boreal regions, in which also the presence of the indigenous Sami culture and reindeer husbandry is an important characteristic of forest landscapes.

### Data sources

We used the publicly available national Swedish HCVF database (see Data availability section for more information about all data sources used in this study) with 641 095 polygons delineating c. 3.44 Mha of forests with known high levels of naturalness and conservation values^34^. This data is a comprehensive compilation of ten different data sources describing forests known to have high conservation value identified via field surveys and delineated based on the forest cover of the national topographic terrain (1:50000) and road maps (1:100000). The database documents the conservation value of HCVFs as the level of naturalness assessed in the field, indicated by dead wood in different stages of decay, multi-layered old-growth vegetation structure, and presence of indicator species. This database originally provided the status up to 2016 but was updated in 2019 and 2020 with the new areas in the mountain region having been added. The database includes different categories of long-and short-term formally protected, voluntarily set aside, and unprotected areas.

We used the Swedish National Land Cover data (NMD) as a source of information on forests type, height and productivity, as well as non-forest land cover (e.g. open land, water, agricultural land). NMD is provided in raster format with a spatial resolution of 10×10 m^2^ and was produced based on a combination of data sources including existing digital maps (from 2018), satellite images (from 2015-2018) and airborne LiDAR (from 2009-2018). The tree height and individual tree coverage were derived from NMD auxiliary layers based on airborne LiDAR data.

The other publicly available datasets used in this study were 1) a digital elevation model of Sweden (DEM) with a spatial resolution of 50×50 m^2^, 2) Global Forest Change maps with data on global forest loss (2000 - 2020) and gain (2000 - 2012) with a spatial resolution of 30×30 m^2^ (GFC)^39^, 3) a harmonised global night time light dataset with a spatial resolution of 1×1 km^2^ (1992 - 2018; LIGHTS)^86^ and 4) a database of total human population in Sweden with spatial resolution of 1×1 km^2^ (POP).

### Data pre-processing

All raster and vector datasets (see Table 2 and Table 3) were organized into a country-wide GIS database using the GRASS GIS software v8.2.1^87^. We defined the minimum mapping unit as 1 ha (100×100 m^2^) for our predictive modelling, and re-sampled all imported raster and rasterized vector layers to this target resolution. We used dataset-specific re-sampling procedures that, in most cases, preserved the spatial information from the original dataset’s resolution (see below).

The original 10×10 m^2^ NMD raster was decomposed into multiple target land cover classes (or a combination of classes; see Table 2) using the parallelized version of GRASS GIS “r.resamp.stats” resampling algorithm, which aggregates raster values at coarser resolutions using statistical functions. For each target class its area proportion within 1 ha raster cells was computed. The same resampling algorithm was used to process the other categorical raster layers (e.g. the forest productivity layer). The categorical maps of forest cover loss (2000-2020) and gain (2000-2012) (GFC) were converted into binary maps (loss/gain vs no change) and merged prior to re-sampling.

The distance-based explanatory proxy variables for high forest management intensity (e.g. distance to roads and built-up areas) were calculated using the original resolution and the GRASS GIS algorithm “r.grow.distance” generating raster maps containing distances to the nearest target features and then re-sampled using the average as an aggregation function. A similar re-sampling approach was used to compute the variables originating from continuous rasters (i.e. maps of landscape or forest structure attributes expressed as arrays of float numbers, e.g. elevation, tree height or night-time lights). In the latter cases, in addition to an average, we also calculated variance among pixels aggregated at a coarser resolution (e.g. variation in forest height).

As tree height is determined by numerous factors, including climate (varying along a latitudinal gradient, from temperate to boreal forests), site productivity and elevation, we developed a simple procedure to compute a corrected (or regionalized) version of the original LiDAR-based tree height variable. For each forest type (derived from NMD) we computed the height deviation from an average value calculated within a 10×10 km^2^ moving window (the variable H5c in Table 3).

The other explanatory variables derived from the “decomposed” NMD layer were two Shannon’s indices expressing diversity of 1) different forest types (SHAFOR) and 2) all the land cover classes that we considered natural elements of a landscape (i.e. not containing human-made features; SHANAT). Both indices were calculated at multiple scales.

In the next step, similar to Cushman et al.^88^, we computed multi-scale versions of selected variables (see Table 2, Supplementary Figure 1) to represent information about the neighbourhood of each target 1 ha forest pixel. Hence, we strengthened our model with information about patterns of landscape configuration and composition across multiple spatial scales^89,90^. We used the moving window algorithm (MW) available in the ndimage module from the scipy v1.6.0 Python package for scientific computing^91^. We ran MW with a variable-specific aggregation function (see Table 2) for five different spatial scales (expressed here as the length of a squared window placed at the centre of a target 1 ha pixel): 0.3, 0.5, 1.1, 5.1, and 10.1 km.

Finally, to focus our predictive modelling of HCVF on 1 ha pixels dominated by forest we developed a forest mask by applying a threshold of 0.5 to the proportion of total forest cover. All pixels below this threshold were not taken into account when preparing the training and validation datasets and making the final prediction maps.

### Modelling approach

The RF classifier is an ensemble model (or meta classifier) that fits many decision tree classifiers (individual models) on various sub-samples of a dataset and then combines predictions from all decision trees to improve the predictive accuracy and to control for over-fitting^43^. It uses bagging (bootstrap aggregation) as an ensemble method. The RF has several advantages over other statistical classifiers, including the ability to model complex interactions among predictor variables and its known robustness in generating useful predictions from noisy, non-normal data^43,52,88^. Another useful built-in feature of RF is that, by design, it provides a probability-like unbiased individual estimate that a given sample belongs to a certain class. However, this estimate is just a fraction of decision trees that vote for a certain class and thus it is not a true probability (i.e. there is no strong theoretical foundation for such an interpretation).

The national HCVF database contains information about forest areas verified as HCVF (i.e. true presences) but not about forests classified as non-HCVF (i.e. true absences). Hence, this is a presence-only (or presence-background) data type^92,93^ for which training and validation subsets need to be selected even more carefully than in the other classification tasks so that they are representative and reflect the composition of an actual landscape. Moreover, this type of data influences how a model output can be interpreted. The HCVF detection process could not be explicitly modelled and there was likely spatial sampling bias when identifying new HCVF areas introduced by various regional socio-economic factors and/or land-use history. Thus, we interpreted and ranked the model predictions as a relative likelihood of occurrence, rather than an actual probability of occurrence^93^. We minimized the spatial sampling bias and ensured that landscape sampling was representative with our procedure of generating training and validation datasets as described below. When designing this procedure, our aim was to mimic (to the largest possible extent) a random probabilistic sampling process *sensu* Meyer & Pebesma^62^.

The HCVF training samples (true presences) were generated using both formally protected and unprotected HCVF areas distributed over all of Sweden. First, the HCVF vector layer was rasterized and overlaid with the forest mask based on the original NMD dataset (10×10 m^2^). Next, we re-sampled the HCVF layer to 1 ha resolution (the result was a proportion of HCVF pixels) and applied a threshold of 0.5 to select only those 1 ha forest pixels that are dominated by HCVF. To reduce noise in the training samples we excluded HCVF areas smaller than 10 ha (i.e. areas consisting of less than 10 connected 1 ha pixels). Finally, we used the GRASS GIS “r.random.cells” algorithm to randomly distribute sampling locations over these areas, assuring a 5 km minimum distance between each two selected pixels to minimize the effect of any potential spatial bias that may have been present in the original HCVF database.

To generate pseudo-absences (background data) we applied a 1 km buffer around HCVF areas to lower the chance of unrecognized HCVFs in the nearest neighbourhood being used as pseudo-absence samples for model training, i.e. minimizing the number of false negative predictions (or increasing model sensitivity/recall). We then subtracted the buffered areas from the forest mask raster and we excluded areas smaller than 10 ha. In the last step we used the same algorithm as for the HCVF presence samples to distribute sampling locations over these subtracted areas, keeping a 5 km minimum distance between selected pixels. After this procedure the number of HCVF pseudo-absences was larger than the number of HCVF presences. This made the training dataset moderately imbalanced.

We trained all RF models and computed performance metrics using the Python library scikit-learn v1.2.1^56^ and imbalanced-learn v0.10.1, and following the recommendation of Valavi et al.^94^, used a balanced RF^95^ implementation, which is more robust in dealing with imbalanced datasets. In this implementation, each tree of RF is provided with a balanced bootstrap sample (i.e. sampling with replacement) using a random down-sampling procedure at the level of an individual tree. Initially, we trained and evaluated all models with default values for all hyper-parameters (see the official scikit-learn documentation), except that the number of decision tress (estimators) was set to 500 (default value was 100). Random Forests are known to work reasonably well with default parameters^94^, and an expected potential gain in performance metrics is usually only around 1-2%^96^. Moreover, our final model validation procedure (see the next section) was based on external independent spatial datasets (of different structure and properties than data used for model training), and it is not clear how this could be integrated into the tuning procedure. However, to ensure that default hyper-parameters are robust, we ran the tuning procedure for the best-performing models using Bayesian optimization with Gaussian processes implemented in the Python package scikit-optimize v0.9.0. For all models and regions, the difference in performance of tuned models compared to models ran with default hyper-parameters was less than 0.5% as measured by ROC AUC (see the Supplementary Table 10 and the Supplementary Materials – Jupyter Notebooks). This confirmed that default values were robust for our application.

We assessed the performance of each model (internal validation) using a 10-fold spatial cross-validation (SCV)^53–55^ and a set of complementary model performance metrics. For the SCV we overlaid on the study area a 20×20 km^2^ grid and randomly assigned all grid cells to different spatial subsets (i.e. folds).

Since we were more interested in predicting the continuous relative likelihood of HCVF occurrence rather than a binary categorization of target forest pixels, which reduces the information content compared with using the full range of values^93^, we put more emphasis on threshold-independent metrics evaluating continuous patterns of predicted values, such as the area under ROC curve (ROC AUC)^97^, area under Precision-Recall curve (PR AUC)^97^ and Brier’s score^98^. The ROC AUC can be interpreted as the overall probability that a classifier will predict a higher probability for true positive cases than true negative cases. The PR AUC was computed to better understand the behaviour and performance of our models in predicting the positive class. As both the precision and the recall are focused on the positive class (i.e. HCVF presence), the PR AUC score gives a general evaluation of the model’s performance related to this class. Brier’s score is the mean squared difference between the predicted probability and the actual outcome. Additionally, we computed two correlation-like metrics: the Pearson correlation coefficient and Matthews Correlation Coefficient (MCC)^99^, which is regarded as a robust metric of the quality of binary classification especially for imbalanced datasets. For completeness, we provided two threshold-based metrics (calculated for 0.5 threshold): accuracy and True Skill Statistic (TSS)^100^. The latter is commonly used in the ecological literature.

To decrease the level of co-linearity between explanatory variables used for model training and to enhance the interpretability of the final model, we followed the procedure proposed by McGarigal et al.^89^ and Cushman et al.^88^ with some modifications. We fit univariate RF models for all 125 variables (including multi-scale variables), assessed their performance with the 10- fold SCV and computed the average values of two metrics: ROC AUC and Pearson correlation between predicted and test HCVF datasets (as computed for each k-fold). In the next step we ranked variables based on ROC AUC and iteratively checked all pairs of variables for severe co-linearity (Pearson correlation between variables > 0.7)^101^. Where co-linearity was found, we selected the variable with a higher ROC AUC if a difference was >= 0.01 or with a higher Pearson correlation (threshold also set at 0.01) if the ROC AUC difference was < 0.01. If there was no difference in both metrics we kept the variable with the higher absolute value of ROC AUC.

Finally, to test the robustness of our model’s specification (M1, code “selected”), we compared its performance against seven alternative models trained and validated using the same data: (M2) the “global” model using data from the entire Sweden (i.e. without the regional stratification; (M3) the “full” model trained with all spatial predictors, including all derived multi-scale features, and ignoring a strong (> 0.7) pairwise correlation between some variables; (M4) the “onescale” model trained with all spatial predictors but excluding all multi-scale features; (M5) the “baseline” model trained using only 4 key spatial predictors, all hypothetically having a strong effect on the probability of HCVF occurrence (elevation, tree height, distance to roads and % of logged areas within 1ha); Additionally, the “full”, “onescale” and “baseline” models were trained with and without longitude and latitude as the auxiliary variables. The results of 10-fold Spatial Cross-Validation (SCV) confirmed the robustness of our model’s specification as it performed best or equally good as the “full” model for all regions. (Supplementary Table 9). Moreover, we have not noticed any significant differences when comparing the results of the external validation (as described in the next section; Supplementary Materials – Jupyter Notebooks). The “global” model performed equally good but, based on pixel-to-pixel pairwise Pearson correlation coefficients, produced different predictions for the Hemiboreal and Nemoral regions when compared with the best “regionalized” models (Supplementary Figure 7), suggesting that global predictions where mainly driven by the patterns that model learnt for the northern Sweden. As expected, the weakest performance was noticed for the “baseline” model followed by the “onescale” model, but interestingly the latter performed almost as good as the best models in all regions.

### Model validation with external data

Two independent sources of relevant spatially explicit information were used for model validation. The first is the forest management data from Sweden’s largest forest owner – the state company Sveaskog. Their forest management data provides information at tree stand (compartment) level. It is based on field sampling that assesses variables such as total wood volume (living trees and deadwood), proportion of wood volume of living trees divided into different species, stand age, site type and many other characteristics. We used two categorical variables from the Sveaskog dataset expressing different levels of forest conservation and naturalness values on an ordinal scale: 1) forest management type (5 levels) and 2) forest naturalness (binary). The forest management type is set for each tree stand using the following categories^102^: 1) NF = Nature conservation, non-intervention; 2) NF_NM = Nature conservation, not yet specified; 3) NM = Nature conservation-oriented active management, often implying restoration measures; 4) PE = Production with enhanced conservation concern; 5) PG = Production with general conservation concern (green-tree and deadwood retention for biodiversity at harvest). For each stand polygon that had a minimum 10 pixels we calculated the mean value of predicted HCVF relative likelihoods. In total, we used data from 57548 Sveaskog tree stand polygons.

The second source of information for model validation was the Swedish National Forest Inventory (NFI), which uses a randomly planned regular sampling grid^103^, including around 4 500 permanent tracts with each tract being surveyed once every 5 years. The tracts have a rectangular shape of different sizes in different parts of the country and consist of 4-8 circular sample plots (each plot 314 m^2^). We used two NFI categorical variables relevant to conservation and forest naturalness values: 1) the level of forest naturalness (three levels: natural, normal, plantation) and 2) a binary indicator if plots meet the minimum requirements to be considered Natura 2000 habitat according to the EU Habitats Directive based on its interpretation in Sweden. NFI data used in this study originated from the 2015 to 2019 inventories. For each NFI plot selected for validation we extracted the average predicted HCVF relative likelihood from the pixel spatially overlapping with the plot and its four nearest neighbours. In total, we used data from 13775 NFI plots spread across all study regions.

We summarized both datasets for each study region using boxplots produced with the Python package Seaborn v0.11.1. Finally, we used Tukey’s posthoc test to check for the statistical significance of the differences between different levels of all validation variables.

### Final predictions

After the 10-fold SCV validation procedure, to achieve the highest prediction performance (see^53^), we produced the final predictions using all available training data to fit new models for all four regions. Here we assume that the model performance metrics estimated from the 10-fold SCV are conservative, i.e. the final models can still perform better^53^. These final predictions were used for model validation with external independent spatial datasets as described above.

To understand which variables were the main drivers of the predicted values for each model, we estimated the impurity-based variable importance^43,56^. Furthermore, to visualise and explore the relationships between the top six most important variables and predict the relative likelihood of HCVF occurrence, we generated plots of partial dependence, which is the dependence of the relative likelihood of HCVF presence on one predictor variable after averaging out the effects of the other predictor variables in the model^43,56^. The 95% confidence intervals of the mean response were obtained from 500 bootstrap replicates.

All resulting maps were created using QGIS v3.22 “Białowieża” (QGIS Development Team, 2022) and Python package matplotlib v3.3.3. The interactive visualisation of the prediction map was implemented using Google Earth Engine platform and is available online at https://bubnicki.users.earthengine.app/view/swedentest.

## Data availability

The spatial datasets used in this study are publicly available:

DEM. Terrain Model Download, grid 50+. Lantmateriet, Swedish Ministry of Finance. Available online at https://www.lantmateriet.se/en/maps-and-geographic-information/geodataprodukter/ produktlista/terrain-model-download-grid-50/ (accessed April 28, 2022).

GFC. Global Forest Change. Global Land Analysis and Discovery, Department of Geographical Sciences, University of Maryland. Available online at https://glad.earthengine.app (accessed April 28, 2022).

HCVF. A database of High Conservation Value Forests in Sweden. Swedish Environmental Protection Agency. Available online at https://geodata.naturvardsverket.se/nedladdning/land/skogliga_vardekarnor_2016.zip (accessed April 28, 2022). [Please note that this database originally provides the status up to 2016 but has been updated in 2019 and 2020 with the new areas in the mountain region which were not publicly available at the moment of writing this manuscript but are available on request from the Swedish Environmental Protection Agency.]

LIGHTS. A harmonized global nighttime light dataset 1992–2018. Available online at https://doi.org/10.6084/m9.figshare.9828827.v2 (accessed April 28, 2022).

NMD. National Land Cover Data. Swedish Environmental Protection Agency. Available online at https://www.naturvardsverket.se/en/services-and-permits/maps-and-map-services/national-land-cover-database/ (accessed April 28, 2022).

POP. Total Population in Sweden. Statistics Sweden. Available online at https://www.scb.se/en/services/open-data-api/open-geodata/grid-statistics/ (accessed April 28, 2022).

The independent spatial datasets used for validation (Sveaskog and NFI) are available upon request from the corresponding author.

## Code availability

The source code to re-produce the spatial data processing pipeline, training and validation of the Random Forest models, and all other analyses presented in this paper is available at https://gitlab.com/oscf/hcvf-model-sweden.

## Supporting information

Jupyter Notebooks

## Acknowledgements

We thank Prof. Carsten Dormann for his valuable feedback and critical comments on an earlier version of the manuscript. We thank X anonymous reviewers for their important feedback during the review process. We thank Tom Diserens for his great help with improving the English language in this manuscript. This study was funded by the Swedish Environmental Protection Agency, grant NV-03728–17, to Bengt Gunnar Jonsson.

## Authors contribution

B.G.J., P.A., J.S., G.M. and J.W.B. originally conceived the idea. J.W.B. developed the methodological approach with help from G.M., B.G.J., P.A., and J.S. J.W.B. collected spatial data with help from J.S., G.M. and B.G.J. J.W.B. processed the spatial data, developed the computer code, trained and validated the machine learning models and produced all figures and maps. J.W.B. led the writing with substantial help from P.A., B.G.J., J.S. and G.M.

