## Supplementary Information for "Mapping forests with different levels of naturalness using machine learning and landscape data mining"

### Supplementary Figure 1

The examples of multi-scale variables derived from a) Global Forest Change maps providing data on global forest loss (2000 - 2020) and gain (2000 - 2012) with spatial resolution of 30x30 m2 (GFC); “HANComb” is a compilation of forest loss and gain layers b) Swedish national land cover dataset (NMD); “SHANAT” is the Shannon's Index calculated for all natural land cover classes excluding human made structures. To generate multi-scale variables we run moving window algorithm with a variable-specific aggregation function (see Table 2) for five different spatial scales (expressed here as a length of a squared window placed at the centre of a target 1ha pixel): 0.3, 0.5, 1.1, 5.1, and 10.1 km.

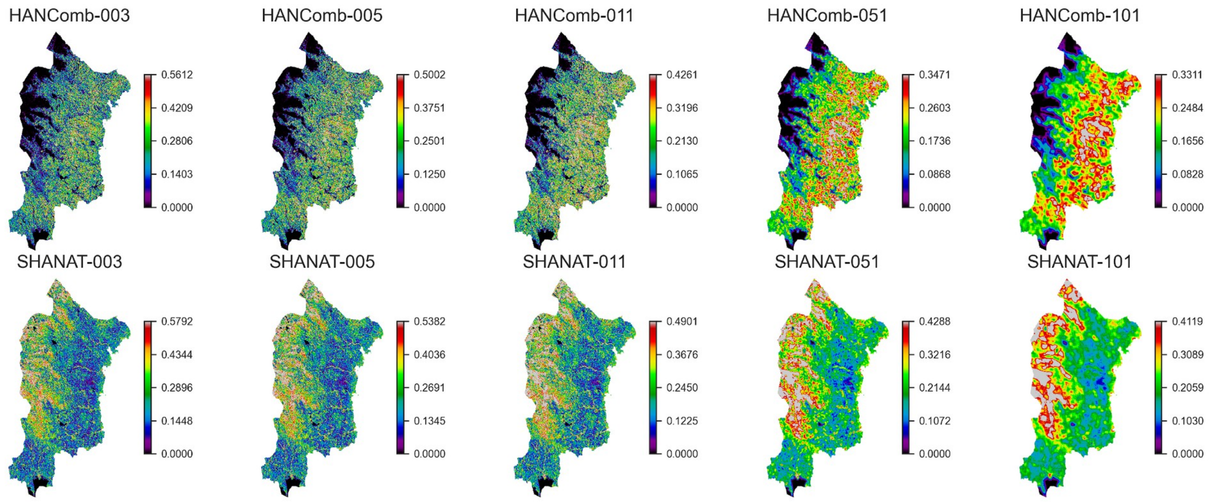

### Supplementary Figure 2

The barplot of the impurity-based variables importance estimated from the final Random Forest model for the North boreal region (see Table 2 for all variable acronyms).

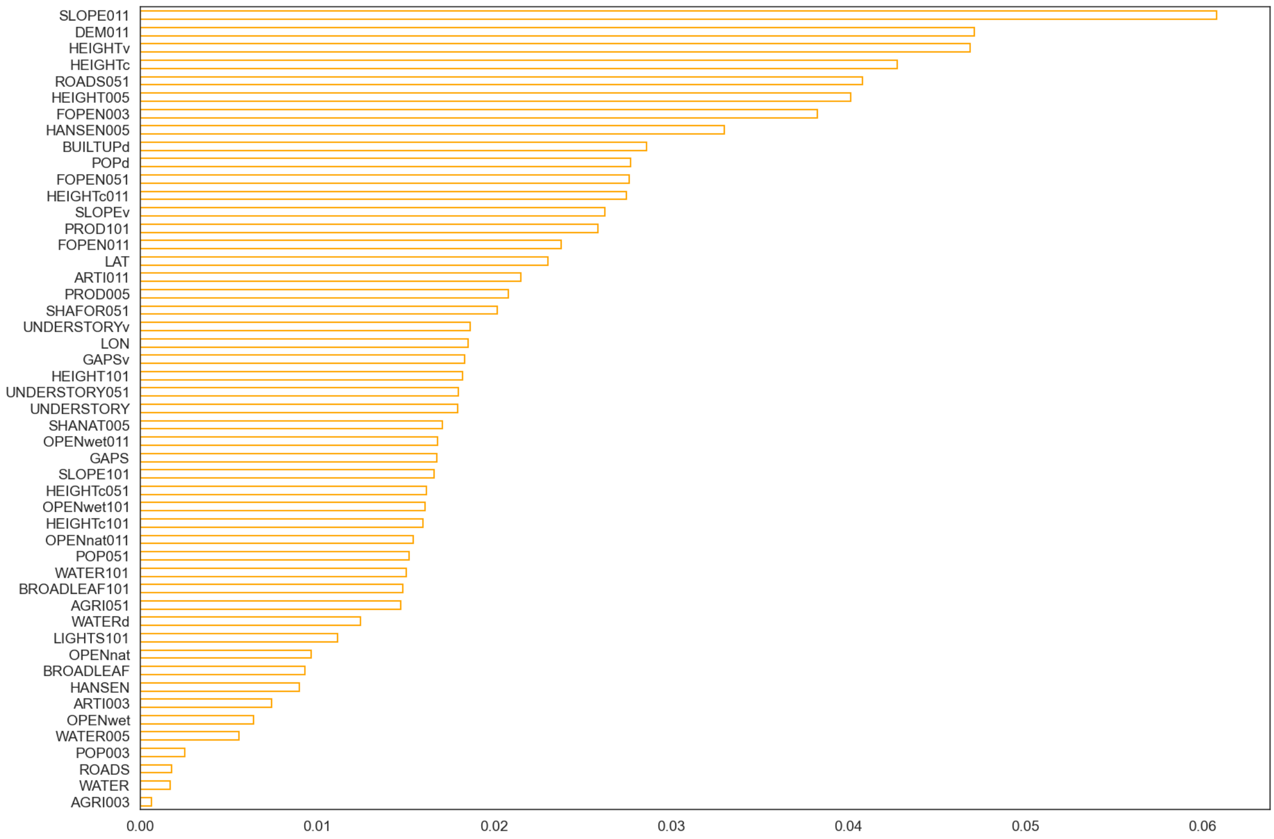

#### Supplementary Figure 3

The barplot of the impurity-based variables importance estimated from the final Random Forest model for the South boreal region (see Table 2 for all variable acronyms).

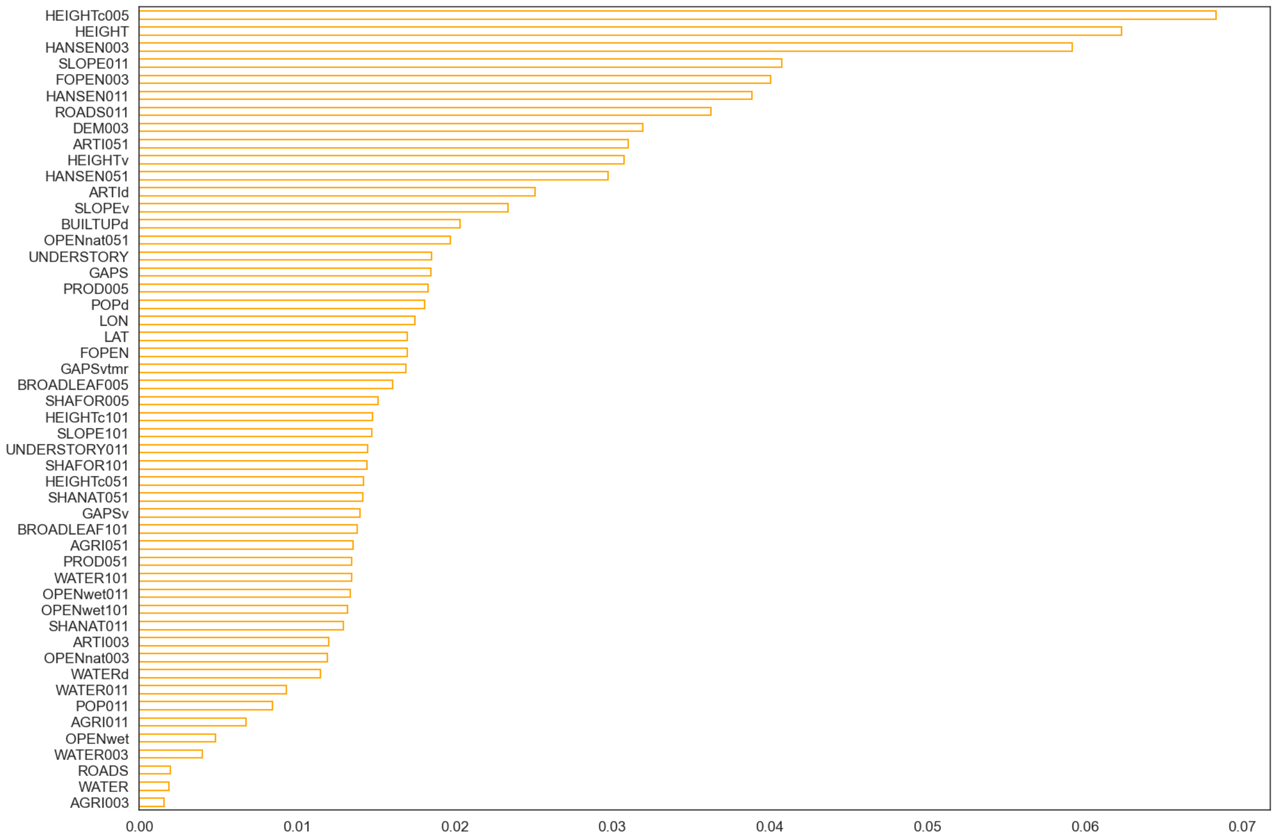

#### Supplementary Figure 4

The barplot of the impurity-based variables importance estimated from the final Random Forest model for the Hemiboreal region (see Table 2 for all variable acronyms).

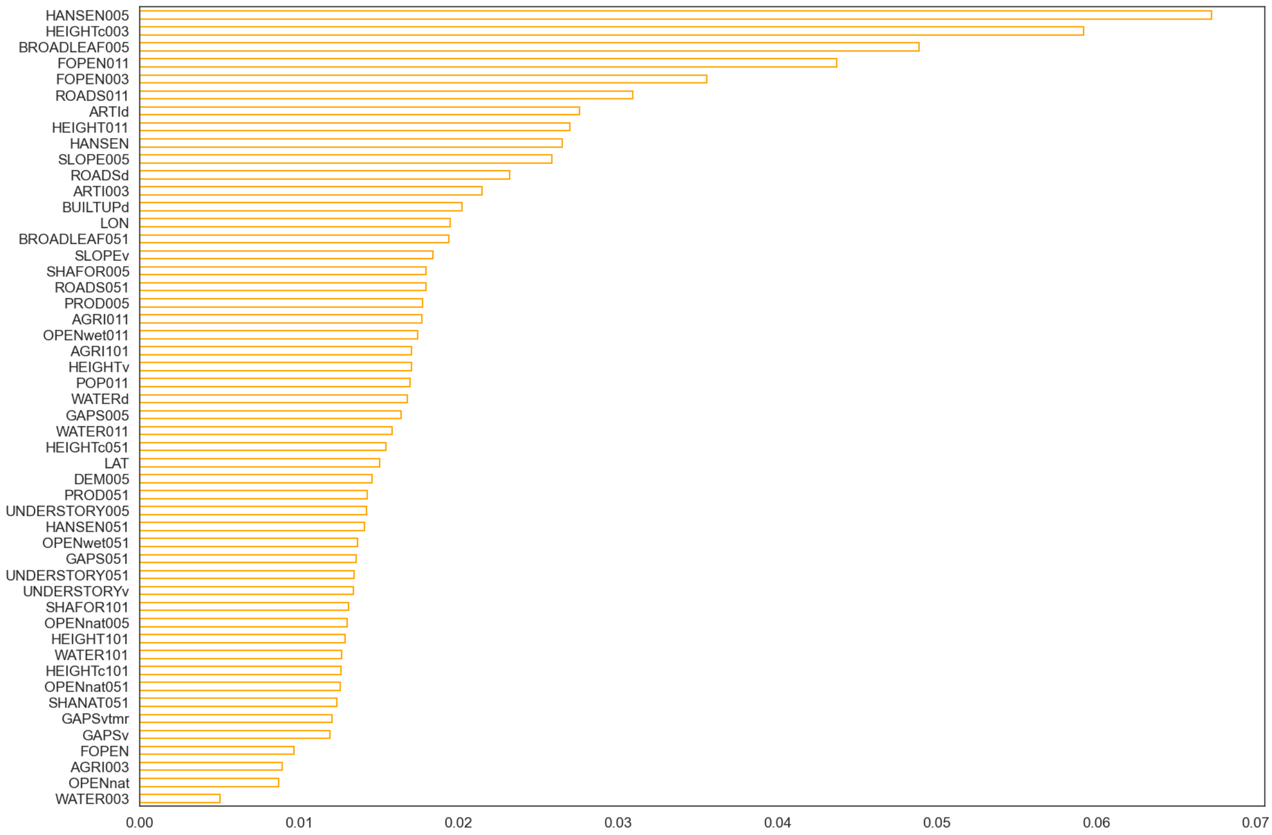

### Supplementary Figure 5

The barplot of the impurity-based variables importance estimated from the final Random Forest model for the Nemoral region (see Table 2 for all variable acronyms).

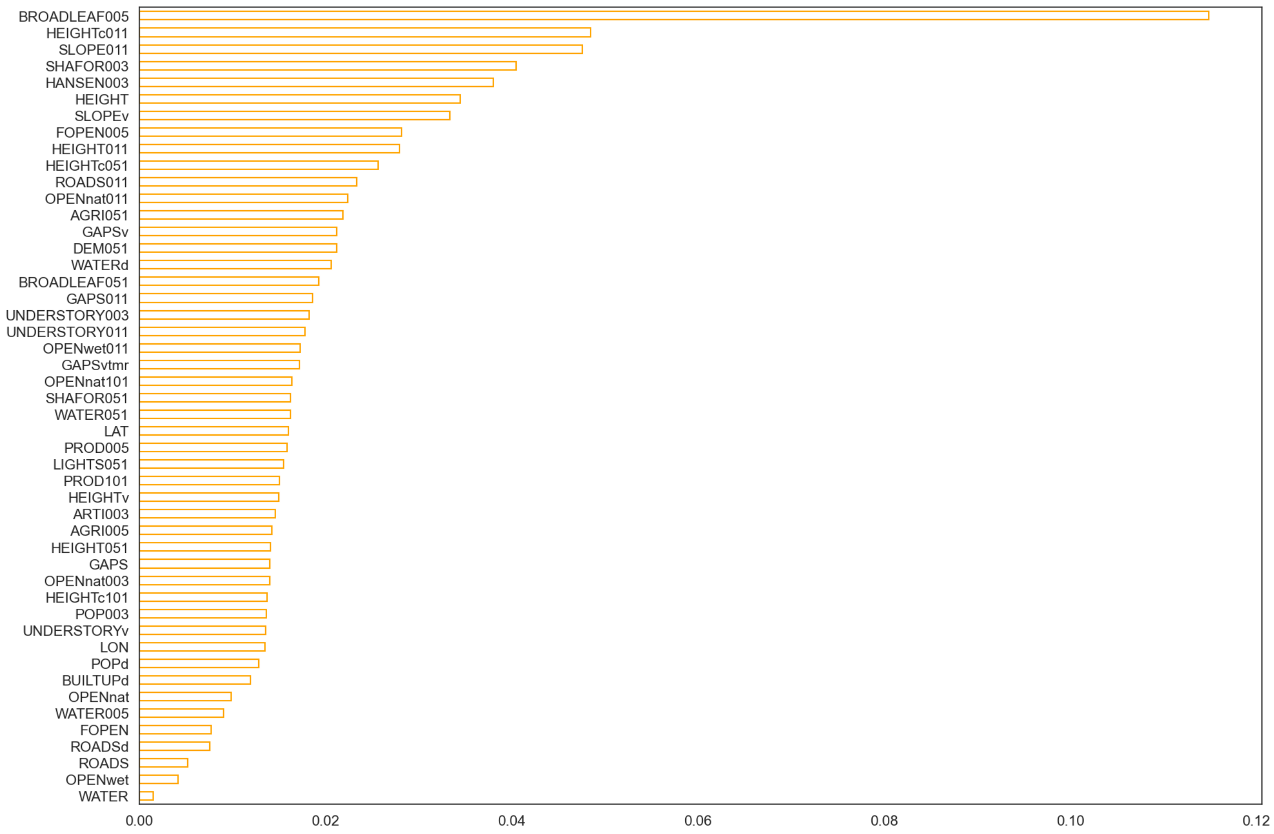

#### Supplementary Figure 6

The comparison of the relative likelihood of HCVF occurrence as predicted by our models with the map of proxy continuity forests (pCF) for the boreal regions. The pCF dataset represents a publicly available, high-resolution (10x10 m2), complete and consistent mapping of forests that have not been clear at least cut since 1950s. The mapping was performed as an automatic retrospective change detection analysis of satellite images from 1973 to 2016 and aerial photos from the 1950s and 1960s by Metria AB on commission by the Swedish Environmental Protection Agency for the entire boreal biome in Sweden. Both maps show the sum of pixel values in 1x1 $km^2$ moving window (MW); the pCF binary raster was re-sampled from 10x10 $m^2$ to 100x100 $m^2$ resolution prior to MW analysis. Please note that these two maps are not directly comparable as they describe different forest properties and are also different numerically (data type, range of values etc), however, the spatial patterns visualised on both maps are very similar, with pixel-to-pixel Pearson's correlation equals to 0.78.

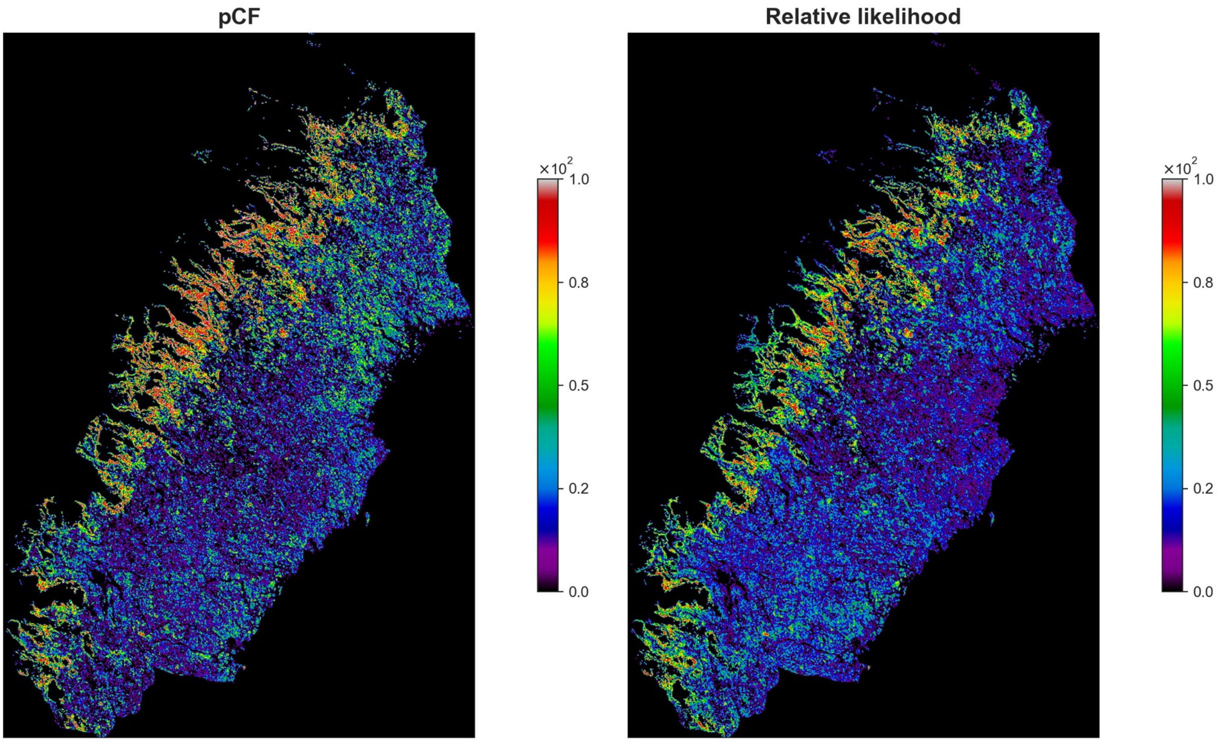

#### Supplementary Figure 7

The heatmaps of pixel-to-pixel pairwise Pearson correlation coefficients for predictions by the eight alternative model's specifications. The comparison of predictions is presented per region and globally.

**North boreal**

| | (M1) selected | (M2) global | (M3a) full | (M3b) full_nolatlon | (M4a) onescale | (M4b) onescale_nolatlon | (M5a) baseline | (M5b) baseline_nolatlon |
|---|---|---|---|---|---|---|---|---|
| (M1) selected | 1 | 0.97 | 0.98 | 0.98 | 0.94 | 0.93 | 0.85 | 0.77 |
| (M2) global | 0.97 | 1 | 0.98 | 0.98 | 0.93 | 0.92 | 0.87 | 0.8 |
| (M3a) full | 0.98 | 0.98 | 1 | 1 | 0.94 | 0.93 | 0.86 | 0.78 |
| (M3b) full_nolatlon | 0.98 | 0.98 | 1 | 1 | 0.94 | 0.93 | 0.86 | 0.78 |
| (M4a) onescale | 0.94 | 0.93 | 0.94 | 0.94 | 1 | 0.99 | 0.86 | 0.8 |
| (M4b) onescale_nolatlon | 0.93 | 0.92 | 0.93 | 0.93 | 0.99 | 1 | 0.84 | 0.79 |
| (M5a) baseline | 0.85 | 0.87 | 0.86 | 0.86 | 0.86 | 0.84 | 1 | 0.89 |
| (M5b) baseline_nolatlon | 0.77 | 0.8 | 0.78 | 0.78 | 0.8 | 0.79 | 0.89 | 1 |

**South boreal**

| | (M1) selected | (M2) global | (M3a) full | (M3b) full_nolatlon | (M4a) onescale | (M4b) onescale_nolatlon | (M5a) baseline | (M5b) baseline_nolatlon |
|---|---|---|---|---|---|---|---|---|
| (M1) selected | 1 | 0.96 | 0.98 | 0.98 | 0.91 | 0.9 | 0.81 | 0.76 |
| (M2) global | 0.96 | 1 | 0.98 | 0.98 | 0.9 | 0.9 | 0.81 | 0.77 |
| (M3a) full | 0.98 | 0.98 | 1 | 0.99 | 0.91 | 0.9 | 0.83 | 0.78 |
| (M3b) full_nolatlon | 0.98 | 0.98 | 0.99 | 1 | 0.91 | 0.9 | 0.83 | 0.78 |
| (M4a) onescale | 0.91 | 0.9 | 0.91 | 0.91 | 1 | 0.99 | 0.86 | 0.81 |
| (M4b) onescale_nolatlon | 0.9 | 0.9 | 0.9 | 0.9 | 0.99 | 1 | 0.84 | 0.81 |
| (M5a) baseline | 0.81 | 0.81 | 0.83 | 0.83 | 0.86 | 0.84 | 1 | 0.91 |
| (M5b) baseline_nolatlon | 0.76 | 0.77 | 0.78 | 0.78 | 0.81 | 0.81 | 0.91 | 1 |

**Hemiboreal**

| | (M1) selected | (M2) global | (M3a) full | (M3b) full_nolatlon | (M4a) onescale | (M4b) onescale_nolatlon | (M5a) baseline | (M5b) baseline_nolatlon |
|---|---|---|---|---|---|---|---|---|
| (M1) selected | 1 | 0.93 | 0.97 | 0.97 | 0.81 | 0.8 | 0.66 | 0.59 |
| (M2) global | 0.93 | 1 | 0.96 | 0.96 | 0.81 | 0.81 | 0.66 | 0.6 |
| (M3a) full | 0.97 | 0.96 | 1 | 0.99 | 0.81 | 0.81 | 0.66 | 0.59 |
| (M3b) full_nolatlon | 0.97 | 0.96 | 0.99 | 1 | 0.81 | 0.8 | 0.65 | 0.59 |
| (M4a) onescale | 0.81 | 0.81 | 0.81 | 0.81 | 1 | 0.98 | 0.77 | 0.7 |
| (M4b) onescale_nolatlon | 0.8 | 0.81 | 0.81 | 0.8 | 0.98 | 1 | 0.74 | 0.7 |
| (M5a) baseline | 0.66 | 0.66 | 0.66 | 0.65 | 0.77 | 0.74 | 1 | 0.88 |
| (M5b) baseline_nolatlon | 0.59 | 0.6 | 0.59 | 0.59 | 0.7 | 0.7 | 0.88 | 1 |

**Nemoral**

| | (M1) selected | (M2) global | (M3a) full | (M3b) full_nolatlon | (M4a) onescale | (M4b) onescale_nolatlon | (M5a) baseline | (M5b) baseline_nolatlon |
|---|---|---|---|---|---|---|---|---|
| (M1) selected | 1 | 0.91 | 0.94 | 0.95 | 0.74 | 0.73 | 0.49 | 0.45 |
| (M2) global | 0.91 | 1 | 0.91 | 0.92 | 0.76 | 0.76 | 0.55 | 0.54 |
| (M3a) full | 0.94 | 0.91 | 1 | 0.99 | 0.79 | 0.78 | 0.48 | 0.45 |
| (M3b) full_nolatlon | 0.95 | 0.92 | 0.99 | 1 | 0.79 | 0.79 | 0.49 | 0.46 |
| (M4a) onescale | 0.74 | 0.76 | 0.79 | 0.79 | 1 | 0.99 | 0.63 | 0.57 |
| (M4b) onescale_nolatlon | 0.73 | 0.76 | 0.78 | 0.79 | 0.99 | 1 | 0.62 | 0.59 |
| (M5a) baseline | 0.49 | 0.55 | 0.48 | 0.49 | 0.63 | 0.62 | 1 | 0.87 |
| (M5b) baseline_nolatlon | 0.45 | 0.54 | 0.45 | 0.46 | 0.57 | 0.59 | 0.87 | 1 |

**Global**

| | (M1) selected | (M2) global | (M3a) full | (M3b) full_nolatlon | (M4a) onescale | (M4b) onescale_nolatlon | (M5a) baseline | (M5b) baseline_nolatlon |
|---|---|---|---|---|---|---|---|---|
| (M1) selected | 1 | 0.95 | 0.98 | 0.98 | 0.9 | 0.89 | 0.79 | 0.72 |
| (M2) global | 0.95 | 1 | 0.97 | 0.97 | 0.88 | 0.88 | 0.79 | 0.74 |
| (M3a) full | 0.98 | 0.97 | 1 | 0.99 | 0.9 | 0.9 | 0.8 | 0.74 |
| (M3b) full_nolatlon | 0.98 | 0.97 | 0.99 | 1 | 0.9 | 0.9 | 0.8 | 0.74 |
| (M4a) onescale | 0.9 | 0.88 | 0.9 | 0.9 | 1 | 0.99 | 0.84 | 0.78 |
| (M4b) onescale_nolatlon | 0.89 | 0.88 | 0.9 | 0.9 | 0.99 | 1 | 0.82 | 0.78 |
| (M5a) baseline | 0.79 | 0.79 | 0.8 | 0.8 | 0.84 | 0.82 | 1 | 0.9 |
| (M5b) baseline_nolatlon | 0.72 | 0.74 | 0.74 | 0.74 | 0.78 | 0.78 | 0.9 | 1 |

#### Supplementary Table 1

The results of external validation with the Sveaskog forest management compartment dataset (stand-level; n=57548 polygons). The table presents the basic statistics of predicted values of relative likelihood of HCVF occurrence for stand polygons belonging to different "Forest management objectives" categories: "NF" (Nature conservation, non-intervention), "NF_NM" (Nature conservation, not yet specified), "NM" (Nature conservation-oriented active management), "PF" (Production with enhanced conservation concern), and "PG" (Production with general conservation concern).

| Region | Management | Mean | Count | STD | CI95_low | CI95_high |
|---|---|---|---|---|---|---|
| North boreal | NF | 0.637 | 4148 | 0.203 | 0.631 | 0.644 |
| North boreal | NF_NM | 0.716 | 2401 | 0.164 | 0.709 | 0.723 |
| North boreal | NM | 0.570 | 903 | 0.195 | 0.557 | 0.582 |
| North boreal | PE | 0.416 | 2516 | 0.189 | 0.409 | 0.424 |
| North boreal | PG | 0.291 | 31816 | 0.168 | 0.289 | 0.293 |
| South boreal | NF | 0.673 | 1245 | 0.192 | 0.663 | 0.684 |
| South boreal | NF_NM | 0.697 | 82 | 0.172 | 0.660 | 0.734 |
| South boreal | NM | 0.525 | 253 | 0.205 | 0.500 | 0.550 |
| South boreal | PE | 0.396 | 868 | 0.200 | 0.383 | 0.410 |
| South boreal | PG | 0.282 | 5541 | 0.175 | 0.277 | 0.286 |
| Hemiboreal | NF | 0.593 | 297 | 0.184 | 0.573 | 0.614 |
| Hemiboreal | NF_NM | 0.667 | 172 | 0.160 | 0.643 | 0.691 |
| Hemiboreal | NM | 0.580 | 303 | 0.185 | 0.559 | 0.601 |
| Hemiboreal | PE | 0.432 | 608 | 0.181 | 0.418 | 0.446 |
| Hemiboreal | PG | 0.322 | 6139 | 0.162 | 0.318 | 0.326 |
| Nemoral | NF | 0.727 | 5 | 0.061 | 0.674 | 0.781 |
| Nemoral | NF_NM | 0.000 | 0 | 0.000 | 0.000 | 0.000 |
| Nemoral | NM | 0.632 | 19 | 0.183 | 0.549 | 0.714 |
| Nemoral | PE | 0.592 | 13 | 0.165 | 0.502 | 0.682 |
| Nemoral | PG | 0.372 | 219 | 0.149 | 0.353 | 0.392 |

#### Supplementary Table 2

The results of external validation with the Sveaskog forest management compartment dataset (stand-level; n=57548 polygons). The table summarizes the Tukey's posthoc tests to check for statistical significance of differences between predicted values of relative likelihood of HCVF occurrence for stand polygons belonging to different "Forest management objectives" categories: "NF" (Nature conservation, non-intervention), "NF_NM" (Nature conservation, not yet specified), "NM" (Nature conservation-oriented active management), "PF" (Production with enhanced conservation concern), and "PG" (Production with general conservation concern).

**a) North boreal**

| group1 | group2 | Diff | Lower | Upper | q-value | p-value |
|---|---|---|---|---|---|---|
| NF | NF_NM | 0.078 | 0.066 | 0.090 | 24.517 | 0.001 |
| NF | PG | 0.348 | 0.340 | 0.356 | 169.699 | 0.001 |
| NF | PE | 0.202 | 0.189 | 0.216 | 58.861 | 0.001 |
| NF | NM | 0.070 | 0.052 | 0.087 | 15.272 | 0.001 |
| NF_NM | PG | 0.427 | 0.416 | 0.437 | 161.944 | 0.001 |
| NF_NM | PE | 0.280 | 0.266 | 0.295 | 73.556 | 0.001 |
| NF_NM | NM | 0.148 | 0.129 | 0.167 | 30.441 | 0.001 |
| PG | PE | 0.146 | 0.135 | 0.157 | 49.816 | 0.001 |
| PG | NM | 0.279 | 0.262 | 0.295 | 66.260 | 0.001 |
| PE | NM | 0.133 | 0.113 | 0.152 | 26.358 | 0.001 |

**b) South boreal**

| group1 | group2 | Diff | Lower | Upper | q-value | p-value |
|---|---|---|---|---|---|---|
| PG | PE | 0.114 | 0.093 | 0.134 | 21.241 | 0.001 |
| PG | NF | 0.397 | 0.381 | 0.412 | 96.913 | 0.001 |
| PG | NM | 0.249 | 0.217 | 0.281 | 29.783 | 0.001 |
| PG | NF_NM | 0.421 | 0.365 | 0.476 | 29.159 | 0.001 |
| PE | NF | 0.283 | 0.259 | 0.307 | 45.286 | 0.001 |
| PE | NM | 0.135 | 0.098 | 0.172 | 14.090 | 0.001 |
| PE | NF_NM | 0.307 | 0.248 | 0.366 | 20.223 | 0.001 |
| NF | NM | 0.148 | 0.113 | 0.182 | 16.475 | 0.001 |
| NF | NF_NM | 0.024 | -0.033 | 0.081 | 1.633 | 0.750 |
| NM | NF_NM | 0.172 | 0.108 | 0.235 | 10.418 | 0.001 |

**c) Hemiboreal**

| group1 | group2 | Diff | Lower | Upper | q-value | p-value |
|---|---|---|---|---|---|---|
| PE | NF | 0.153 | 0.121 | 0.186 | 18.247 | 0.001 |
| PE | PG | 0.117 | 0.097 | 0.137 | 22.570 | 0.001 |
| PE | NF_NM | 0.230 | 0.191 | 0.270 | 22.482 | 0.001 |
| PE | NM | 0.140 | 0.108 | 0.173 | 16.623 | 0.001 |
| NF | PG | 0.271 | 0.244 | 0.297 | 38.934 | 0.001 |
| NF | NF_NM | 0.077 | 0.034 | 0.120 | 6.851 | 0.001 |
| NF | NM | 0.013 | -0.024 | 0.050 | 1.354 | 0.862 |
| PG | NF_NM | 0.348 | 0.313 | 0.383 | 38.230 | 0.001 |
| PG | NM | 0.258 | 0.231 | 0.285 | 36.820 | 0.001 |
| NF_NM | NM | 0.090 | 0.047 | 0.134 | 7.991 | 0.001 |

**d) Nemoral**

| group1 | group2 | Diff | Lower | Upper | q-value | p-value |
|---|---|---|---|---|---|---|
| PG | NM | 0.266 | 0.172 | 0.361 | 10.312 | 0.001 |
| PG | NF | 0.367 | 0.168 | 0.566 | 6.761 | 0.001 |
| PG | PE | 0.215 | 0.093 | 0.337 | 6.442 | 0.001 |

| | | | | | | |
|---|---|---|---|---|---|---|
| NM | NF | 0.101 | -0.116 | 0.318 | 1.707 | 0.608 |
| NM | PE | 0.052 | -0.098 | 0.201 | 1.267 | 0.783 |
| NF | PE | 0.153 | -0.077 | 0.383 | 2.430 | 0.316 |

#### Supplementary Table 3

The results of external validation with the Sveaskog forest management compartment dataset (stand-level; n=57548 polygons). The table presents the basic statistics of predicted values of relative likelihood of HCVF occurrence for stand polygons belonging to different "Forest Naturalness" categories (binary variable).

| Region | Naturalness | Mean | Count | STD | CI95_low | CI95_high |
|---|---|---|---|---|---|---|
| North boreal | No | 0.299 | 34130 | 0.172 | 0.297 | 0.301 |
| North boreal | Yes | 0.649 | 7654 | 0.200 | 0.645 | 0.654 |
| South boreal | No | 0.296 | 6306 | 0.182 | 0.291 | 0.300 |
| South boreal | Yes | 0.634 | 1683 | 0.212 | 0.624 | 0.645 |
| Hemiboreal | No | 0.330 | 6602 | 0.166 | 0.326 | 0.334 |
| Hemiboreal | Yes | 0.578 | 917 | 0.188 | 0.566 | 0.590 |
| Nemoral | No | 0.386 | 233 | 0.159 | 0.366 | 0.407 |
| Nemoral | Yes | 0.647 | 23 | 0.171 | 0.577 | 0.717 |

#### Supplementary Table 4

The results of external validation with the Sveaskog forest management compartment dataset (stand-level; n=57548 polygons). The table summarizes the Tukey's posthoc tests to check for statistical significance of differences between predicted values of relative likelihood of HCVF occurrence for stand polygons belonging to different "Forest naturalness" categories (binary variable).

**a) North boreal**

| group1 | group2 | Diff | Lower | Upper | q-value | p-value |
|---|---|---|---|---|---|---|
| Yes | No | 0.352 | 0.348 | 0.357 | 218.033 | 0.001 |

**b) South boreal**

| group1 | group2 | Diff | Lower | Upper | q-value | p-value |
|---|---|---|---|---|---|---|
| No | Yes | 0.347 | 0.337 | 0.358 | 93.380 | 0.001 |

**c) Hemiboreal**

| group1 | group2 | Diff | Lower | Upper | q-value | p-value |
|---|---|---|---|---|---|---|
| No | Yes | 0.248 | 0.236 | 0.260 | 58.224 | 0.001 |

**d) Nemoral**

| group1 | group2 | Diff | Lower | Upper | q-value | p-value |
|---|---|---|---|---|---|---|
| No | Yes | 0.266 | 0.195 | 0.337 | 10.477 | 0.001 |

#### Supplementary Table 5

The results of external validation with the NFI (plot-level; n=13775 plots). The table presents the basic statistics of predicted values of relative likelihood of HCVF occurrence for stand polygons belonging to different "Forest naturalness" categories: "plantation", "normal", "natural".

| Region | Naturalness | Mean | Count | STD | CI95_low | CI95_high |
|---:|---:|---:|---:|---:|---:|---:|
| North boreal | plantation | 0.201 | 19 | 0.117 | 0.148 | 0.253 |
| North boreal | normal | 0.361 | 2897 | 0.250 | 0.352 | 0.370 |
| North boreal | natural | 0.840 | 156 | 0.176 | 0.813 | 0.868 |
| South boreal | plantation | 0.240 | 74 | 0.142 | 0.208 | 0.273 |
| South boreal | normal | 0.372 | 5200 | 0.231 | 0.366 | 0.379 |
| South boreal | natural | 0.761 | 123 | 0.231 | 0.720 | 0.801 |
| Hemiboreal | plantation | 0.273 | 103 | 0.186 | 0.237 | 0.308 |
| Hemiboreal | normal | 0.375 | 4494 | 0.208 | 0.369 | 0.381 |
| Hemiboreal | natural | 0.631 | 26 | 0.206 | 0.551 | 0.710 |
| Nemoral | plantation | 0.387 | 69 | 0.174 | 0.346 | 0.428 |
| Nemoral | normal | 0.419 | 612 | 0.202 | 0.403 | 0.435 |
| Nemoral | natural | 0.423 | 2 | 0.299 | 0.008 | 0.837 |

### Supplementary Table 6

The results of external validation with the NFI (plot-level; n=13775 plots). The table summarizes the Tukey's posthoc tests to check for statistical significance of differences between predicted values of relative likelihood of HCVF occurrence for stand polygons belonging to different "Forest naturalness" categories: "plantation", "normal", "natural".

**a) North boreal**

| group1 | group2 | Diff | Lower | Upper | q-value | p-value |
|---|---|---|---|---|---|---|
| normal | natural | 0.479 | 0.432 | 0.527 | 33.515 | 0.001 |
| normal | plantation | 0.160 | 0.027 | 0.293 | 3.992 | 0.013 |
| natural | plantation | 0.639 | 0.499 | 0.780 | 15.118 | 0.001 |

**b) South boreal**

| group1 | group2 | Diff | Lower | Upper | q-value | p-value |
|---|---|---|---|---|---|---|
| normal | natural | 0.388 | 0.339 | 0.437 | 26.183 | 0.001 |
| normal | plantation | 0.132 | 0.069 | 0.195 | 6.947 | 0.001 |
| natural | plantation | 0.520 | 0.441 | 0.600 | 21.764 | 0.001 |

**c) Hemiboreal**

| group1 | group2 | Diff | Lower | Upper | q-value | p-value |
|---|---|---|---|---|---|---|
| normal | plantation | 0.103 | 0.054 | 0.151 | 7.015 | 0.001 |
| normal | natural | 0.256 | 0.160 | 0.351 | 8.863 | 0.001 |
| plantation | natural | 0.358 | 0.251 | 0.465 | 11.128 | 0.001 |

**d) Nemoral**

| group1 | group2 | Diff | Lower | Upper | q-value | p-value |
|---|---|---|---|---|---|---|
| natural | normal | 0.003 | -0.329 | 0.335 | 0.033 | 0.900 |
| natural | plantation | 0.036 | -0.300 | 0.372 | 0.353 | 0.900 |
| normal | plantation | 0.032 | -0.027 | 0.092 | 1.812 | 0.408 |

#### Supplementary Table 7

The results of external validation with the NFI (plot-level; n=13775 plots). The table presents the basic statistics of predicted values of relative likelihood of HCVF occurrence for stand polygons belonging to different "Natura2000" categories (binary variable).

| Region | Natura2000 | Mean | Count | STD | CI95_low | CI95_high |
|---|---|---|---|---|---|---|
| North boreal | No | 0.348 | 2841 | 0.236 | 0.339 | 0.356 |
| North boreal | Yes | 0.832 | 231 | 0.235 | 0.802 | 0.862 |
| South boreal | No | 0.364 | 5216 | 0.223 | 0.358 | 0.370 |
| South boreal | Yes | 0.833 | 181 | 0.191 | 0.805 | 0.860 |
| Hemiboreal | No | 0.367 | 4528 | 0.203 | 0.361 | 0.373 |
| Hemiboreal | Yes | 0.722 | 95 | 0.181 | 0.686 | 0.758 |
| Nemoral | No | 0.407 | 660 | 0.195 | 0.392 | 0.421 |
| Nemoral | Yes | 0.687 | 23 | 0.136 | 0.632 | 0.743 |

#### Supplementary Table 8

The results of external validation with the NFI (plot-level; n=13775 plots). The table summarizes the Tukey's posthoc tests to check for statistical significance of differences between predicted values of relative likelihood of HCVF occurrence for stand polygons belonging to different "Natura2000" categories (binary variable).

**a) North boreal**

| group1 | group2 | Diff | Lower | Upper | q-value | p-value |
|---|---|---|---|---|---|---|
| No | Yes | 0.484 | 0.452 | 0.516 | 42.450 | 0.000 |

**b) South boreal**

| group1 | group2 | Diff | Lower | Upper | q-value | p-value |
|---|---|---|---|---|---|---|
| No | Yes | 0.469 | 0.436 | 0.502 | 39.481 | 0.000 |

**c) Hemiboreal**

| group1 | group2 | 0.000 | 0.000 | 0.000 | 0.000 | 0.000 |
|---|---|---|---|---|---|---|
| No | Yes | 0.355 | 0.314 | 0.396 | 23.888 | 0.000 |

**d) Nemoral**

| group1 | group2 | 0.000 | 0.000 | 0.000 | 0.000 | 0.000 |
|---|---|---|---|---|---|---|
| No | Yes | 0.280 | 0.200 | 0.361 | 9.697 | 0.000 |

#### Supplementary Table 9

To test the robustness of our model's specification (M1, code "selected"), we compared its performance against seven alternative models trained and validated using the same data: (M2) the "global" model using data from the entire Sweden (i.e. without the regional stratification; (M3) the "full" model trained with all spatial predictors, including all derived multi-scale features, and ignoring a strong (> 0.7) pairwise correlation between some variables; (M4) the "onescale" model trained with all spatial predictors but excluding all multi-scale features; (M5) the "baseline" model trained using only 4 key spatial predictors, all hypothetically having a strong effect on the probability of HCVF occurrence (elevation, tree height, distance to roads and % of logged areas within 1ha);  Additionally, the "full", "onescale" and "baseline" models were trained with and without longitude and latitude as the auxiliary variables. The results of 10-fold Spatial Cross-Validation (SCV) confirmed the robustness of our model's specification as it performed best or equally good as the "full" model for all regions.

| Region | Model | Accuracy | TSS | ROC AUC | PR AUC | Pearson r | Briers score | MCC |
|---|---|---|---|---|---|---|---|---|
| North boreal | (M3a) full | 0.8146 | 0.6226 | 0.8958 | 0.8899 | 0.6855 | 0.1355 | 0.6276 |
| North boreal | (M3b) full_nolatlon | 0.8096 | 0.6082 | 0.8946 | 0.8892 | 0.6841 | 0.1360 | 0.6170 |
| **North boreal** | **(M1) selected** | **0.8111** | **0.6077** | **0.8944** | **0.8909** | **0.6827** | **0.1377** | **0.6215** |
| North boreal | (M4a) onescale | 0.7965 | 0.5825 | 0.8805 | 0.8734 | 0.6595 | 0.1428 | 0.5907 |
| North boreal | (M4b) onescale_nolatlon | 0.7928 | 0.5866 | 0.8760 | 0.8666 | 0.6515 | 0.1455 | 0.5852 |
| North boreal | (M5a) baseline | 0.7587 | 0.5074 | 0.8387 | 0.8314 | 0.5882 | 0.1622 | 0.5133 |
| North boreal | (M5b) baseline_nolatlon | 0.7193 | 0.4241 | 0.7842 | 0.7515 | 0.4922 | 0.1912 | 0.4366 |
| South boreal | (M3a) full | 0.8162 | 0.6193 | 0.8999 | 0.8781 | 0.6908 | 0.1335 | 0.6260 |
| South boreal | (M3b) full_nolatlon | 0.8151 | 0.6098 | 0.8985 | 0.8754 | 0.6887 | 0.1343 | 0.6247 |
| **South boreal** | **(M1) selected** | **0.8124** | **0.6089** | **0.8978** | **0.8716** | **0.6861** | **0.1357** | **0.6176** |
| South boreal | (M4a) onescale | 0.7996 | 0.5867 | 0.8836 | 0.8502 | 0.6615 | 0.1412 | 0.5907 |
| South boreal | (M4b) onescale_nolatlon | 0.7989 | 0.5921 | 0.8812 | 0.8462 | 0.6583 | 0.1423 | 0.5912 |
| South boreal | (M5a) baseline | 0.7499 | 0.4967 | 0.8295 | 0.7947 | 0.5682 | 0.1677 | 0.4923 |
| South boreal | (M5b) baseline_nolatlon | 0.7307 | 0.4533 | 0.8080 | 0.7683 | 0.5300 | 0.1804 | 0.4562 |
| Hemiboreal | (M3a) full | 0.8068 | 0.6030 | 0.8942 | 0.8440 | 0.6661 | 0.1414 | 0.6093 |
| **Hemiboreal** | **(M1) selected** | **0.8082** | **0.5918** | **0.8932** | **0.8418** | **0.6668** | **0.1418** | **0.6096** |
| Hemiboreal | (M3b) full_nolatlon | 0.8067 | 0.5953 | 0.8908 | 0.8385 | 0.6624 | 0.1421 | 0.6070 |
| Hemiboreal | (M4a) onescale | 0.7805 | 0.5392 | 0.8592 | 0.7928 | 0.6058 | 0.1582 | 0.5506 |
| Hemiboreal | (M4b) onescale_nolatlon | 0.7746 | 0.5184 | 0.8498 | 0.7746 | 0.5904 | 0.1616 | 0.5377 |
| Hemiboreal | (M5a) baseline | 0.7073 | 0.4035 | 0.7806 | 0.6791 | 0.4724 | 0.1916 | 0.4073 |
| Hemiboreal | (M5b) baseline_nolatlon | 0.6764 | 0.3576 | 0.7399 | 0.6274 | 0.4047 | 0.2113 | 0.3472 |
| Nemoral | (M3a) full | 0.8156 | 0.4991 | 0.9091 | 0.8809 | 0.6966 | 0.1394 | 0.6354 |
| Nemoral | (M3b) full_nolatlon | 0.8079 | 0.5263 | 0.9020 | 0.8696 | 0.6855 | 0.1411 | 0.6121 |
| **Nemoral** | **(M1) selected** | **0.8061** | **0.5679** | **0.8921** | **0.8497** | **0.6588** | **0.1526** | **0.6115** |
| Nemoral | (M4b) onescale_nolatlon | 0.7859 | 0.5451 | 0.8611 | 0.8043 | 0.6132 | 0.1611 | 0.5730 |
| Nemoral | (M4a) onescale | 0.7898 | 0.5074 | 0.8591 | 0.8112 | 0.6117 | 0.1603 | 0.5639 |
| Nemoral | (M5a) baseline | 0.6560 | 0.2462 | 0.7161 | 0.5662 | 0.3538 | 0.2200 | 0.3245 |
| Nemoral | (M5b) baseline_nolatlon | 0.6289 | 0.1876 | 0.6851 | 0.5750 | 0.3198 | 0.2296 | 0.2584 |
| **All** | **(M2) global** | **0.8149** | **0.6314** | **0.8970** | **0.8729** | **0.6836** | **0.1364** | **0.6270** |

### Supplementary Table 10

The results of the hyperparameters tuning for the two best-performing models (i.e., the full model and the model with preselected variables only) using Bayesian optimization with Gaussian processes implemented in the Python package scikit-optimize v0.9.0. We tuned the following hyperparameters (a range of values and data type in parentheses): max_depth (3, 30; integer), min_samples_split (0.0001, 0.3; real; fraction of samples), min_samples_leaf (0.0001, 0.3; real; fraction of samples), max_features (0.1, 1.0; real; fraction), criterion ("gini", "entropy", "log_loss"; categorical). We ran the optimization algorithm for 100 iterations (with nested 5-fold SCV) for each model (2) and region (4) and, to speed up computations, we fixed the number of estimators (trees) to 200. For both models and for all regions, the difference in performance of tuned models in comparison to models run with default hyperparameters (except the number of trees that was set to 500) was < 0.5% as measured by ROC AUC (compare b) with Table 3 in the main text and see the compiled Jupyter notebooks). This confirms that default values are robust for our specific application.

a) The full model (“M3a”)

| region | accuracy | roc_auc | pr_auc | pearson_r | briers_score | MCC |
|---|---|---|---|---|---|---|
| 1 | 0.812498 | 0.898076 | 0.893445 | 0.692487 | 0.131931 | 0.624822 |
| 2 | 0.816374 | 0.898897 | 0.875666 | 0.691302 | 0.131223 | 0.625582 |
| 3 | 0.802748 | 0.887519 | 0.83507 | 0.659658 | 0.142107 | 0.596534 |
| 4 | 0.815944 | 0.906412 | 0.877308 | 0.693585 | 0.139676 | 0.625998 |

| region | criterion | max_depth | min_samples_split | min_samples_leaf | max_features |
|---|---|---|---|---|---|
| 1 | log_loss | 24 | 0.0001 | 0.0001 | 1 |
| 2 | log_loss | 30 | 0.0001 | 0.0001 | 0.562738 |
| 3 | log_loss | 30 | 0.0001 | 0.0001 | 0.1 |
| 4 | log_loss | 30 | 0.0001 | 0.0001 | 0.1 |

b) The model with preselected variables (“M1”)

| region | accuracy | roc_auc | pr_auc | pearson_r | briers_score | MCC |
|---|---|---|---|---|---|---|
| 1 | 0.811888 | 0.891874 | 0.883525 | 0.680121 | 0.135028 | 0.621279 |
| 2 | 0.809701 | 0.895349 | 0.868471 | 0.682789 | 0.133356 | 0.611816 |
| 3 | 0.812218 | 0.892466 | 0.839481 | 0.669244 | 0.138165 | 0.616012 |
| 4 | 0.79805 | 0.888916 | 0.840164 | 0.661486 | 0.152758 | 0.584113 |

| region | criterion | max_depth | min_samples_split | min_samples_leaf | max_features |
|---|---|---|---|---|---|
| 1 | entropy | 28 | 0.0001 | 0.0001 | 1 |
| 2 | entropy | 27 | 0.0001 | 0.0001 | 0.420203 |
| 3 | log_loss | 24 | 0.0001 | 0.0001 | 0.426681 |
| 4 | log_loss | 30 | 0.0001 | 0.0001 | 0.1 |
