## Supplementary material for "Mapping forests with different levels of naturalness using machine learning and landscape data mining": Jupyter Notebooks: M1_selected.html

validation


In [2]:

```
%load_ext autoreload
%autoreload 2

import warnings
warnings.filterwarnings("ignore")

from matplotlib import pyplot as plt
import matplotlib as mpl
mpl.rcParams["figure.dpi"] = 150

import json
import sys
import numpy as np
import pandas
import seaborn as sns
sns.set_style("white")

sys.path.insert(0, "/app/src")

from plots import plots
from config import settings
```

```
The autoreload extension is already loaded. To reload it, use:
  %reload_ext autoreload
```

In [3]:

```
with open("./config.json", "r") as _file:
    mconfig = json.load(_file)
print(json.dumps(mconfig, indent=2))

model_name = mconfig["model_metadata"]["name"]
target_var = mconfig["model_specification"]["target_variable"]
region_var = mconfig["model_specification"]["stratification_layer"]
```

```
{
  "model_metadata": {
    "name": "S1_selected",
    "description": [
      "Full model using selected variables expressed at all target spatial scales. ",
      "Maximum allowed correlation between variables < 0.7. Seperate model for each ",
      "each region is fitted."
    ]
  },
  "model_data": {
    "samples_layer": "SAMPLES_1"
  },
  "model_specification": {
    "target_variable": "HCVF",
    "stratification_layer": "ecoregions",
    "stratification_submodels": [
      {
        "regions": [
          1
        ],
        "variables_list": [
          "HEIGHT005",
          "SLOPE011",
          "DEM011",
          "HEIGHTc",
          "HEIGHTv",
          "HANSEN005",
          "FOPEN003",
          "HEIGHTc011",
          "ROADS051",
          "ARTI011",
          "BUILTUPd",
          "POPd",
          "FOPEN011",
          "SLOPEv",
          "AGRI051",
          "FOPEN051",
          "POP051",
          "GAPS",
          "ARTI003",
          "UNDERSTORYv",
          "HANSEN",
          "OPENnat011",
          "SHAFOR051",
          "OPENwet011",
          "UNDERSTORY",
          "SHANAT005",
          "POP003",
          "UNDERSTORY051",
          "GAPSv",
          "PROD005",
          "WATER005",
          "SLOPE101",
          "ROADS",
          "LIGHTS101",
          "WATER101",
          "HEIGHTc051",
          "WATERd",
          "AGRI003",
          "OPENwet101",
          "LAT",
          "PROD101",
          "BROADLEAF",
          "OPENwet",
          "HEIGHT101",
          "BROADLEAF101",
          "WATER",
          "OPENnat",
          "HEIGHTc101"
        ]
      },
      {
        "regions": [
          2
        ],
        "variables_list": [
          "HEIGHTc005",
          "HEIGHT",
          "HANSEN003",
          "SLOPE011",
          "FOPEN003",
          "ROADS011",
          "ARTId",
          "HEIGHTv",
          "HANSEN011",
          "DEM003",
          "GAPS",
          "FOPEN",
          "ARTI003",
          "UNDERSTORY",
          "BROADLEAF005",
          "ARTI051",
          "BUILTUPd",
          "SHAFOR005",
          "POPd",
          "SLOPEv",
          "GAPSvtmr",
          "UNDERSTORY011",
          "PROD005",
          "OPENwet011",
          "ROADS",
          "OPENnat051",
          "AGRI011",
          "HANSEN051",
          "AGRI051",
          "GAPSv",
          "SHANAT051",
          "OPENnat003",
          "POP011",
          "PROD051",
          "WATER011",
          "SHANAT011",
          "BROADLEAF101",
          "HEIGHTc051",
          "WATER101",
          "WATERd",
          "SLOPE101",
          "OPENwet101",
          "WATER003",
          "AGRI003",
          "HEIGHTc101",
          "WATER",
          "SHAFOR101",
          "LAT",
          "OPENwet",
          "LON"
        ]
      },
      {
        "regions": [
          3
        ],
        "variables_list": [
          "HANSEN005",
          "FOPEN011",
          "BROADLEAF005",
          "HEIGHTc003",
          "FOPEN003",
          "HANSEN",
          "ROADS011",
          "WATER011",
          "HEIGHT011",
          "ROADSd",
          "ARTId",
          "SLOPE005",
          "ARTI003",
          "WATERd",
          "SHAFOR005",
          "OPENwet011",
          "BUILTUPd",
          "FOPEN",
          "AGRI011",
          "GAPS005",
          "ROADS051",
          "OPENwet051",
          "BROADLEAF051",
          "POP011",
          "PROD005",
          "HEIGHTc051",
          "WATER003",
          "DEM005",
          "UNDERSTORY005",
          "AGRI101",
          "GAPSvtmr",
          "HANSEN051",
          "UNDERSTORYv",
          "SLOPEv",
          "OPENnat005",
          "OPENnat051",
          "PROD051",
          "SHANAT051",
          "HEIGHTv",
          "AGRI003",
          "WATER101",
          "GAPS051",
          "ROADS",
          "UNDERSTORY051",
          "WATER",
          "OPENnat",
          "SHAFOR101",
          "GAPSv",
          "LON",
          "OPENwet",
          "HEIGHTc101",
          "LAT",
          "HEIGHT101"
        ]
      },
      {
        "regions": [
          4
        ],
        "variables_list": [
          "BROADLEAF005",
          "SLOPE011",
          "HEIGHT011",
          "HEIGHTc011",
          "FOPEN005",
          "HEIGHT",
          "SLOPEv",
          "SHAFOR003",
          "HANSEN003",
          "BROADLEAF051",
          "AGRI051",
          "GAPS011",
          "HEIGHTc051",
          "SHAFOR051",
          "WATER005",
          "AGRI005",
          "HEIGHTv",
          "DEM051",
          "PROD005",
          "ARTI003",
          "OPENnat101",
          "WATER051",
          "POP003",
          "UNDERSTORY003",
          "UNDERSTORY011",
          "LIGHTS051",
          "GAPSvtmr",
          "WATERd",
          "PROD101",
          "FOPEN",
          "GAPSv",
          "OPENwet011",
          "ROADS011",
          "OPENnat",
          "HEIGHT051",
          "GAPS",
          "ROADS",
          "BUILTUPd",
          "HEIGHTc101",
          "WATER",
          "POPd",
          "OPENnat011",
          "ROADSd",
          "UNDERSTORYv",
          "LAT",
          "OPENnat003",
          "OPENwet",
          "LON"
        ]
      }
    ],
    "lat_field": "LAT",
    "lon_field": "LON",
    "target_epsg": 3006,
    "variables_list": [],
    "variables_drop_regexp": "",
    "variables_keep_regexp": "",
    "variables_latlon": true,
    "balanced_rf": true
  },
  "model_crossvalidation": {
    "n_splits": 10,
    "spatial": true,
    "spatial_grid": "grid_20.geojson"
  },
  "model_hyperparameters": {
    "n_estimators": 500,
    "oob_score": true
  },
  "prediction": {
    "predict": true
  }
}
```

In [4]:

```
models = plots.load_models("./", regexp=fr"{model_name}_R[1-5]*.pickle")
models_stats = plots.get_cv_validation_stats(models)
```

```
Loading S1_selected_R1.pickle...
Loading S1_selected_R2.pickle...
Loading S1_selected_R3.pickle...
Loading S1_selected_R4.pickle...
```

In [5]:

```
models[0].cv_results.shape
```

Out[5]:

```
(10, 13)
```

In [6]:

```
print(models[0].data.shape)
models_labels=["North boreal", "South boreal", "Hemiboreal", "Nemoral"]

g = sns.FacetGrid(
    data=models[0].data, col=region_var, aspect=0.7, 
    sharey=False
)
g.map_dataframe(sns.countplot, x=target_var, palette=["black","red"])
axes = g.axes.flatten()
for i, ax in enumerate(axes):
    ax.set_title(models_labels[i], fontweight="bold", fontsize=16)
```

```
(14090, 130)
```

In [7]:

```
plots.plot_validation_curves_multimodel(
    models_stats, colors="bgrcmykw", figsize=(10, 5), plot_std=True, std_alpha=0.1,
    models_labels=["North boreal", "South boreal", "Hemiboreal", "Nemoral"],
    save_name="Figure2.png"
)
```

In [9]:

```
# performance metrics
means, stds = plots.get_metrics(models_stats)
display(means)
display(stds)
```

|  | accuracy | TSS | roc\_auc | pr\_auc | pearson\_r | briers\_score | MCC |
| --- | --- | --- | --- | --- | --- | --- | --- |
| 0 | 0.811071 | 0.607707 | 0.894391 | 0.890909 | 0.682667 | 0.137704 | 0.621510 |
| 1 | 0.812354 | 0.608865 | 0.897814 | 0.871556 | 0.686108 | 0.135682 | 0.617628 |
| 2 | 0.808228 | 0.591838 | 0.893231 | 0.841799 | 0.666823 | 0.141793 | 0.609604 |
| 3 | 0.806138 | 0.567882 | 0.892063 | 0.849692 | 0.658849 | 0.152614 | 0.611538 |

|  | accuracy | TSS | roc\_auc | pr\_auc | pearson\_r | briers\_score | MCC |
| --- | --- | --- | --- | --- | --- | --- | --- |
| 0 | 0.019681 | 0.045446 | 0.014750 | 0.019952 | 0.026054 | 0.007905 | 0.039076 |
| 1 | 0.020558 | 0.046115 | 0.018678 | 0.038806 | 0.036213 | 0.008786 | 0.046483 |
| 2 | 0.019474 | 0.064574 | 0.015308 | 0.024282 | 0.028949 | 0.009160 | 0.036099 |
| 3 | 0.051995 | 0.164805 | 0.032248 | 0.054130 | 0.056224 | 0.017430 | 0.090672 |

In [50]:

```
# number of variables per model
for i, model in enumerate(models):
    print(f"Model_{i+1}: ", len(model.cov_names))
```

```
Model_1:  49
Model_2:  50
Model_3:  53
Model_4:  48
```

In [51]:

```
# plot features Gini's importances
for i, model in enumerate(models):
    print(models_labels[i])
    _plot = plots.plot_gini_importances(model, figsize=(14, 10), n=50)
```

```
North boreal
```

```
South boreal
```

```
Hemiboreal
```

```
Nemoral
```

##### Partial Dependence Plot¶

In [52]:

```
plots.plot_partial_dependences(models, n_features=6)
```

##### Accumulated Local Effects Plot¶

In [53]:

```
plots.plot_ales(models, n_features=6, xlims=(-2, 2))
```

In [54]:

```
# average max tree depth per model
import numpy as np
for i, model in enumerate(models):
    max_depths = []
    for tree_idx, est in enumerate(model.model.estimators_):
        max_depths.append(est.tree_.max_depth)
    print(f"{models_labels[i]}: ", np.array(max_depths).mean())
```

```
North boreal:  21.492
South boreal:  20.882
Hemiboreal:  19.87
Nemoral:  11.546
```

### Validation¶

#### Sveaskog validation¶

tree stand (polygon) level validation; minimum 10 pixels with predicted value

In [55]:

```
# load a dataframe with predictions and Sveaskog atributes
svea_df = pandas.read_csv("svea_predictions.csv")
svea_df = plots.svea_preprocess(svea_df)

# filter out data not reliable for validation
# min number of 1 ha pixels to calculate the mean probability
svea_df = svea_df[svea_df.pred_count > 10]

# show the number of validation polygons per region
display(svea_df.region.value_counts())
print("TOTAL POLYGONS N: ", svea_df.region.value_counts().sum())
```

```
1    41784
2     7989
3     7519
4      256
Name: region, dtype: int64
```

```
TOTAL POLYGONS N:  57548
```

##### Boxplot -> forest management classes¶

"NF" (conservation, no management)
"NF\_NM" (conservation, no yet specified)
"NM" (conservation, management)
"PE" (production forest, increased conservation considerations)
"PG" (production forest, general conservation conciderations)

Management categories for stands. A goal for long-term management is set for each stand, in one of four categories: (i) NF = Nature conservation, free development; (ii) NM = Nature conservation-oriented management, often implying restoration measures; (iii) PG = Production with general conservation concern (green-tree and deadwood retention for biodiversity at harvest, represented by black dots). The minimum level of retention is specified, and always >15 % of the harvested area; (iv) PE = Production with enhanced conservation concern.

In [56]:

```
plots.plot_boxplots(
    svea_df, "management", height=4, aspect=0.75, fig_title="a) Stand-level validation - Forest management objectives",
    save_name="Figure4a.png", width=0.85
)
```

In [57]:

```
plots.describe_ci(svea_df, "pred_mean", ["region", "management"])
```

Out[57]:

|  |  | mean | count | std | ci95\_lo | ci95\_hi |
| --- | --- | --- | --- | --- | --- | --- |
| region | management |  |  |  |  |  |
| 1 | NF | 0.637480 | 4148 | 0.203391 | 0.631291 | 0.643670 |
|  | NF\_NM | 0.716035 | 2401 | 0.164490 | 0.709456 | 0.722615 |
|  | NM | 0.569681 | 903 | 0.194839 | 0.556973 | 0.582389 |
|  | PE | 0.416132 | 2516 | 0.188875 | 0.408751 | 0.423512 |
|  | PG | 0.290654 | 31816 | 0.168310 | 0.288805 | 0.292504 |
| 2 | NF | 0.673201 | 1245 | 0.191506 | 0.662563 | 0.683839 |
|  | NF\_NM | 0.697029 | 82 | 0.172051 | 0.659789 | 0.734268 |
|  | NM | 0.524889 | 253 | 0.205333 | 0.499587 | 0.550191 |
|  | PE | 0.396282 | 868 | 0.200499 | 0.382944 | 0.409621 |
|  | PG | 0.281819 | 5541 | 0.175251 | 0.277205 | 0.286434 |
| 3 | NF | 0.593441 | 297 | 0.183688 | 0.572550 | 0.614332 |
|  | NF\_NM | 0.666963 | 172 | 0.160221 | 0.643018 | 0.690908 |
|  | NM | 0.580199 | 303 | 0.185349 | 0.559328 | 0.601069 |
|  | PE | 0.431979 | 608 | 0.180530 | 0.417629 | 0.446330 |
|  | PG | 0.322415 | 6139 | 0.161501 | 0.318375 | 0.326455 |
| 4 | NF | 0.727443 | 5 | 0.060889 | 0.674071 | 0.780815 |
|  | NF\_NM | NaN | 0 | NaN | NaN | NaN |
|  | NM | 0.631810 | 19 | 0.183305 | 0.549387 | 0.714234 |
|  | PE | 0.592005 | 13 | 0.164902 | 0.502363 | 0.681647 |
|  | PG | 0.372429 | 219 | 0.148708 | 0.352733 | 0.392125 |

In [58]:

```
plots.tukey_subsets(svea_df, "pred_mean", "management", "region")
```

```
region: 1

  group1 group2      Diff     Lower     Upper     q-value  p-value
0     NF  NF_NM  0.078124  0.065830  0.090417   24.516707    0.001
1     NF     PG  0.348378  0.340458  0.356298  169.699296    0.001
2     NF     PE  0.202319  0.189059  0.215579   58.861092    0.001
3     NF     NM  0.069806  0.052173  0.087440   15.271846    0.001
4  NF_NM     PG  0.426502  0.416342  0.436662  161.943916    0.001
5  NF_NM     PE  0.280443  0.265734  0.295151   73.556231    0.001
6  NF_NM     NM  0.147930  0.129183  0.166677   30.441261    0.001
7     PG     PE  0.146059  0.134748  0.157370   49.815737    0.001
8     PG     NM  0.278572  0.262353  0.294791   66.260177    0.001
9     PE     NM  0.132513  0.113118  0.151908   26.357702    0.001


region: 2

  group1 group2      Diff     Lower     Upper    q-value   p-value
0     PG     PE  0.113750  0.093087  0.134414  21.241189  0.001000
1     PG     NF  0.396645  0.380853  0.412437  96.912573  0.001000
2     PG     NM  0.249043  0.216777  0.281308  29.782553  0.001000
3     PG  NF_NM  0.420791  0.365109  0.476474  29.159271  0.001000
4     PE     NF  0.282895  0.258791  0.306999  45.285589  0.001000
5     PE     NM  0.135292  0.098243  0.172342  14.090187  0.001000
6     PE  NF_NM  0.307041  0.248457  0.365625  20.222820  0.001000
7     NF     NM  0.147602  0.113032  0.182172  16.474705  0.001000
8     NF  NF_NM  0.024146 -0.032902  0.081195   1.633172  0.749959
9     NM  NF_NM  0.171749  0.108139  0.235358  10.418266  0.001000


region: 3

  group1 group2      Diff     Lower     Upper    q-value   p-value
0     PE     NF  0.153445  0.120996  0.185894  18.246654  0.001000
1     PE     PG  0.117185  0.097151  0.137220  22.569601  0.001000
2     PE  NF_NM  0.230495  0.190935  0.270055  22.481917  0.001000
3     PE     NM  0.140411  0.107819  0.173004  16.623060  0.001000
4     NF     PG  0.270630  0.243809  0.297451  38.934267  0.001000
5     NF  NF_NM  0.077050  0.033657  0.120443   6.851377  0.001000
6     NF     NM  0.013034 -0.024118  0.050186   1.353687  0.862098
7     PG  NF_NM  0.347680  0.312588  0.382772  38.229900  0.001000
8     PG     NM  0.257597  0.230602  0.284591  36.820446  0.001000
9  NF_NM     NM  0.090084  0.046583  0.133585   7.990528  0.001000


region: 4

  group1 group2      Diff     Lower     Upper    q-value   p-value
0     PG     NM  0.266247  0.171752  0.360741  10.311813  0.001000
1     PG     NF  0.367293  0.168459  0.566127   6.760504  0.001000
2     PG     PE  0.214608  0.092686  0.336529   6.442029  0.001000
3     NM     NF  0.101046 -0.115584  0.317677   1.707094  0.608433
4     NM     PE  0.051639 -0.097555  0.200832   1.266724  0.782804
5     NF     PE  0.152685 -0.077238  0.382608   2.430363  0.316499
```

##### Boxplot -> forest naturalness classes¶

In [59]:

```
plots.plot_boxplots(
    svea_df, "naturalness", height=4, aspect=0.75, fig_title="b) Stand-level validation - Forest naturalness",
    save_name="Figure4b.png", width=0.5
)
```

In [60]:

```
plots.describe_ci(svea_df, "pred_mean", ["region", "naturalness"])
```

Out[60]:

|  |  | mean | count | std | ci95\_lo | ci95\_hi |
| --- | --- | --- | --- | --- | --- | --- |
| region | naturalness |  |  |  |  |  |
| 1 | No | 0.298961 | 34130 | 0.172301 | 0.297133 | 0.300789 |
|  | Yes | 0.649176 | 7654 | 0.199848 | 0.644699 | 0.653653 |
| 2 | No | 0.295876 | 6306 | 0.182034 | 0.291383 | 0.300369 |
|  | Yes | 0.634478 | 1683 | 0.211631 | 0.624367 | 0.644589 |
| 3 | No | 0.330051 | 6602 | 0.165981 | 0.326047 | 0.334054 |
|  | Yes | 0.577669 | 917 | 0.187817 | 0.565513 | 0.589826 |
| 4 | No | 0.386335 | 233 | 0.159168 | 0.365897 | 0.406773 |
|  | Yes | 0.647114 | 23 | 0.171101 | 0.577187 | 0.717040 |

In [61]:

```
plots.tukey_subsets(svea_df, "pred_mean", "naturalness", "region")
```

```
region: 1

  group1 group2      Diff     Lower     Upper     q-value  p-value
0    Yes     No  0.352235  0.347757  0.356713  218.032878    0.001


region: 2

  group1 group2      Diff     Lower     Upper    q-value  p-value
0     No    Yes  0.347495  0.337178  0.357811  93.379863    0.001


region: 3

  group1 group2      Diff     Lower     Upper    q-value  p-value
0     No    Yes  0.247793  0.235994  0.259591  58.223501    0.001


region: 4

  group1 group2      Diff     Lower     Upper    q-value  p-value
0     No    Yes  0.266018  0.195276  0.336759  10.477331    0.001
```

#### NFI validation¶

plot level validation (1 central pixel + 4 neighbours)

In [62]:

```
# load a dataframe with predictions and NFI atributes
nfi_df = pandas.read_csv("nfi_predictions.csv")

# at least 4 pixels needed
nfi_df = nfi_df[nfi_df.pred_count > 3]

nfi_df["region"] = pandas.Categorical(nfi_df.region)
nfi_df["naturalness"] = pandas.Categorical(nfi_df.naturalness, categories=[3, 1, 2])
nfi_df.naturalness.cat.rename_categories({
    1.0: "normal", 2.0: "natural", 3.0: "plantation"
}, inplace=True)

# drop rows with NA in natura2000 column
nfi_df.dropna(subset=["natura2000"], inplace=True)
nfi_df.natura2000.replace({0: "No", 1: "Yes"}, inplace=True)
nfi_df["natura2000"] = pandas.Categorical(nfi_df.natura2000)

# show the number of validation polygons per region
display(nfi_df.region.value_counts())
print("TOTAL PLOTS N: ", nfi_df.region.value_counts().sum())
```

```
2    5397
3    4623
1    3072
4     683
Name: region, dtype: int64
```

```
TOTAL PLOTS N:  13775
```

##### Boxplot -> forest naturalness classes¶

1=normal; 2=natural forest; 3=plantation forest

In [63]:

```
plots.plot_boxplots(
    nfi_df, "naturalness", height=4, aspect=0.75, fig_title="c) Plot-level validation - Forest naturalness",
    save_name="Figure4c.png", width=0.85
)
```

In [64]:

```
plots.describe_ci(nfi_df, "pred_mean", ["region", "naturalness"])
```

Out[64]:

|  |  | mean | count | std | ci95\_lo | ci95\_hi |
| --- | --- | --- | --- | --- | --- | --- |
| region | naturalness |  |  |  |  |  |
| 1 | plantation | 0.200842 | 19 | 0.117059 | 0.148206 | 0.253478 |
|  | normal | 0.360762 | 2897 | 0.249938 | 0.351661 | 0.369864 |
|  | natural | 0.840240 | 156 | 0.176087 | 0.812607 | 0.867872 |
| 2 | plantation | 0.240282 | 74 | 0.141615 | 0.208016 | 0.272549 |
|  | normal | 0.372475 | 5200 | 0.230861 | 0.366200 | 0.378750 |
|  | natural | 0.760741 | 123 | 0.230626 | 0.719984 | 0.801499 |
| 3 | plantation | 0.272588 | 103 | 0.185641 | 0.236737 | 0.308440 |
|  | normal | 0.375125 | 4494 | 0.207887 | 0.369047 | 0.381203 |
|  | natural | 0.630788 | 26 | 0.206469 | 0.551425 | 0.710152 |
| 4 | plantation | 0.386862 | 69 | 0.173586 | 0.345904 | 0.427821 |
|  | normal | 0.419328 | 612 | 0.202014 | 0.403323 | 0.435333 |
|  | natural | 0.422600 | 2 | 0.298965 | 0.008256 | 0.836944 |

In [65]:

```
plots.describe_ci(nfi_df, "pred_mean", ["region", "naturalness"])["count"].sum()
```

Out[65]:

```
13775
```

In [66]:

```
plots.tukey_subsets(nfi_df, "pred_mean", "naturalness", "region")
```

```
region: 1

    group1      group2      Diff     Lower     Upper    q-value   p-value
0   normal     natural  0.479477  0.432036  0.526919  33.515176  0.001000
1   normal  plantation  0.159920  0.027067  0.292774   3.991732  0.013303
2  natural  plantation  0.639398  0.499145  0.779650  15.117899  0.001000


region: 2

    group1      group2      Diff     Lower     Upper    q-value  p-value
0   normal     natural  0.388266  0.339102  0.437430  26.183086    0.001
1   normal  plantation  0.132193  0.069100  0.195285   6.946571    0.001
2  natural  plantation  0.520459  0.441174  0.599744  21.763918    0.001


region: 3

       group1      group2      Diff     Lower     Upper    q-value  p-value
0      normal  plantation  0.102537  0.054076  0.150997   7.015430    0.001
1      normal     natural  0.255663  0.160022  0.351305   8.862987    0.001
2  plantation     natural  0.358200  0.251474  0.464927  11.127909    0.001


region: 4

    group1      group2      Diff     Lower     Upper   q-value   p-value
0  natural      normal  0.003272 -0.328680  0.335225  0.032745  0.900000
1  natural  plantation  0.035738 -0.300442  0.371918  0.353132  0.900000
2   normal  plantation  0.032465 -0.027054  0.091984  1.811960  0.407916
```

In [67]:

```
plots.plot_boxplots(
    nfi_df, "natura2000", height=4, aspect=0.75, fig_title="d) Plot-level validation - Natura 2000 habitat quality",
    save_name="Figure4d.png", width=0.5
)
```

In [68]:

```
plots.describe_ci(nfi_df, "pred_mean", ["region", "natura2000"])
```

Out[68]:

|  |  | mean | count | std | ci95\_lo | ci95\_hi |
| --- | --- | --- | --- | --- | --- | --- |
| region | natura2000 |  |  |  |  |  |
| 1 | No | 0.347721 | 2841 | 0.235795 | 0.339050 | 0.356392 |
|  | Yes | 0.831806 | 231 | 0.234736 | 0.801535 | 0.862077 |
| 2 | No | 0.363788 | 5216 | 0.223105 | 0.357733 | 0.369843 |
|  | Yes | 0.832618 | 181 | 0.191194 | 0.804764 | 0.860472 |
| 3 | No | 0.366985 | 4528 | 0.203106 | 0.361069 | 0.372901 |
|  | Yes | 0.721889 | 95 | 0.180615 | 0.685569 | 0.758210 |
| 4 | No | 0.406601 | 660 | 0.194696 | 0.391747 | 0.421455 |
|  | Yes | 0.687417 | 23 | 0.135743 | 0.631941 | 0.742894 |

In [69]:

```
plots.tukey_subsets(nfi_df, "pred_mean", "natura2000", "region")
```

```
region: 1

  group1 group2      Diff     Lower     Upper    q-value  p-value
0     No    Yes  0.484085  0.452464  0.515706  42.450157    0.001


region: 2

  group1 group2     Diff     Lower     Upper    q-value  p-value
0     No    Yes  0.46883  0.435908  0.501752  39.480747    0.001


region: 3

  group1 group2      Diff     Lower     Upper    q-value  p-value
0     No    Yes  0.354904  0.313713  0.396095  23.888206    0.001


region: 4

  group1 group2      Diff     Lower     Upper  q-value  p-value
0     No    Yes  0.280816  0.200405  0.361228  9.69706    0.001
```

### Hyperparameters tuning¶

In [17]:

```
# performance metrics after the hyperparameter tuning procedure using Bayesian optimization with Gaussian processes
# implemented in the scikit-optimize Python package

"""
# Search space for hyperparameters tuning has been defined as follows:

search_space = list()
search_space.append(Categorical(["gini", "entropy", "log_loss"], name="criterion"))
search_space.append(Integer(3, 30, name="max_depth"))
search_space.append(Real(0.0001, 0.3, name="min_samples_split"))
search_space.append(Real(0.0001, 0.3, name="min_samples_leaf"))
search_space.append(Real(0.1, 1.0, name="max_features"))
"""

results = plots.load_models("./", r".*tuned\.pickle$")[0]

# get a table with metrics and associated best hyperparameters
metrics_sel = ["accuracy", "roc_auc", "pr_auc", "pearson_r", "briers_score", "MCC"]
columns = ["region"] + metrics_sel + results[0]["gp_results"].space.dimension_names
df_report = []

for key, result in results.items():
    values = [key + 1]
    metrics = result["calls_scv"][-1][metrics_sel].mean().tolist()
    hparams = result["gp_results"].x_iters[-1]
    values.extend(metrics + hparams)
    df_report.append(values)

df_report = pandas.DataFrame(df_report, columns=columns)
display(df_report)
```

```
Loading S1_selected_models_tuned.pickle...
```

|  | region | accuracy | roc\_auc | pr\_auc | pearson\_r | briers\_score | MCC | criterion | max\_depth | min\_samples\_split | min\_samples\_leaf | max\_features |
| --- | --- | --- | --- | --- | --- | --- | --- | --- | --- | --- | --- | --- |
| 0 | 1 | 0.811888 | 0.891874 | 0.883525 | 0.680121 | 0.135028 | 0.621279 | entropy | 28 | 0.0001 | 0.0001 | 1.000000 |
| 1 | 2 | 0.809701 | 0.895349 | 0.868471 | 0.682789 | 0.133356 | 0.611816 | entropy | 27 | 0.0001 | 0.0001 | 0.420203 |
| 2 | 3 | 0.812218 | 0.892466 | 0.839481 | 0.669244 | 0.138165 | 0.616012 | log\_loss | 24 | 0.0001 | 0.0001 | 0.426681 |
| 3 | 4 | 0.798050 | 0.888916 | 0.840164 | 0.661486 | 0.152758 | 0.584113 | log\_loss | 30 | 0.0001 | 0.0001 | 0.100000 |

In [18]:

```
# performance metrics (default hyperparameters)
means, stds = plots.get_metrics(models_stats)
display(means)
display(stds)
```

|  | accuracy | TSS | roc\_auc | pr\_auc | pearson\_r | briers\_score | MCC |
| --- | --- | --- | --- | --- | --- | --- | --- |
| 0 | 0.811071 | 0.607707 | 0.894391 | 0.890909 | 0.682667 | 0.137704 | 0.621510 |
| 1 | 0.812354 | 0.608865 | 0.897814 | 0.871556 | 0.686108 | 0.135682 | 0.617628 |
| 2 | 0.808228 | 0.591838 | 0.893231 | 0.841799 | 0.666823 | 0.141793 | 0.609604 |
| 3 | 0.806138 | 0.567882 | 0.892063 | 0.849692 | 0.658849 | 0.152614 | 0.611538 |

|  | accuracy | TSS | roc\_auc | pr\_auc | pearson\_r | briers\_score | MCC |
| --- | --- | --- | --- | --- | --- | --- | --- |
| 0 | 0.019681 | 0.045446 | 0.014750 | 0.019952 | 0.026054 | 0.007905 | 0.039076 |
| 1 | 0.020558 | 0.046115 | 0.018678 | 0.038806 | 0.036213 | 0.008786 | 0.046483 |
| 2 | 0.019474 | 0.064574 | 0.015308 | 0.024282 | 0.028949 | 0.009160 | 0.036099 |
| 3 | 0.051995 | 0.164805 | 0.032248 | 0.054130 | 0.056224 | 0.017430 | 0.090672 |

In [ ]:
