## Supplementary material for "Mapping forests with different levels of naturalness using machine learning and landscape data mining": Jupyter Notebooks: M2_global.html

validation


In [1]:

```
%load_ext autoreload
%autoreload 2

import warnings
warnings.filterwarnings("ignore")

from matplotlib import pyplot as plt
import matplotlib as mpl
mpl.rcParams["figure.dpi"] = 150

import json
import sys
import numpy as np
import pandas
import seaborn as sns
sns.set_style("white")

sys.path.insert(0, "/app/src")

from plots import plots
from config import settings
```

In [2]:

```
with open("./config.json", "r") as _file:
    mconfig = json.load(_file)
print(json.dumps(mconfig, indent=2))

model_name = mconfig["model_metadata"]["name"]
target_var = mconfig["model_specification"]["target_variable"]
region_var = mconfig["model_specification"]["stratification_layer"]
```

```
{
  "model_metadata": {
    "name": "S1_global",
    "description": "Full global model using all available variables expressed at all target spatial scales. One global model fitted to all ecoregions."
  },
  "model_data": {
    "samples_layer": "SAMPLES_1"
  },
  "model_specification": {
    "target_variable": "HCVF",
    "stratification_layer": "ecoregions",
    "stratification_submodels": [
      {
        "regions": [
          1,
          2,
          3,
          4
        ],
        "variables_list": []
      }
    ],
    "lat_field": "LAT",
    "lon_field": "LON",
    "target_epsg": 3006,
    "variables_list": [],
    "variables_drop_regexp": "",
    "variables_keep_regexp": "",
    "variables_latlon": true,
    "balanced_rf": true
  },
  "model_crossvalidation": {
    "n_splits": 10,
    "spatial": true,
    "spatial_grid": "grid_20.geojson"
  },
  "model_hyperparameters": {
    "n_estimators": 500,
    "oob_score": true
  },
  "prediction": {
    "predict": true
  }
}
```

In [3]:

```
models = plots.load_models("./", regexp=fr"{model_name}_R[1-5]*.pickle")
models_stats = plots.get_cv_validation_stats(models)
```

```
Loading S1_global_R1234.pickle...
```

In [4]:

```
print(models[0].data.shape)
models_labels=["North boreal", "South boreal", "Hemiboreal", "Nemoral"]

g = sns.FacetGrid(
    data=models[0].data, col=region_var, aspect=0.7, 
    sharey=False
)
g.map_dataframe(sns.countplot, x=target_var, palette=["black","red"])
axes = g.axes.flatten()
for i, ax in enumerate(axes):
    ax.set_title(models_labels[i], fontweight="bold", fontsize=16)
```

```
(14090, 130)
```

In [5]:

```
plots.plot_validation_curves_multimodel(
    models_stats, colors="bgrcmykw", figsize=(10, 5), plot_std=True, std_alpha=0.1,
    models_labels=["North boreal", "South boreal", "Hemiboreal", "Nemoral"],
    save_name="Figure2.png"
)
```

In [6]:

```
# performance metrics
means, stds = plots.get_metrics(models_stats)
display(means)
display(stds)
```

|  | accuracy | TSS | roc\_auc | pr\_auc | pearson\_r | briers\_score | MCC |
| --- | --- | --- | --- | --- | --- | --- | --- |
| 0 | 0.814932 | 0.631365 | 0.896997 | 0.872892 | 0.68359 | 0.136406 | 0.627011 |

|  | accuracy | TSS | roc\_auc | pr\_auc | pearson\_r | briers\_score | MCC |
| --- | --- | --- | --- | --- | --- | --- | --- |
| 0 | 0.010898 | 0.017799 | 0.00792 | 0.021635 | 0.015701 | 0.004563 | 0.020601 |

In [7]:

```
# number of variables per model
for i, model in enumerate(models):
    print(f"Model_{i+1}: ", len(model.cov_names))
```

```
Model_1:  128
```

In [8]:

```
# plot features Gini's importances
for i, model in enumerate(models):
    print(models_labels[i])
    _plot = plots.plot_gini_importances(model, figsize=(14, 10), n=50)
```

```
North boreal
```

##### Partial Dependence Plot¶

In [11]:

```
plots.plot_partial_dependences(models, n_features=6, figsize=(8, 2))
```

##### Accumulated Local Effects Plot¶

In [13]:

```
plots.plot_ales(models, n_features=6, xlims=(-2, 2), figsize=(8, 2))
```

In [14]:

```
# average max tree depth per model
import numpy as np
for i, model in enumerate(models):
    max_depths = []
    for tree_idx, est in enumerate(model.model.estimators_):
        max_depths.append(est.tree_.max_depth)
    print(f"{models_labels[i]}: ", np.array(max_depths).mean())
```

```
North boreal:  26.68
```

### Validation¶

#### Sveaskog validation¶

tree stand (polygon) level validation; minimum 10 pixels with predicted value

In [15]:

```
# load a dataframe with predictions and Sveaskog atributes
svea_df = pandas.read_csv("svea_predictions.csv")
svea_df = plots.svea_preprocess(svea_df)

# filter out data not reliable for validation
# min number of 1 ha pixels to calculate the mean probability
svea_df = svea_df[svea_df.pred_count > 10]

# show the number of validation polygons per region
display(svea_df.region.value_counts())
print("TOTAL POLYGONS N: ", svea_df.region.value_counts().sum())
```

```
1    41784
2     7989
3     7519
4      256
Name: region, dtype: int64
```

```
TOTAL POLYGONS N:  57548
```

##### Boxplot -> forest management classes¶

"NF" (conservation, no management)
"NF\_NM" (conservation, no yet specified)
"NM" (conservation, management)
"PE" (production forest, increased conservation considerations)
"PG" (production forest, general conservation conciderations)

Management categories for stands. A goal for long-term management is set for each stand, in one of four categories: (i) NF = Nature conservation, free development; (ii) NM = Nature conservation-oriented management, often implying restoration measures; (iii) PG = Production with general conservation concern (green-tree and deadwood retention for biodiversity at harvest, represented by black dots). The minimum level of retention is specified, and always >15 % of the harvested area; (iv) PE = Production with enhanced conservation concern.

In [16]:

```
plots.plot_boxplots(
    svea_df, "management", height=4, aspect=0.75, fig_title="a) Stand-level validation - Forest management objectives",
    save_name="Figure4a.png", width=0.85
)
```

In [17]:

```
plots.describe_ci(svea_df, "pred_mean", ["region", "management"])
```

Out[17]:

|  |  | mean | count | std | ci95\_lo | ci95\_hi |
| --- | --- | --- | --- | --- | --- | --- |
| region | management |  |  |  |  |  |
| 1 | NF | 0.652419 | 4148 | 0.208494 | 0.646074 | 0.658764 |
|  | NF\_NM | 0.736538 | 2401 | 0.169667 | 0.729752 | 0.743325 |
|  | NM | 0.592179 | 903 | 0.200799 | 0.579081 | 0.605276 |
|  | PE | 0.425892 | 2516 | 0.198216 | 0.418147 | 0.433637 |
|  | PG | 0.301942 | 31816 | 0.174297 | 0.300027 | 0.303858 |
| 2 | NF | 0.668832 | 1245 | 0.187604 | 0.658411 | 0.679253 |
|  | NF\_NM | 0.709016 | 82 | 0.176820 | 0.670744 | 0.747288 |
|  | NM | 0.529012 | 253 | 0.198286 | 0.504578 | 0.553446 |
|  | PE | 0.393918 | 868 | 0.199898 | 0.380620 | 0.407217 |
|  | PG | 0.277086 | 5541 | 0.172672 | 0.272539 | 0.281632 |
| 3 | NF | 0.548057 | 297 | 0.186949 | 0.526795 | 0.569319 |
|  | NF\_NM | 0.599652 | 172 | 0.173364 | 0.573743 | 0.625561 |
|  | NM | 0.496324 | 303 | 0.199782 | 0.473828 | 0.518819 |
|  | PE | 0.309776 | 608 | 0.172668 | 0.296051 | 0.323501 |
|  | PG | 0.201473 | 6139 | 0.130012 | 0.198221 | 0.204725 |
| 4 | NF | 0.730533 | 5 | 0.096658 | 0.645809 | 0.815258 |
|  | NF\_NM | NaN | 0 | NaN | NaN | NaN |
|  | NM | 0.562205 | 19 | 0.177748 | 0.482279 | 0.642130 |
|  | PE | 0.553246 | 13 | 0.210772 | 0.438669 | 0.667823 |
|  | PG | 0.261058 | 219 | 0.158504 | 0.240065 | 0.282051 |

In [18]:

```
plots.tukey_subsets(svea_df, "pred_mean", "management", "region")
```

```
region: 1

  group1 group2      Diff     Lower     Upper     q-value  p-value
0     NF  NF_NM  0.083813  0.071092  0.096534   25.417063    0.001
1     NF     PG  0.351712  0.343517  0.359908  165.558013    0.001
2     NF     PE  0.204877  0.191155  0.218599   57.599477    0.001
3     NF     NM  0.061928  0.043680  0.080175   13.092237    0.001
4  NF_NM     PG  0.435525  0.425011  0.446039  159.805172    0.001
5  NF_NM     PE  0.288690  0.273469  0.303911   73.171342    0.001
6  NF_NM     NM  0.145741  0.126341  0.165140   28.981485    0.001
7     PG     PE  0.146835  0.135130  0.158540   48.395116    0.001
8     PG     NM  0.289784  0.273001  0.306568   66.607719    0.001
9     PE     NM  0.142949  0.122879  0.163020   27.476829    0.001


region: 2

  group1 group2      Diff     Lower     Upper    q-value   p-value
0     PG     PE  0.120151  0.099718  0.140584  22.689684  0.001000
1     PG     NF  0.395885  0.380269  0.411501  97.818639  0.001000
2     PG     NM  0.256770  0.224864  0.288675  31.053180  0.001000
3     PG  NF_NM  0.436406  0.381345  0.491466  30.582609  0.001000
4     PE     NF  0.275734  0.251899  0.299569  44.637494  0.001000
5     PE     NM  0.136619  0.099983  0.173255  14.388887  0.001000
6     PE  NF_NM  0.316255  0.258324  0.374185  21.064751  0.001000
7     NF     NM  0.139115  0.104931  0.173300  15.702697  0.001000
8     NF  NF_NM  0.040520 -0.015891  0.096932   2.771598  0.286101
9     NM  NF_NM  0.179636  0.116736  0.242536  11.019702  0.001000


region: 3

  group1 group2      Diff     Lower     Upper    q-value  p-value
0     PE     NF  0.237738  0.210043  0.265433  33.122582    0.001
1     PE     PG  0.110208  0.093108  0.127307  24.869069    0.001
2     PE  NF_NM  0.291920  0.258155  0.325684  33.360385    0.001
3     PE     NM  0.184732  0.156914  0.212549  25.623916    0.001
4     NF     PG  0.347946  0.325054  0.370837  58.649297    0.001
5     NF  NF_NM  0.054182  0.017146  0.091218   5.644895    0.001
6     NF     NM  0.053006  0.021297  0.084716   6.450173    0.001
7     PG  NF_NM  0.402128  0.372177  0.432079  51.806335    0.001
8     PG     NM  0.294939  0.271899  0.317980  49.394381    0.001
9  NF_NM     NM  0.107188  0.070060  0.144316  11.139672    0.001


region: 4

  group1 group2      Diff     Lower     Upper    q-value   p-value
0     PG     NM  0.310251  0.209767  0.410736  11.299803  0.001000
1     PG     NF  0.492240  0.280801  0.703679   8.520197  0.001000
2     PG     PE  0.269243  0.139592  0.398894   7.600239  0.001000
3     NM     NF  0.181988 -0.048375  0.412352   2.891262  0.174999
4     NM     PE  0.041009 -0.117643  0.199660   0.945991  0.900000
5     NF     PE  0.222997 -0.021501  0.467495   3.337955  0.087675
```

##### Boxplot -> forest naturalness classes¶

In [19]:

```
plots.plot_boxplots(
    svea_df, "naturalness", height=4, aspect=0.75, fig_title="b) Stand-level validation - Forest naturalness",
    save_name="Figure4b.png", width=0.5
)
```

In [22]:

```
plots.describe_ci(svea_df, "pred_mean", ["region", "naturalness"])
```

Out[22]:

|  |  | mean | count | std | ci95\_lo | ci95\_hi |
| --- | --- | --- | --- | --- | --- | --- |
| region | naturalness |  |  |  |  |  |
| 1 | No | 0.310156 | 34130 | 0.178419 | 0.308264 | 0.312049 |
|  | Yes | 0.666567 | 7654 | 0.205259 | 0.661969 | 0.671166 |
| 2 | No | 0.291303 | 6306 | 0.179833 | 0.286864 | 0.295742 |
|  | Yes | 0.632782 | 1683 | 0.207190 | 0.622883 | 0.642681 |
| 3 | No | 0.209552 | 6602 | 0.137390 | 0.206237 | 0.212866 |
|  | Yes | 0.499483 | 917 | 0.207228 | 0.486070 | 0.512896 |
| 4 | No | 0.279994 | 233 | 0.178750 | 0.257041 | 0.302946 |
|  | Yes | 0.585215 | 23 | 0.170289 | 0.515620 | 0.654810 |

In [23]:

```
plots.tukey_subsets(svea_df, "pred_mean", "naturalness", "region")
```

```
region: 1

  group1 group2      Diff     Lower     Upper     q-value  p-value
0    Yes     No  0.358113  0.353484  0.362743  214.413017    0.001


region: 2

  group1 group2      Diff     Lower     Upper    q-value  p-value
0     No    Yes  0.348786  0.338577  0.358995  94.714515    0.001


region: 3

  group1 group2      Diff    Lower     Upper    q-value  p-value
0     No    Yes  0.291827  0.28153  0.302124  78.566672    0.001


region: 4

  group1 group2      Diff     Lower     Upper    q-value  p-value
0     No    Yes  0.312391  0.234873  0.389908  11.228277    0.001
```

#### NFI validation¶

plot level validation (1 central pixel + 4 neighbours)

In [24]:

```
# load a dataframe with predictions and NFI atributes
nfi_df = pandas.read_csv("nfi_predictions.csv")

# at least 4 pixels needed
nfi_df = nfi_df[nfi_df.pred_count > 3]

nfi_df["region"] = pandas.Categorical(nfi_df.region)
nfi_df["naturalness"] = pandas.Categorical(nfi_df.naturalness, categories=[3, 1, 2])
nfi_df.naturalness.cat.rename_categories({
    1.0: "normal", 2.0: "natural", 3.0: "plantation"
}, inplace=True)

# drop rows with NA in natura2000 column
nfi_df.dropna(subset=["natura2000"], inplace=True)
nfi_df.natura2000.replace({0: "No", 1: "Yes"}, inplace=True)
nfi_df["natura2000"] = pandas.Categorical(nfi_df.natura2000)

# show the number of validation polygons per region
display(nfi_df.region.value_counts())
print("TOTAL PLOTS N: ", nfi_df.region.value_counts().sum())
```

```
2    5397
3    4623
1    3072
4     683
Name: region, dtype: int64
```

```
TOTAL PLOTS N:  13775
```

##### Boxplot -> forest naturalness classes¶

1=normal; 2=natural forest; 3=plantation forest

In [25]:

```
plots.plot_boxplots(
    nfi_df, "naturalness", height=4, aspect=0.75, fig_title="c) Plot-level validation - Forest naturalness",
    save_name="Figure4c.png", width=0.85
)
```

In [26]:

```
plots.describe_ci(nfi_df, "pred_mean", ["region", "naturalness"])
```

Out[26]:

|  |  | mean | count | std | ci95\_lo | ci95\_hi |
| --- | --- | --- | --- | --- | --- | --- |
| region | naturalness |  |  |  |  |  |
| 1 | plantation | 0.209521 | 19 | 0.122561 | 0.154411 | 0.264631 |
|  | normal | 0.391918 | 2897 | 0.256867 | 0.382564 | 0.401272 |
|  | natural | 0.868117 | 156 | 0.174687 | 0.840704 | 0.895529 |
| 2 | plantation | 0.241836 | 74 | 0.164380 | 0.204383 | 0.279290 |
|  | normal | 0.372588 | 5200 | 0.241598 | 0.366021 | 0.379155 |
|  | natural | 0.765439 | 123 | 0.237271 | 0.723507 | 0.807371 |
| 3 | plantation | 0.215325 | 103 | 0.176856 | 0.181170 | 0.249481 |
|  | normal | 0.302917 | 4494 | 0.198063 | 0.297126 | 0.308708 |
|  | natural | 0.558204 | 26 | 0.215528 | 0.475358 | 0.641050 |
| 4 | plantation | 0.293449 | 69 | 0.191339 | 0.248302 | 0.338597 |
|  | normal | 0.360155 | 612 | 0.225793 | 0.342266 | 0.378044 |
|  | natural | 0.430200 | 2 | 0.472064 | -0.224048 | 1.084448 |

In [29]:

```
plots.describe_ci(nfi_df, "pred_mean", ["region", "naturalness"])["count"].sum()
```

Out[29]:

```
13775
```

In [30]:

```
plots.tukey_subsets(nfi_df, "pred_mean", "naturalness", "region")
```

```
region: 1

    group1      group2      Diff     Lower     Upper    q-value   p-value
0   normal     natural  0.476198  0.427483  0.524913  32.415810  0.001000
1   normal  plantation  0.182397  0.045977  0.318817   4.433751  0.004934
2  natural  plantation  0.658596  0.514578  0.802613  15.164731  0.001000


region: 2

    group1      group2      Diff     Lower     Upper    q-value  p-value
0   normal     natural  0.392851  0.341390  0.444312  25.309639    0.001
1   normal  plantation  0.130752  0.064711  0.196792   6.564126    0.001
2  natural  plantation  0.523603  0.440613  0.606592  20.917957    0.001


region: 3

       group1      group2      Diff     Lower     Upper    q-value  p-value
0      normal  plantation  0.087592  0.041397  0.133787   6.286811    0.001
1      normal     natural  0.255286  0.164115  0.346458   9.283876    0.001
2  plantation     natural  0.342879  0.241141  0.444616  11.174248    0.001


region: 4

    group1      group2      Diff     Lower     Upper   q-value   p-value
0  natural      normal  0.070045 -0.301206  0.441296  0.626749  0.890327
1  natural  plantation  0.136751 -0.239228  0.512730  1.208227  0.655090
2   normal  plantation  0.066705  0.000140  0.133271  3.328867  0.049384
```

In [31]:

```
plots.plot_boxplots(
    nfi_df, "natura2000", height=4, aspect=0.75, fig_title="d) Plot-level validation - Natura 2000 habitat quality",
    save_name="Figure4d.png", width=0.5
)
```

In [32]:

```
plots.describe_ci(nfi_df, "pred_mean", ["region", "natura2000"])
```

Out[32]:

|  |  | mean | count | std | ci95\_lo | ci95\_hi |
| --- | --- | --- | --- | --- | --- | --- |
| region | natura2000 |  |  |  |  |  |
| 1 | No | 0.379802 | 2841 | 0.244509 | 0.370811 | 0.388794 |
|  | Yes | 0.847513 | 231 | 0.246147 | 0.815770 | 0.879256 |
| 2 | No | 0.363628 | 5216 | 0.234127 | 0.357274 | 0.369981 |
|  | Yes | 0.844320 | 181 | 0.182455 | 0.817739 | 0.870901 |
| 3 | No | 0.295026 | 4528 | 0.192709 | 0.289412 | 0.300639 |
|  | Yes | 0.653967 | 95 | 0.178182 | 0.618136 | 0.689798 |
| 4 | No | 0.343582 | 660 | 0.219800 | 0.326813 | 0.360351 |
|  | Yes | 0.641700 | 23 | 0.123856 | 0.591082 | 0.692318 |

In [33]:

```
plots.tukey_subsets(nfi_df, "pred_mean", "natura2000", "region")
```

```
region: 1

  group1 group2      Diff     Lower     Upper    q-value  p-value
0     No    Yes  0.467711  0.434893  0.500528  39.519375    0.001


region: 2

  group1 group2      Diff     Lower     Upper    q-value  p-value
0     No    Yes  0.480692  0.446218  0.515167  38.656876    0.001


region: 3

  group1 group2      Diff     Lower    Upper    q-value  p-value
0     No    Yes  0.358942  0.319833  0.39805  25.446673    0.001


region: 4

  group1 group2      Diff    Lower     Upper   q-value  p-value
0     No    Yes  0.298118  0.20759  0.388646  9.144128    0.001
```

In [ ]:
