## Supplementary material for "Mapping forests with different levels of naturalness using machine learning and landscape data mining": Jupyter Notebooks: M3a_full.html

validation


In [2]:

```
%load_ext autoreload
%autoreload 2

import warnings
warnings.filterwarnings("ignore")

from matplotlib import pyplot as plt
import matplotlib as mpl
mpl.rcParams["figure.dpi"] = 150

import json
import sys
import numpy as np
import pandas
import seaborn as sns
sns.set_style("white")

sys.path.insert(0, "/app/src")

from plots import plots
from config import settings
```

In [5]:

```
with open("./config.json", "r") as _file:
    mconfig = json.load(_file)
print(json.dumps(mconfig, indent=2))

model_name = mconfig["model_metadata"]["name"]
target_var = mconfig["model_specification"]["target_variable"]
region_var = mconfig["model_specification"]["stratification_layer"]
```

```
{
  "model_metadata": {
    "name": "S1_full",
    "description": "Full model using all available variables expressed at all target spatial scales. Seperate model for each ecoregion is fitted."
  },
  "model_data": {
    "samples_layer": "SAMPLES_1"
  },
  "model_specification": {
    "target_variable": "HCVF",
    "stratification_layer": "ecoregions",
    "stratification_submodels": [
      {
        "regions": [
          1
        ],
        "variables_list": []
      },
      {
        "regions": [
          2
        ],
        "variables_list": []
      },
      {
        "regions": [
          3
        ],
        "variables_list": []
      },
      {
        "regions": [
          4
        ],
        "variables_list": []
      }
    ],
    "lat_field": "LAT",
    "lon_field": "LON",
    "target_epsg": 3006,
    "variables_list": [],
    "variables_drop_regexp": "",
    "variables_keep_regexp": "",
    "variables_latlon": true,
    "balanced_rf": true
  },
  "model_crossvalidation": {
    "n_splits": 10,
    "spatial": true,
    "spatial_grid": "grid_20.geojson"
  },
  "model_hyperparameters": {
    "n_estimators": 500,
    "oob_score": true
  },
  "prediction": {
    "predict": true
  }
}
```

In [6]:

```
models = plots.load_models("./", regexp=fr"{model_name}_R[1-5]*.pickle")
models_stats = plots.get_cv_validation_stats(models)
```

```
Loading S1_full_R1.pickle...
Loading S1_full_R2.pickle...
Loading S1_full_R3.pickle...
Loading S1_full_R4.pickle...
```

In [7]:

```
models[0].model.get_params()
```

Out[7]:

```
{'bootstrap': True,
 'ccp_alpha': 0.0,
 'class_weight': None,
 'criterion': 'gini',
 'max_depth': None,
 'max_features': 'sqrt',
 'max_leaf_nodes': None,
 'max_samples': None,
 'min_impurity_decrease': 0.0,
 'min_samples_leaf': 1,
 'min_samples_split': 2,
 'min_weight_fraction_leaf': 0.0,
 'n_estimators': 500,
 'n_jobs': None,
 'oob_score': True,
 'random_state': None,
 'replacement': False,
 'sampling_strategy': 'auto',
 'verbose': 0,
 'warm_start': False}
```

In [7]:

```
mconfig["model_hyperparameters"]
```

Out[7]:

```
{'n_estimators': 500, 'oob_score': True}
```

In [4]:

```
print(models[0].data.shape)
models_labels=["North boreal", "South boreal", "Hemiboreal", "Nemoral"]

g = sns.FacetGrid(
    data=models[0].data, col=region_var, aspect=0.7, 
    sharey=False
)
g.map_dataframe(sns.countplot, x=target_var, palette=["black","red"])
axes = g.axes.flatten()
for i, ax in enumerate(axes):
    ax.set_title(models_labels[i], fontweight="bold", fontsize=16)
```

```
(14090, 130)
```

In [5]:

```
plots.plot_validation_curves_multimodel(
    models_stats, colors="bgrcmykw", figsize=(10, 5), plot_std=True, std_alpha=0.1,
    models_labels=["North boreal", "South boreal", "Hemiboreal", "Nemoral"],
    save_name="Figure2.png"
)
```

In [6]:

```
# performance metrics
means, stds = plots.get_metrics(models_stats)
display(means)
display(stds)
```

|  | accuracy | TSS | roc\_auc | pr\_auc | pearson\_r | briers\_score | MCC |
| --- | --- | --- | --- | --- | --- | --- | --- |
| 0 | 0.814602 | 0.622555 | 0.895849 | 0.889916 | 0.685549 | 0.135498 | 0.627630 |
| 1 | 0.816158 | 0.619318 | 0.899882 | 0.878063 | 0.690780 | 0.133521 | 0.626025 |
| 2 | 0.806799 | 0.603023 | 0.894206 | 0.844005 | 0.666070 | 0.141412 | 0.609277 |
| 3 | 0.815561 | 0.499055 | 0.909073 | 0.880873 | 0.696555 | 0.139441 | 0.635410 |

|  | accuracy | TSS | roc\_auc | pr\_auc | pearson\_r | briers\_score | MCC |
| --- | --- | --- | --- | --- | --- | --- | --- |
| 0 | 0.027193 | 0.046479 | 0.022523 | 0.038950 | 0.040787 | 0.011069 | 0.056361 |
| 1 | 0.024461 | 0.029008 | 0.015541 | 0.027983 | 0.028191 | 0.008481 | 0.048373 |
| 2 | 0.023409 | 0.030500 | 0.020834 | 0.024094 | 0.034813 | 0.010400 | 0.045407 |
| 3 | 0.067950 | 0.134525 | 0.034206 | 0.052976 | 0.066607 | 0.021271 | 0.124319 |

In [7]:

```
# number of variables per model
for i, model in enumerate(models):
    print(f"Model_{i+1}: ", len(model.cov_names))
```

```
Model_1:  128
Model_2:  128
Model_3:  128
Model_4:  128
```

In [8]:

```
# plot features Gini's importances
for i, model in enumerate(models):
    print(models_labels[i])
    _plot = plots.plot_gini_importances(model, figsize=(14, 10), n=50)
```

```
North boreal
```

```
South boreal
```

```
Hemiboreal
```

```
Nemoral
```

##### Partial Dependence Plot¶

In [9]:

```
plots.plot_partial_dependences(models, n_features=6)
```

##### Accumulated Local Effects Plot¶

In [10]:

```
plots.plot_ales(models, n_features=6, xlims=(-2, 2))
```

In [11]:

```
# average max tree depth per model
import numpy as np
for i, model in enumerate(models):
    max_depths = []
    for tree_idx, est in enumerate(model.model.estimators_):
        max_depths.append(est.tree_.max_depth)
    print(f"{models_labels[i]}: ", np.array(max_depths).mean())
```

```
North boreal:  21.734
South boreal:  21.27
Hemiboreal:  20.0
Nemoral:  10.624
```

### Validation¶

#### Sveaskog validation¶

tree stand (polygon) level validation; minimum 10 pixels with predicted value

In [12]:

```
# load a dataframe with predictions and Sveaskog atributes
svea_df = pandas.read_csv("svea_predictions.csv")
svea_df = plots.svea_preprocess(svea_df)

# filter out data not reliable for validation
# min number of 1 ha pixels to calculate the mean probability
svea_df = svea_df[svea_df.pred_count > 10]

# show the number of validation polygons per region
display(svea_df.region.value_counts())
print("TOTAL POLYGONS N: ", svea_df.region.value_counts().sum())
```

```
1    41784
2     7989
3     7519
4      256
Name: region, dtype: int64
```

```
TOTAL POLYGONS N:  57548
```

##### Boxplot -> forest management classes¶

"NF" (conservation, no management)
"NF\_NM" (conservation, no yet specified)
"NM" (conservation, management)
"PE" (production forest, increased conservation considerations)
"PG" (production forest, general conservation conciderations)

Management categories for stands. A goal for long-term management is set for each stand, in one of four categories: (i) NF = Nature conservation, free development; (ii) NM = Nature conservation-oriented management, often implying restoration measures; (iii) PG = Production with general conservation concern (green-tree and deadwood retention for biodiversity at harvest, represented by black dots). The minimum level of retention is specified, and always >15 % of the harvested area; (iv) PE = Production with enhanced conservation concern.

In [13]:

```
plots.plot_boxplots(
    svea_df, "management", height=4, aspect=0.75, fig_title="a) Stand-level validation - Forest management objectives",
    save_name="Figure4a.png", width=0.85
)
```

In [14]:

```
plots.describe_ci(svea_df, "pred_mean", ["region", "management"])
```

Out[14]:

|  |  | mean | count | std | ci95\_lo | ci95\_hi |
| --- | --- | --- | --- | --- | --- | --- |
| region | management |  |  |  |  |  |
| 1 | NF | 0.633403 | 4148 | 0.220756 | 0.626685 | 0.640121 |
|  | NF\_NM | 0.719298 | 2401 | 0.179023 | 0.712137 | 0.726459 |
|  | NM | 0.567539 | 903 | 0.202613 | 0.554324 | 0.580755 |
|  | PE | 0.403016 | 2516 | 0.196755 | 0.395327 | 0.410704 |
|  | PG | 0.273840 | 31816 | 0.169551 | 0.271977 | 0.275703 |
| 2 | NF | 0.680336 | 1245 | 0.196830 | 0.669402 | 0.691270 |
|  | NF\_NM | 0.699105 | 82 | 0.181707 | 0.659775 | 0.738435 |
|  | NM | 0.533241 | 253 | 0.211276 | 0.507206 | 0.559275 |
|  | PE | 0.398226 | 868 | 0.211668 | 0.384144 | 0.412307 |
|  | PG | 0.275440 | 5541 | 0.185826 | 0.270547 | 0.280333 |
| 3 | NF | 0.601827 | 297 | 0.184480 | 0.580846 | 0.622808 |
|  | NF\_NM | 0.674658 | 172 | 0.160649 | 0.650649 | 0.698667 |
|  | NM | 0.582323 | 303 | 0.189865 | 0.560945 | 0.603702 |
|  | PE | 0.413626 | 608 | 0.180030 | 0.399316 | 0.427936 |
|  | PG | 0.299134 | 6139 | 0.161174 | 0.295102 | 0.303166 |
| 4 | NF | 0.744903 | 5 | 0.058850 | 0.693319 | 0.796487 |
|  | NF\_NM | NaN | 0 | NaN | NaN | NaN |
|  | NM | 0.625529 | 19 | 0.188044 | 0.540974 | 0.710084 |
|  | PE | 0.576362 | 13 | 0.168604 | 0.484707 | 0.668016 |
|  | PG | 0.326950 | 219 | 0.156015 | 0.306287 | 0.347613 |

In [15]:

```
plots.tukey_subsets(svea_df, "pred_mean", "management", "region")
```

```
region: 1

  group1 group2      Diff     Lower     Upper     q-value  p-value
0     NF  NF_NM  0.085634  0.073031  0.098237   26.212733    0.001
1     NF     PG  0.361283  0.353164  0.369403  171.657895    0.001
2     NF     PE  0.211585  0.197991  0.225179   60.043121    0.001
3     NF     NM  0.067749  0.049671  0.085828   14.457323    0.001
4  NF_NM     PG  0.446917  0.436501  0.457333  165.522727    0.001
5  NF_NM     PE  0.297219  0.282140  0.312298   76.039390    0.001
6  NF_NM     NM  0.153383  0.134163  0.172603   30.787248    0.001
7     PG     PE  0.149698  0.138102  0.161294   49.801389    0.001
8     PG     NM  0.293534  0.276906  0.310162   68.102132    0.001
9     PE     NM  0.143836  0.123952  0.163720   27.906399    0.001


region: 2

  group1 group2      Diff     Lower     Upper    q-value  p-value
0     PG     PE  0.122427  0.100664  0.144189  21.706877    0.001
1     PG     NF  0.410096  0.393464  0.426728  95.139106    0.001
2     PG     NM  0.263536  0.229554  0.297517  29.924239    0.001
3     PG  NF_NM  0.429199  0.370555  0.487843  28.239949    0.001
4     PE     NF  0.287669  0.262283  0.313056  43.724446    0.001
5     PE     NM  0.141109  0.102089  0.180129  13.953854    0.001
6     PE  NF_NM  0.306772  0.245072  0.368472  19.184785    0.001
7     NF     NM  0.146560  0.110151  0.182969  15.532307    0.001
8     NF  NF_NM  0.019103 -0.040980  0.079186   1.226806    0.900
9     NM  NF_NM  0.165663  0.098670  0.232656   9.541649    0.001


region: 3

  group1 group2      Diff     Lower     Upper    q-value   p-value
0     PE     NF  0.182229  0.149774  0.214684  21.665331  0.001000
1     PE     PG  0.120812  0.100774  0.140850  23.263681  0.001000
2     PE  NF_NM  0.257890  0.218322  0.297458  25.149182  0.001000
3     PE     NM  0.162724  0.130125  0.195323  19.260955  0.001000
4     NF     PG  0.303041  0.276215  0.329867  43.588754  0.001000
5     NF  NF_NM  0.075661  0.032259  0.119062   6.726593  0.001000
6     NF     NM  0.019505 -0.017654  0.056664   2.025414  0.592576
7     PG  NF_NM  0.378702  0.343603  0.413800  41.633052  0.001000
8     PG     NM  0.283536  0.256536  0.310536  40.520479  0.001000
9  NF_NM     NM  0.095166  0.051657  0.138675   8.439740  0.001000


region: 4

  group1 group2      Diff     Lower     Upper    q-value   p-value
0     PG     NM  0.306652  0.208478  0.404826  11.431579  0.001000
1     PG     NF  0.413989  0.207412  0.620565   7.334399  0.001000
2     PG     PE  0.242656  0.115987  0.369325   7.010962  0.001000
3     NM     NF  0.107337 -0.117730  0.332403   1.745399  0.593264
4     NM     PE  0.063996 -0.091007  0.218999   1.511013  0.686076
5     NF     PE  0.171332 -0.067544  0.410209   2.624970  0.250070
```

##### Boxplot -> forest naturalness classes¶

In [16]:

```
plots.plot_boxplots(
    svea_df, "naturalness", height=4, aspect=0.75, fig_title="b) Stand-level validation - Forest naturalness",
    save_name="Figure4b.png", width=0.5
)
```

In [17]:

```
plots.describe_ci(svea_df, "pred_mean", ["region", "naturalness"])
```

Out[17]:

|  |  | mean | count | std | ci95\_lo | ci95\_hi |
| --- | --- | --- | --- | --- | --- | --- |
| region | naturalness |  |  |  |  |  |
| 1 | No | 0.282339 | 34130 | 0.174194 | 0.280491 | 0.284187 |
|  | Yes | 0.647650 | 7654 | 0.214776 | 0.642838 | 0.652461 |
| 2 | No | 0.290504 | 6306 | 0.193006 | 0.285740 | 0.295267 |
|  | Yes | 0.641243 | 1683 | 0.217385 | 0.630857 | 0.651629 |
| 3 | No | 0.307351 | 6602 | 0.165934 | 0.303349 | 0.311354 |
|  | Yes | 0.577932 | 917 | 0.194790 | 0.565324 | 0.590539 |
| 4 | No | 0.342671 | 233 | 0.168353 | 0.321054 | 0.364289 |
|  | Yes | 0.646169 | 23 | 0.177978 | 0.573431 | 0.718906 |

In [18]:

```
plots.tukey_subsets(svea_df, "pred_mean", "naturalness", "region")
```

```
region: 1

  group1 group2      Diff     Lower     Upper     q-value  p-value
0    Yes     No  0.367673  0.363082  0.372264  221.992128    0.001


region: 2

  group1 group2      Diff     Lower     Upper   q-value  p-value
0     No    Yes  0.359667  0.348815  0.370519  91.88084    0.001


region: 3

  group1 group2      Diff     Lower     Upper    q-value  p-value
0     No    Yes  0.271418  0.259554  0.283282  63.421614    0.001


region: 4

  group1 group2      Diff    Lower     Upper    q-value  p-value
0     No    Yes  0.306138  0.23199  0.380286  11.503637    0.001
```

#### NFI validation¶

plot level validation (1 central pixel + 4 neighbours)

In [19]:

```
# load a dataframe with predictions and NFI atributes
nfi_df = pandas.read_csv("nfi_predictions.csv")

# at least 4 pixels needed
nfi_df = nfi_df[nfi_df.pred_count > 3]

nfi_df["region"] = pandas.Categorical(nfi_df.region)
nfi_df["naturalness"] = pandas.Categorical(nfi_df.naturalness, categories=[3, 1, 2])
nfi_df.naturalness.cat.rename_categories({
    1.0: "normal", 2.0: "natural", 3.0: "plantation"
}, inplace=True)

# drop rows with NA in natura2000 column
nfi_df.dropna(subset=["natura2000"], inplace=True)
nfi_df.natura2000.replace({0: "No", 1: "Yes"}, inplace=True)
nfi_df["natura2000"] = pandas.Categorical(nfi_df.natura2000)

# show the number of validation polygons per region
display(nfi_df.region.value_counts())
print("TOTAL PLOTS N: ", nfi_df.region.value_counts().sum())
```

```
2    5397
3    4623
1    3072
4     683
Name: region, dtype: int64
```

```
TOTAL PLOTS N:  13775
```

##### Boxplot -> forest naturalness classes¶

1=normal; 2=natural forest; 3=plantation forest

In [20]:

```
plots.plot_boxplots(
    nfi_df, "naturalness", height=4, aspect=0.75, fig_title="c) Plot-level validation - Forest naturalness",
    save_name="Figure4c.png", width=0.85
)
```

In [21]:

```
plots.describe_ci(nfi_df, "pred_mean", ["region", "naturalness"])
```

Out[21]:

|  |  | mean | count | std | ci95\_lo | ci95\_hi |
| --- | --- | --- | --- | --- | --- | --- |
| region | naturalness |  |  |  |  |  |
| 1 | plantation | 0.201100 | 19 | 0.125866 | 0.144504 | 0.257696 |
|  | normal | 0.354677 | 2897 | 0.257027 | 0.345317 | 0.364036 |
|  | natural | 0.849752 | 156 | 0.188187 | 0.820221 | 0.879283 |
| 2 | plantation | 0.239934 | 74 | 0.161381 | 0.203164 | 0.276704 |
|  | normal | 0.366578 | 5200 | 0.241487 | 0.360014 | 0.373141 |
|  | natural | 0.765518 | 123 | 0.241715 | 0.722800 | 0.808236 |
| 3 | plantation | 0.261172 | 103 | 0.195376 | 0.223440 | 0.298904 |
|  | normal | 0.365416 | 4494 | 0.213505 | 0.359174 | 0.371659 |
|  | natural | 0.626181 | 26 | 0.212064 | 0.544666 | 0.707695 |
| 4 | plantation | 0.320529 | 69 | 0.180607 | 0.277914 | 0.363144 |
|  | normal | 0.392742 | 612 | 0.223917 | 0.375002 | 0.410483 |
|  | natural | 0.374150 | 2 | 0.293237 | -0.032256 | 0.780556 |

In [22]:

```
plots.describe_ci(nfi_df, "pred_mean", ["region", "naturalness"])["count"].sum()
```

Out[22]:

```
13775
```

In [23]:

```
plots.tukey_subsets(nfi_df, "pred_mean", "naturalness", "region")
```

```
region: 1

    group1      group2      Diff     Lower     Upper    q-value   p-value
0   normal     natural  0.495075  0.446235  0.543916  33.614141  0.001000
1   normal  plantation  0.153577  0.016805  0.290349   3.723580  0.023137
2  natural  plantation  0.648652  0.504263  0.793041  14.897369  0.001000


region: 2

    group1      group2      Diff     Lower     Upper    q-value  p-value
0   normal     natural  0.398940  0.347486  0.450394  25.705613    0.001
1   normal  plantation  0.126644  0.060613  0.192675   6.358808    0.001
2  natural  plantation  0.525584  0.442607  0.608562  21.000113    0.001


region: 3

       group1      group2      Diff     Lower     Upper    q-value  p-value
0      normal  plantation  0.104245  0.054453  0.154036   6.941521    0.001
1      normal     natural  0.260764  0.162494  0.359035   8.798038    0.001
2  plantation     natural  0.365009  0.255350  0.474668  11.036146    0.001


region: 4

    group1      group2      Diff     Lower     Upper   q-value   p-value
0  natural      normal  0.018592 -0.347559  0.384743  0.168677  0.900000
1  natural  plantation  0.053621 -0.317193  0.424435  0.480354  0.900000
2   normal  plantation  0.072213  0.006563  0.137864  3.653927  0.026945
```

In [24]:

```
plots.plot_boxplots(
    nfi_df, "natura2000", height=4, aspect=0.75, fig_title="d) Plot-level validation - Natura 2000 habitat quality",
    save_name="Figure4d.png", width=0.5
)
```

In [25]:

```
plots.describe_ci(nfi_df, "pred_mean", ["region", "natura2000"])
```

Out[25]:

|  |  | mean | count | std | ci95\_lo | ci95\_hi |
| --- | --- | --- | --- | --- | --- | --- |
| region | natura2000 |  |  |  |  |  |
| 1 | No | 0.341749 | 2841 | 0.243279 | 0.332803 | 0.350695 |
|  | Yes | 0.835381 | 231 | 0.245292 | 0.803748 | 0.867014 |
| 2 | No | 0.357606 | 5216 | 0.233690 | 0.351264 | 0.363948 |
|  | Yes | 0.844459 | 181 | 0.188942 | 0.816932 | 0.871985 |
| 3 | No | 0.356986 | 4528 | 0.208634 | 0.350909 | 0.363063 |
|  | Yes | 0.725558 | 95 | 0.179260 | 0.689510 | 0.761606 |
| 4 | No | 0.375130 | 660 | 0.215492 | 0.358689 | 0.391570 |
|  | Yes | 0.679887 | 23 | 0.164647 | 0.612598 | 0.747176 |

In [26]:

```
plots.tukey_subsets(nfi_df, "pred_mean", "natura2000", "region")
```

```
region: 1

  group1 group2      Diff     Lower     Upper    q-value  p-value
0     No    Yes  0.493632  0.460976  0.526288  41.915508    0.001


region: 2

  group1 group2      Diff     Lower    Upper    q-value  p-value
0     No    Yes  0.486853  0.452416  0.52129  39.194787    0.001


region: 3

  group1 group2      Diff     Lower     Upper    q-value  p-value
0     No    Yes  0.368571  0.326282  0.410861  24.163736    0.001


region: 4

  group1 group2      Diff     Lower   Upper   q-value  p-value
0     No    Yes  0.304757  0.215614  0.3939  9.493002    0.001
```

### Hyperparameters tuning¶

In [3]:

```
# performance metrics after the hyperparameter tuning procedure using Bayesian optimization with Gaussian processes
# implemented in the scikit-optimize Python package

"""
# Search space for hyperparameters tuning has been define as follows:

search_space = list()
search_space.append(Categorical(["gini", "entropy", "log_loss"], name="criterion"))
search_space.append(Integer(3, 30, name="max_depth"))
search_space.append(Real(0.0001, 0.3, name="min_samples_split"))
search_space.append(Real(0.0001, 0.3, name="min_samples_leaf"))
search_space.append(Real(0.1, 1.0, name="max_features"))
"""

results = plots.load_models("./", r".*tuned\.pickle$")[0]

# get a table with metrics and associated best hyperparameters
metrics_sel = ["accuracy", "roc_auc", "pr_auc", "pearson_r", "briers_score", "MCC"]
columns = ["region"] + metrics_sel + results[0]["gp_results"].space.dimension_names
df_report = []

for key, result in results.items():
    values = [key + 1]
    metrics = result["calls_scv"][-1][metrics_sel].mean().tolist()
    hparams = result["gp_results"].x_iters[-1]
    values.extend(metrics + hparams)
    df_report.append(values)

df_report = pandas.DataFrame(df_report, columns=columns)
display(df_report)
```

```
Loading S1_full_models_tuned.pickle...
```

|  | region | accuracy | roc\_auc | pr\_auc | pearson\_r | briers\_score | MCC | criterion | max\_depth | min\_samples\_split | min\_samples\_leaf | max\_features |
| --- | --- | --- | --- | --- | --- | --- | --- | --- | --- | --- | --- | --- |
| 0 | 1 | 0.812498 | 0.898076 | 0.893445 | 0.692487 | 0.131931 | 0.624822 | log\_loss | 24 | 0.0001 | 0.0001 | 1.000000 |
| 1 | 2 | 0.816374 | 0.898897 | 0.875666 | 0.691302 | 0.131223 | 0.625582 | log\_loss | 30 | 0.0001 | 0.0001 | 0.562738 |
| 2 | 3 | 0.802748 | 0.887519 | 0.835070 | 0.659658 | 0.142107 | 0.596534 | log\_loss | 30 | 0.0001 | 0.0001 | 0.100000 |
| 3 | 4 | 0.815944 | 0.906412 | 0.877308 | 0.693585 | 0.139676 | 0.625998 | log\_loss | 30 | 0.0001 | 0.0001 | 0.100000 |

In [8]:

```
# performance metrics (default hyperparameters)
means, stds = plots.get_metrics(models_stats)
display(means)
display(stds)
```

|  | accuracy | TSS | roc\_auc | pr\_auc | pearson\_r | briers\_score | MCC |
| --- | --- | --- | --- | --- | --- | --- | --- |
| 0 | 0.814602 | 0.622555 | 0.895849 | 0.889916 | 0.685549 | 0.135498 | 0.627630 |
| 1 | 0.816158 | 0.619318 | 0.899882 | 0.878063 | 0.690780 | 0.133521 | 0.626025 |
| 2 | 0.806799 | 0.603023 | 0.894206 | 0.844005 | 0.666070 | 0.141412 | 0.609277 |
| 3 | 0.815561 | 0.499055 | 0.909073 | 0.880873 | 0.696555 | 0.139441 | 0.635410 |

|  | accuracy | TSS | roc\_auc | pr\_auc | pearson\_r | briers\_score | MCC |
| --- | --- | --- | --- | --- | --- | --- | --- |
| 0 | 0.027193 | 0.046479 | 0.022523 | 0.038950 | 0.040787 | 0.011069 | 0.056361 |
| 1 | 0.024461 | 0.029008 | 0.015541 | 0.027983 | 0.028191 | 0.008481 | 0.048373 |
| 2 | 0.023409 | 0.030500 | 0.020834 | 0.024094 | 0.034813 | 0.010400 | 0.045407 |
| 3 | 0.067950 | 0.134525 | 0.034206 | 0.052976 | 0.066607 | 0.021271 | 0.124319 |

In [ ]:
