## Supplementary material for "Mapping forests with different levels of naturalness using machine learning and landscape data mining": Jupyter Notebooks: M3b_full_nolatlon.html

validation


In [1]:

```
%load_ext autoreload
%autoreload 2

import warnings
warnings.filterwarnings("ignore")

from matplotlib import pyplot as plt
import matplotlib as mpl
mpl.rcParams["figure.dpi"] = 150

import json
import sys
import numpy as np
import pandas
import seaborn as sns
sns.set_style("white")

sys.path.insert(0, "/app/src")

from plots import plots
from config import settings
```

In [2]:

```
with open("./config.json", "r") as _file:
    mconfig = json.load(_file)
print(json.dumps(mconfig, indent=2))

model_name = mconfig["model_metadata"]["name"]
target_var = mconfig["model_specification"]["target_variable"]
region_var = mconfig["model_specification"]["stratification_layer"]
```

```
{
  "model_metadata": {
    "name": "S1_full_nolatlon",
    "description": "Full model using all available variables expressed at all target spatial scales. Seperate model for each ecoregion is fitted. Latitude and longitude excluded from the predictors list."
  },
  "model_data": {
    "samples_layer": "SAMPLES_1"
  },
  "model_specification": {
    "target_variable": "HCVF",
    "stratification_layer": "ecoregions",
    "stratification_submodels": [
      {
        "regions": [
          1
        ],
        "variables_list": []
      },
      {
        "regions": [
          2
        ],
        "variables_list": []
      },
      {
        "regions": [
          3
        ],
        "variables_list": []
      },
      {
        "regions": [
          4
        ],
        "variables_list": []
      }
    ],
    "lat_field": "LAT",
    "lon_field": "LON",
    "target_epsg": 3006,
    "variables_list": [],
    "variables_drop_regexp": "",
    "variables_keep_regexp": "",
    "variables_latlon": false,
    "balanced_rf": true
  },
  "model_crossvalidation": {
    "n_splits": 10,
    "spatial": true,
    "spatial_grid": "grid_20.geojson"
  },
  "model_hyperparameters": {
    "n_estimators": 500,
    "oob_score": true
  },
  "prediction": {
    "predict": true
  }
}
```

In [3]:

```
models = plots.load_models("./", regexp=fr"{model_name}_R[1-5]*.pickle")
models_stats = plots.get_cv_validation_stats(models)
```

```
Loading S1_full_nolatlon_R1.pickle...
Loading S1_full_nolatlon_R2.pickle...
Loading S1_full_nolatlon_R3.pickle...
Loading S1_full_nolatlon_R4.pickle...
```

In [4]:

```
print(models[0].data.shape)
models_labels=["North boreal", "South boreal", "Hemiboreal", "Nemoral"]

g = sns.FacetGrid(
    data=models[0].data, col=region_var, aspect=0.7, 
    sharey=False
)
g.map_dataframe(sns.countplot, x=target_var, palette=["black","red"])
axes = g.axes.flatten()
for i, ax in enumerate(axes):
    ax.set_title(models_labels[i], fontweight="bold", fontsize=16)
```

```
(14090, 130)
```

In [5]:

```
plots.plot_validation_curves_multimodel(
    models_stats, colors="bgrcmykw", figsize=(10, 5), plot_std=True, std_alpha=0.1,
    models_labels=["North boreal", "South boreal", "Hemiboreal", "Nemoral"],
    save_name="Figure2.png"
)
```

In [6]:

```
# performance metrics
means, stds = plots.get_metrics(models_stats)
display(means)
display(stds)
```

|  | accuracy | TSS | roc\_auc | pr\_auc | pearson\_r | briers\_score | MCC |
| --- | --- | --- | --- | --- | --- | --- | --- |
| 0 | 0.809611 | 0.608183 | 0.894559 | 0.889208 | 0.684084 | 0.135995 | 0.616974 |
| 1 | 0.815120 | 0.609761 | 0.898547 | 0.875440 | 0.688655 | 0.134326 | 0.624699 |
| 2 | 0.806709 | 0.595280 | 0.890767 | 0.838454 | 0.662392 | 0.142076 | 0.606959 |
| 3 | 0.807932 | 0.526326 | 0.901987 | 0.869634 | 0.685516 | 0.141097 | 0.612103 |

|  | accuracy | TSS | roc\_auc | pr\_auc | pearson\_r | briers\_score | MCC |
| --- | --- | --- | --- | --- | --- | --- | --- |
| 0 | 0.016914 | 0.045022 | 0.014472 | 0.021868 | 0.025100 | 0.007148 | 0.034500 |
| 1 | 0.014486 | 0.037441 | 0.013196 | 0.012126 | 0.021392 | 0.005554 | 0.030702 |
| 2 | 0.027766 | 0.044678 | 0.017914 | 0.031528 | 0.034952 | 0.012771 | 0.046903 |
| 3 | 0.042135 | 0.182705 | 0.028922 | 0.043243 | 0.049742 | 0.014293 | 0.084718 |

In [7]:

```
# number of variables per model
for i, model in enumerate(models):
    print(f"Model_{i+1}: ", len(model.cov_names))
```

```
Model_1:  126
Model_2:  126
Model_3:  126
Model_4:  126
```

In [8]:

```
# plot features Gini's importances
for i, model in enumerate(models):
    print(models_labels[i])
    _plot = plots.plot_gini_importances(model, figsize=(14, 10), n=50)
```

```
North boreal
```

```
South boreal
```

```
Hemiboreal
```

```
Nemoral
```

##### Partial Dependence Plot¶

In [9]:

```
plots.plot_partial_dependences(models, n_features=6)
```

##### Accumulated Local Effects Plot¶

In [10]:

```
plots.plot_ales(models, n_features=6, xlims=(-2, 2))
```

In [11]:

```
# average max tree depth per model
import numpy as np
for i, model in enumerate(models):
    max_depths = []
    for tree_idx, est in enumerate(model.model.estimators_):
        max_depths.append(est.tree_.max_depth)
    print(f"{models_labels[i]}: ", np.array(max_depths).mean())
```

```
North boreal:  22.066
South boreal:  21.082
Hemiboreal:  20.01
Nemoral:  10.672
```

### Validation¶

#### Sveaskog validation¶

tree stand (polygon) level validation; minimum 10 pixels with predicted value

In [12]:

```
# load a dataframe with predictions and Sveaskog atributes
svea_df = pandas.read_csv("svea_predictions.csv")
svea_df = plots.svea_preprocess(svea_df)

# filter out data not reliable for validation
# min number of 1 ha pixels to calculate the mean probability
svea_df = svea_df[svea_df.pred_count > 10]

# show the number of validation polygons per region
display(svea_df.region.value_counts())
print("TOTAL POLYGONS N: ", svea_df.region.value_counts().sum())
```

```
1    41784
2     7989
3     7519
4      256
Name: region, dtype: int64
```

```
TOTAL POLYGONS N:  57548
```

##### Boxplot -> forest management classes¶

"NF" (conservation, no management)
"NF\_NM" (conservation, no yet specified)
"NM" (conservation, management)
"PE" (production forest, increased conservation considerations)
"PG" (production forest, general conservation conciderations)

Management categories for stands. A goal for long-term management is set for each stand, in one of four categories: (i) NF = Nature conservation, free development; (ii) NM = Nature conservation-oriented management, often implying restoration measures; (iii) PG = Production with general conservation concern (green-tree and deadwood retention for biodiversity at harvest, represented by black dots). The minimum level of retention is specified, and always >15 % of the harvested area; (iv) PE = Production with enhanced conservation concern.

In [13]:

```
plots.plot_boxplots(
    svea_df, "management", height=4, aspect=0.75, fig_title="a) Stand-level validation - Forest management objectives",
    save_name="Figure4a.png", width=0.85
)
```

In [14]:

```
plots.describe_ci(svea_df, "pred_mean", ["region", "management"])
```

Out[14]:

|  |  | mean | count | std | ci95\_lo | ci95\_hi |
| --- | --- | --- | --- | --- | --- | --- |
| region | management |  |  |  |  |  |
| 1 | NF | 0.632428 | 4148 | 0.222637 | 0.625653 | 0.639204 |
|  | NF\_NM | 0.720145 | 2401 | 0.180041 | 0.712943 | 0.727347 |
|  | NM | 0.567502 | 903 | 0.205151 | 0.554121 | 0.580883 |
|  | PE | 0.400831 | 2516 | 0.197640 | 0.393108 | 0.408554 |
|  | PG | 0.271262 | 31816 | 0.170666 | 0.269387 | 0.273137 |
| 2 | NF | 0.681095 | 1245 | 0.195617 | 0.670229 | 0.691962 |
|  | NF\_NM | 0.700983 | 82 | 0.181331 | 0.661735 | 0.740231 |
|  | NM | 0.536016 | 253 | 0.211386 | 0.509968 | 0.562064 |
|  | PE | 0.397541 | 868 | 0.211826 | 0.383449 | 0.411633 |
|  | PG | 0.275590 | 5541 | 0.184478 | 0.270733 | 0.280448 |
| 3 | NF | 0.597362 | 297 | 0.183872 | 0.576450 | 0.618274 |
|  | NF\_NM | 0.670740 | 172 | 0.158920 | 0.646990 | 0.694491 |
|  | NM | 0.582141 | 303 | 0.190301 | 0.560714 | 0.603569 |
|  | PE | 0.414052 | 608 | 0.181787 | 0.399602 | 0.428502 |
|  | PG | 0.298777 | 6139 | 0.161033 | 0.294748 | 0.302805 |
| 4 | NF | 0.747433 | 5 | 0.054077 | 0.700032 | 0.794833 |
|  | NF\_NM | NaN | 0 | NaN | NaN | NaN |
|  | NM | 0.624948 | 19 | 0.181306 | 0.543423 | 0.706473 |
|  | PE | 0.577465 | 13 | 0.164399 | 0.488097 | 0.666834 |
|  | PG | 0.326947 | 219 | 0.157168 | 0.306131 | 0.347763 |

In [15]:

```
plots.tukey_subsets(svea_df, "pred_mean", "management", "region")
```

```
region: 1

  group1 group2      Diff     Lower     Upper     q-value  p-value
0     NF  NF_NM  0.087415  0.074723  0.100108   26.569103    0.001
1     NF     PG  0.362823  0.354646  0.371001  171.172465    0.001
2     NF     PE  0.212347  0.198656  0.226038   59.833925    0.001
3     NF     NM  0.066839  0.048632  0.085046   14.162284    0.001
4  NF_NM     PG  0.450239  0.439748  0.460729  165.575521    0.001
5  NF_NM     PE  0.299762  0.284576  0.314949   76.148641    0.001
6  NF_NM     NM  0.154254  0.134898  0.173610   30.743413    0.001
7     PG     PE  0.150476  0.138798  0.162155   49.706745    0.001
8     PG     NM  0.295985  0.279239  0.312731   68.185871    0.001
9     PE     NM  0.145509  0.125483  0.165534   28.031621    0.001


region: 2

  group1 group2      Diff     Lower     Upper    q-value   p-value
0     PG     PE  0.122578  0.100932  0.144225  21.850133  0.001000
1     PG     NF  0.410579  0.394035  0.427123  95.761153  0.001000
2     PG     NM  0.266009  0.232208  0.299809  30.366752  0.001000
3     PG  NF_NM  0.430858  0.372526  0.489189  28.500885  0.001000
4     PE     NF  0.288001  0.262750  0.313252  44.009188  0.001000
5     PE     NM  0.143430  0.104618  0.182243  14.259330  0.001000
6     PE  NF_NM  0.308280  0.246908  0.369651  19.382267  0.001000
7     NF     NM  0.144570  0.108355  0.180785  15.403460  0.001000
8     NF  NF_NM  0.020279 -0.039484  0.080042   1.309296  0.879913
9     NM  NF_NM  0.164849  0.098213  0.231485   9.545605  0.001000


region: 3

  group1 group2      Diff     Lower     Upper    q-value   p-value
0     PE     NF  0.176872  0.144420  0.209324  21.030271  0.001000
1     PE     PG  0.121903  0.101867  0.141940  23.475891  0.001000
2     PE  NF_NM  0.253348  0.213784  0.292912  24.708482  0.001000
3     PE     NM  0.161529  0.128933  0.194125  19.121180  0.001000
4     NF     PG  0.298775  0.271951  0.325598  42.978953  0.001000
5     NF  NF_NM  0.076477  0.033079  0.119874   6.799710  0.001000
6     NF     NM  0.015343 -0.021813  0.052499   1.593370  0.765927
7     PG  NF_NM  0.375251  0.340156  0.410347  41.257390  0.001000
8     PG     NM  0.283432  0.256434  0.310429  40.509212  0.001000
9  NF_NM     NM  0.091820  0.048314  0.135325   8.143689  0.001000


region: 4

  group1 group2      Diff     Lower     Upper    q-value   p-value
0     PG     NM  0.306285  0.208022  0.404549  11.407483  0.001000
1     PG     NF  0.421839  0.215073  0.628604   7.466646  0.001000
2     PG     PE  0.245474  0.118689  0.372259   7.085908  0.001000
3     NM     NF  0.115553 -0.109719  0.340825   1.877293  0.541039
4     NM     PE  0.060811 -0.094334  0.215956   1.434505  0.716368
5     NF     PE  0.176364 -0.062730  0.415459   2.699594  0.227240
```

##### Boxplot -> forest naturalness classes¶

In [16]:

```
plots.plot_boxplots(
    svea_df, "naturalness", height=4, aspect=0.75, fig_title="b) Stand-level validation - Forest naturalness",
    save_name="Figure4b.png", width=0.5
)
```

In [17]:

```
plots.describe_ci(svea_df, "pred_mean", ["region", "naturalness"])
```

Out[17]:

|  |  | mean | count | std | ci95\_lo | ci95\_hi |
| --- | --- | --- | --- | --- | --- | --- |
| region | naturalness |  |  |  |  |  |
| 1 | No | 0.279801 | 34130 | 0.175276 | 0.277942 | 0.281661 |
|  | Yes | 0.647268 | 7654 | 0.216726 | 0.642412 | 0.652123 |
| 2 | No | 0.290515 | 6306 | 0.191801 | 0.285781 | 0.295249 |
|  | Yes | 0.642413 | 1683 | 0.216421 | 0.632073 | 0.652753 |
| 3 | No | 0.306955 | 6602 | 0.165929 | 0.302953 | 0.310958 |
|  | Yes | 0.576429 | 917 | 0.193508 | 0.563904 | 0.588954 |
| 4 | No | 0.342715 | 233 | 0.169193 | 0.320990 | 0.364440 |
|  | Yes | 0.646392 | 23 | 0.172366 | 0.575949 | 0.716836 |

In [18]:

```
plots.tukey_subsets(svea_df, "pred_mean", "naturalness", "region")
```

```
region: 1

  group1 group2      Diff     Lower     Upper     q-value  p-value
0    Yes     No  0.369747  0.365123  0.374372  221.639364    0.001


region: 2

  group1 group2      Diff     Lower    Upper    q-value  p-value
0     No    Yes  0.360545  0.349749  0.37134  92.586351    0.001


region: 3

  group1 group2      Diff     Lower    Upper    q-value  p-value
0     No    Yes  0.270108  0.258256  0.28196  63.181537    0.001


region: 4

  group1 group2      Diff    Lower     Upper    q-value  p-value
0     No    Yes  0.307271  0.23296  0.381581  11.520886    0.001
```

#### NFI validation¶

plot level validation (1 central pixel + 4 neighbours)

In [19]:

```
# load a dataframe with predictions and NFI atributes
nfi_df = pandas.read_csv("nfi_predictions.csv")

# at least 4 pixels needed
nfi_df = nfi_df[nfi_df.pred_count > 3]

nfi_df["region"] = pandas.Categorical(nfi_df.region)
nfi_df["naturalness"] = pandas.Categorical(nfi_df.naturalness, categories=[3, 1, 2])
nfi_df.naturalness.cat.rename_categories({
    1.0: "normal", 2.0: "natural", 3.0: "plantation"
}, inplace=True)

# drop rows with NA in natura2000 column
nfi_df.dropna(subset=["natura2000"], inplace=True)
nfi_df.natura2000.replace({0: "No", 1: "Yes"}, inplace=True)
nfi_df["natura2000"] = pandas.Categorical(nfi_df.natura2000)

# show the number of validation polygons per region
display(nfi_df.region.value_counts())
print("TOTAL PLOTS N: ", nfi_df.region.value_counts().sum())
```

```
2    5397
3    4623
1    3072
4     683
Name: region, dtype: int64
```

```
TOTAL PLOTS N:  13775
```

##### Boxplot -> forest naturalness classes¶

1=normal; 2=natural forest; 3=plantation forest

In [20]:

```
plots.plot_boxplots(
    nfi_df, "naturalness", height=4, aspect=0.75, fig_title="c) Plot-level validation - Forest naturalness",
    save_name="Figure4c.png", width=0.85
)
```

In [21]:

```
plots.describe_ci(nfi_df, "pred_mean", ["region", "naturalness"])
```

Out[21]:

|  |  | mean | count | std | ci95\_lo | ci95\_hi |
| --- | --- | --- | --- | --- | --- | --- |
| region | naturalness |  |  |  |  |  |
| 1 | plantation | 0.191758 | 19 | 0.120624 | 0.137519 | 0.245997 |
|  | normal | 0.354369 | 2897 | 0.258726 | 0.344948 | 0.363791 |
|  | natural | 0.848196 | 156 | 0.186810 | 0.818880 | 0.877511 |
| 2 | plantation | 0.236561 | 74 | 0.158312 | 0.200490 | 0.272632 |
|  | normal | 0.365877 | 5200 | 0.240849 | 0.359330 | 0.372423 |
|  | natural | 0.764665 | 123 | 0.238379 | 0.722537 | 0.806793 |
| 3 | plantation | 0.258925 | 103 | 0.198589 | 0.220573 | 0.297278 |
|  | normal | 0.365355 | 4494 | 0.214103 | 0.359096 | 0.371615 |
|  | natural | 0.624531 | 26 | 0.219924 | 0.539995 | 0.709067 |
| 4 | plantation | 0.322858 | 69 | 0.178418 | 0.280759 | 0.364957 |
|  | normal | 0.392249 | 612 | 0.224145 | 0.374490 | 0.410007 |
|  | natural | 0.389600 | 2 | 0.328663 | -0.065904 | 0.845104 |

In [22]:

```
plots.describe_ci(nfi_df, "pred_mean", ["region", "naturalness"])["count"].sum()
```

Out[22]:

```
13775
```

In [23]:

```
plots.tukey_subsets(nfi_df, "pred_mean", "naturalness", "region")
```

```
region: 1

    group1      group2      Diff     Lower     Upper    q-value   p-value
0   normal     natural  0.493826  0.444685  0.542967  33.324187  0.001000
1   normal  plantation  0.162612  0.024998  0.300225   3.918510  0.015529
2  natural  plantation  0.656438  0.511160  0.801715  14.983936  0.001000


region: 2

    group1      group2      Diff     Lower     Upper    q-value  p-value
0   normal     natural  0.398788  0.347488  0.450088  25.772981    0.001
1   normal  plantation  0.129316  0.063482  0.195149   6.512459    0.001
2  natural  plantation  0.528104  0.445375  0.610833  21.164160    0.001


region: 3

       group1      group2      Diff     Lower     Upper    q-value  p-value
0      normal  plantation  0.106430  0.056477  0.156383   7.064163    0.001
1      normal     natural  0.259175  0.160587  0.357764   8.716180    0.001
2  plantation     natural  0.365606  0.255591  0.475620  11.018477    0.001


region: 4

    group1      group2      Diff     Lower     Upper   q-value   p-value
0  natural      normal  0.002649 -0.363675  0.368973  0.024020  0.900000
1  natural  plantation  0.066742 -0.304247  0.437732  0.597613  0.900000
2   normal  plantation  0.069391  0.003709  0.135073  3.509455  0.035507
```

In [24]:

```
plots.plot_boxplots(
    nfi_df, "natura2000", height=4, aspect=0.75, fig_title="d) Plot-level validation - Natura 2000 habitat quality",
    save_name="Figure4d.png", width=0.5
)
```

In [25]:

```
plots.describe_ci(nfi_df, "pred_mean", ["region", "natura2000"])
```

Out[25]:

|  |  | mean | count | std | ci95\_lo | ci95\_hi |
| --- | --- | --- | --- | --- | --- | --- |
| region | natura2000 |  |  |  |  |  |
| 1 | No | 0.341310 | 2841 | 0.244891 | 0.332305 | 0.350316 |
|  | Yes | 0.835097 | 231 | 0.245701 | 0.803412 | 0.866782 |
| 2 | No | 0.356871 | 5216 | 0.233020 | 0.350547 | 0.363194 |
|  | Yes | 0.843539 | 181 | 0.186505 | 0.816368 | 0.870710 |
| 3 | No | 0.356872 | 4528 | 0.209334 | 0.350775 | 0.362970 |
|  | Yes | 0.725220 | 95 | 0.181594 | 0.688703 | 0.761737 |
| 4 | No | 0.374657 | 660 | 0.215071 | 0.358248 | 0.391065 |
|  | Yes | 0.688661 | 23 | 0.164415 | 0.621466 | 0.755855 |

In [26]:

```
plots.tukey_subsets(nfi_df, "pred_mean", "natura2000", "region")
```

```
region: 1

  group1 group2      Diff     Lower     Upper    q-value  p-value
0     No    Yes  0.493787  0.460927  0.526647  41.668216    0.001


region: 2

  group1 group2      Diff     Lower     Upper    q-value  p-value
0     No    Yes  0.486668  0.452337  0.520999  39.301272    0.001


region: 3

  group1 group2      Diff    Lower     Upper    q-value  p-value
0     No    Yes  0.368348  0.32591  0.410785  24.064829    0.001


region: 4

  group1 group2      Diff     Lower     Upper   q-value  p-value
0     No    Yes  0.314004  0.225034  0.402974  9.800065    0.001
```

In [ ]:
