## Supplementary material for "Mapping forests with different levels of naturalness using machine learning and landscape data mining": Jupyter Notebooks: M4a_onescale.html

validation


In [1]:

```
%load_ext autoreload
%autoreload 2

import warnings
warnings.filterwarnings("ignore")

from matplotlib import pyplot as plt
import matplotlib as mpl
mpl.rcParams["figure.dpi"] = 150

import json
import sys
import numpy as np
import pandas
import seaborn as sns
sns.set_style("white")

sys.path.insert(0, "/app/src")

from plots import plots
from config import settings
```

In [2]:

```
with open("./config.json", "r") as _file:
    mconfig = json.load(_file)
print(json.dumps(mconfig, indent=2))

model_name = mconfig["model_metadata"]["name"]
target_var = mconfig["model_specification"]["target_variable"]
region_var = mconfig["model_specification"]["stratification_layer"]
```

```
{
  "model_metadata": {
    "name": "S1_onescale",
    "description": "One-scale model i.e. only 1ha scale predictors are included. Seperate model for each region."
  },
  "model_data": {
    "samples_layer": "SAMPLES_1"
  },
  "model_specification": {
    "target_variable": "HCVF",
    "stratification_layer": "ecoregions",
    "stratification_submodels": [
      {
        "regions": [
          1
        ],
        "variables_list": []
      },
      {
        "regions": [
          2
        ],
        "variables_list": []
      },
      {
        "regions": [
          3
        ],
        "variables_list": []
      },
      {
        "regions": [
          4
        ],
        "variables_list": []
      }
    ],
    "lat_field": "LAT",
    "lon_field": "LON",
    "target_epsg": 3006,
    "variables_list": [],
    "variables_drop_regexp": "([A-Z]?[0-9]{3})",
    "variables_keep_regexp": "",
    "variables_latlon": true,
    "balanced_rf": true
  },
  "model_crossvalidation": {
    "n_splits": 10,
    "spatial": true,
    "spatial_grid": "grid_20.geojson"
  },
  "model_hyperparameters": {
    "n_estimators": 500,
    "oob_score": true
  },
  "prediction": {
    "predict": true
  }
}
```

In [3]:

```
models = plots.load_models("./", regexp=fr"{model_name}_R[1-5]*.pickle")
models_stats = plots.get_cv_validation_stats(models)
```

```
Loading S1_onescale_R1.pickle...
Loading S1_onescale_R2.pickle...
Loading S1_onescale_R3.pickle...
Loading S1_onescale_R4.pickle...
```

In [4]:

```
print(models[0].data.shape)
models_labels=["North boreal", "South boreal", "Hemiboreal", "Nemoral"]

g = sns.FacetGrid(
    data=models[0].data, col=region_var, aspect=0.7, 
    sharey=False
)
g.map_dataframe(sns.countplot, x=target_var, palette=["black","red"])
axes = g.axes.flatten()
for i, ax in enumerate(axes):
    ax.set_title(models_labels[i], fontweight="bold", fontsize=16)
```

```
(14090, 130)
```

In [5]:

```
plots.plot_validation_curves_multimodel(
    models_stats, colors="bgrcmykw", figsize=(10, 5), plot_std=True, std_alpha=0.1,
    models_labels=["North boreal", "South boreal", "Hemiboreal", "Nemoral"],
    save_name="Figure2.png"
)
```

In [6]:

```
# performance metrics
means, stds = plots.get_metrics(models_stats)
display(means)
display(stds)
```

|  | accuracy | TSS | roc\_auc | pr\_auc | pearson\_r | briers\_score | MCC |
| --- | --- | --- | --- | --- | --- | --- | --- |
| 0 | 0.796475 | 0.582539 | 0.880505 | 0.873429 | 0.659526 | 0.142797 | 0.590688 |
| 1 | 0.799609 | 0.586713 | 0.883608 | 0.850191 | 0.661544 | 0.141215 | 0.590725 |
| 2 | 0.780493 | 0.539205 | 0.859213 | 0.792796 | 0.605757 | 0.158208 | 0.550600 |
| 3 | 0.789803 | 0.507427 | 0.859122 | 0.811163 | 0.611651 | 0.160327 | 0.563882 |

|  | accuracy | TSS | roc\_auc | pr\_auc | pearson\_r | briers\_score | MCC |
| --- | --- | --- | --- | --- | --- | --- | --- |
| 0 | 0.026135 | 0.066770 | 0.028947 | 0.032046 | 0.050228 | 0.014554 | 0.052531 |
| 1 | 0.015704 | 0.043168 | 0.014701 | 0.036924 | 0.028166 | 0.007134 | 0.034968 |
| 2 | 0.025284 | 0.039980 | 0.020691 | 0.025389 | 0.032796 | 0.009616 | 0.046994 |
| 3 | 0.079842 | 0.124877 | 0.067274 | 0.110909 | 0.117715 | 0.025536 | 0.159735 |

In [7]:

```
# number of variables per model
for i, model in enumerate(models):
    print(f"Model_{i+1}: ", len(model.cov_names))
```

```
Model_1:  28
Model_2:  28
Model_3:  28
Model_4:  28
```

In [8]:

```
# plot features Gini's importances
for i, model in enumerate(models):
    print(models_labels[i])
    _plot = plots.plot_gini_importances(model, figsize=(14, 10), n=50)
```

```
North boreal
```

```
South boreal
```

```
Hemiboreal
```

```
Nemoral
```

##### Partial Dependence Plot¶

In [9]:

```
plots.plot_partial_dependences(models, n_features=6)
```

##### Accumulated Local Effects Plot¶

In [10]:

```
plots.plot_ales(models, n_features=6, xlims=(-2, 2))
```

In [11]:

```
# average max tree depth per model
import numpy as np
for i, model in enumerate(models):
    max_depths = []
    for tree_idx, est in enumerate(model.model.estimators_):
        max_depths.append(est.tree_.max_depth)
    print(f"{models_labels[i]}: ", np.array(max_depths).mean())
```

```
North boreal:  22.142
South boreal:  21.43
Hemiboreal:  21.38
Nemoral:  12.696
```

### Validation¶

#### Sveaskog validation¶

tree stand (polygon) level validation; minimum 10 pixels with predicted value

In [12]:

```
# load a dataframe with predictions and Sveaskog atributes
svea_df = pandas.read_csv("svea_predictions.csv")
svea_df = plots.svea_preprocess(svea_df)

# filter out data not reliable for validation
# min number of 1 ha pixels to calculate the mean probability
svea_df = svea_df[svea_df.pred_count > 10]

# show the number of validation polygons per region
display(svea_df.region.value_counts())
print("TOTAL POLYGONS N: ", svea_df.region.value_counts().sum())
```

```
1    41784
2     7989
3     7519
4      256
Name: region, dtype: int64
```

```
TOTAL POLYGONS N:  57548
```

##### Boxplot -> forest management classes¶

"NF" (conservation, no management)
"NF\_NM" (conservation, no yet specified)
"NM" (conservation, management)
"PE" (production forest, increased conservation considerations)
"PG" (production forest, general conservation conciderations)

Management categories for stands. A goal for long-term management is set for each stand, in one of four categories: (i) NF = Nature conservation, free development; (ii) NM = Nature conservation-oriented management, often implying restoration measures; (iii) PG = Production with general conservation concern (green-tree and deadwood retention for biodiversity at harvest, represented by black dots). The minimum level of retention is specified, and always >15 % of the harvested area; (iv) PE = Production with enhanced conservation concern.

In [13]:

```
plots.plot_boxplots(
    svea_df, "management", height=4, aspect=0.75, fig_title="a) Stand-level validation - Forest management objectives",
    save_name="Figure4a.png", width=0.85
)
```

In [14]:

```
plots.describe_ci(svea_df, "pred_mean", ["region", "management"])
```

Out[14]:

|  |  | mean | count | std | ci95\_lo | ci95\_hi |
| --- | --- | --- | --- | --- | --- | --- |
| region | management |  |  |  |  |  |
| 1 | NF | 0.657104 | 4148 | 0.199209 | 0.651042 | 0.663166 |
|  | NF\_NM | 0.714305 | 2401 | 0.169625 | 0.707520 | 0.721090 |
|  | NM | 0.557253 | 903 | 0.203309 | 0.543992 | 0.570513 |
|  | PE | 0.403988 | 2516 | 0.194195 | 0.396400 | 0.411576 |
|  | PG | 0.269481 | 31816 | 0.160293 | 0.267720 | 0.271243 |
| 2 | NF | 0.666836 | 1245 | 0.174962 | 0.657117 | 0.676555 |
|  | NF\_NM | 0.678004 | 82 | 0.188118 | 0.637287 | 0.718722 |
|  | NM | 0.506855 | 253 | 0.202239 | 0.481934 | 0.531776 |
|  | PE | 0.376521 | 868 | 0.199424 | 0.363254 | 0.389788 |
|  | PG | 0.259233 | 5541 | 0.169185 | 0.254778 | 0.263687 |
| 3 | NF | 0.617998 | 297 | 0.159653 | 0.599840 | 0.636155 |
|  | NF\_NM | 0.640726 | 172 | 0.161886 | 0.616533 | 0.664920 |
|  | NM | 0.541574 | 303 | 0.188397 | 0.520360 | 0.562787 |
|  | PE | 0.374330 | 608 | 0.173271 | 0.360557 | 0.388103 |
|  | PG | 0.294015 | 6139 | 0.143595 | 0.290423 | 0.297607 |
| 4 | NF | 0.688208 | 5 | 0.079932 | 0.618144 | 0.758271 |
|  | NF\_NM | NaN | 0 | NaN | NaN | NaN |
|  | NM | 0.592578 | 19 | 0.188379 | 0.507872 | 0.677283 |
|  | PE | 0.483446 | 13 | 0.151323 | 0.401186 | 0.565707 |
|  | PG | 0.285976 | 219 | 0.135902 | 0.267977 | 0.303976 |

In [15]:

```
plots.tukey_subsets(svea_df, "pred_mean", "management", "region")
```

```
region: 1

  group1 group2      Diff     Lower     Upper     q-value  p-value
0     NF  NF_NM  0.056863  0.044969  0.068757   18.443890    0.001
1     NF     PG  0.389587  0.381924  0.397249  196.144250    0.001
2     NF     PE  0.238275  0.225446  0.251105   71.649493    0.001
3     NF     NM  0.101603  0.084542  0.118663   22.974318    0.001
4  NF_NM     PG  0.446450  0.436620  0.456280  175.209953    0.001
5  NF_NM     PE  0.295138  0.280908  0.309369   80.009883    0.001
6  NF_NM     NM  0.158466  0.140327  0.176604   33.704163    0.001
7     PG     PE  0.151311  0.140368  0.162255   53.339820    0.001
8     PG     NM  0.287984  0.272292  0.303676   70.798912    0.001
9     PE     NM  0.136673  0.117908  0.155438   28.097986    0.001


region: 2

  group1 group2      Diff     Lower     Upper     q-value  p-value
0     PG     PE  0.117648  0.097771  0.137525   22.838473    0.001
1     PG     NF  0.412274  0.397082  0.427465  104.717569    0.001
2     PG     NM  0.253486  0.222449  0.284523   31.513615    0.001
3     PG  NF_NM  0.423783  0.370221  0.477346   30.528795    0.001
4     PE     NF  0.294626  0.271439  0.317812   49.029948    0.001
5     PE     NM  0.135838  0.100199  0.171477   14.706885    0.001
6     PE  NF_NM  0.306135  0.249781  0.362489   20.961123    0.001
7     NF     NM  0.158787  0.125533  0.192042   18.424538    0.001
8     NF  NF_NM  0.011510 -0.043367  0.066387    0.809286    0.900
9     NM  NF_NM  0.170297  0.109109  0.231485   10.739044    0.001


region: 3

  group1 group2      Diff     Lower     Upper    q-value   p-value
0     PE     NF  0.240635  0.211274  0.269997  31.623720  0.001000
1     PE     PG  0.083339  0.065211  0.101467  17.738716  0.001000
2     PE  NF_NM  0.266518  0.230722  0.302314  28.729092  0.001000
3     PE     NM  0.165257  0.135765  0.194748  21.621792  0.001000
4     NF     PG  0.323974  0.299705  0.348243  51.509750  0.001000
5     NF  NF_NM  0.025883 -0.013382  0.065147   2.543543  0.375383
6     NF     NM  0.075378  0.041762  0.108995   8.652043  0.001000
7     PG  NF_NM  0.349857  0.318104  0.381610  42.514506  0.001000
8     PG     NM  0.248596  0.224169  0.273022  39.270472  0.001000
9  NF_NM     NM  0.101261  0.061899  0.140623   9.926498  0.001000


region: 4

  group1 group2      Diff     Lower     Upper    q-value   p-value
0     PG     NM  0.314647  0.227136  0.402158  13.158845  0.001000
1     PG     NF  0.384934  0.200795  0.569074   7.650624  0.001000
2     PG     PE  0.181344  0.068433  0.294255   5.877919  0.001000
3     NM     NF  0.070288 -0.130333  0.270909   1.282211  0.776671
4     NM     PE  0.133303 -0.004865  0.271471   3.530935  0.063131
5     NF     PE  0.203590 -0.009341  0.416521   3.499258  0.066763
```

##### Boxplot -> forest naturalness classes¶

In [16]:

```
plots.plot_boxplots(
    svea_df, "naturalness", height=4, aspect=0.75, fig_title="b) Stand-level validation - Forest naturalness",
    save_name="Figure4b.png", width=0.5
)
```

In [17]:

```
plots.describe_ci(svea_df, "pred_mean", ["region", "naturalness"])
```

Out[17]:

|  |  | mean | count | std | ci95\_lo | ci95\_hi |
| --- | --- | --- | --- | --- | --- | --- |
| region | naturalness |  |  |  |  |  |
| 1 | No | 0.278591 | 34130 | 0.166130 | 0.276828 | 0.280353 |
|  | Yes | 0.656633 | 7654 | 0.201207 | 0.652126 | 0.661141 |
| 2 | No | 0.273923 | 6306 | 0.177203 | 0.269549 | 0.278296 |
|  | Yes | 0.623835 | 1683 | 0.203881 | 0.614094 | 0.633575 |
| 3 | No | 0.300895 | 6602 | 0.148293 | 0.297318 | 0.304472 |
|  | Yes | 0.549498 | 917 | 0.199745 | 0.536570 | 0.562427 |
| 4 | No | 0.298599 | 233 | 0.145489 | 0.279918 | 0.317281 |
|  | Yes | 0.610435 | 23 | 0.178142 | 0.537631 | 0.683240 |

In [18]:

```
plots.tukey_subsets(svea_df, "pred_mean", "naturalness", "region")
```

```
region: 1

  group1 group2      Diff     Lower     Upper     q-value  p-value
0    Yes     No  0.380988  0.376634  0.385343  242.524808    0.001


region: 2

  group1 group2      Diff     Lower     Upper    q-value  p-value
0     No    Yes  0.358029  0.348015  0.368042  99.124733    0.001


region: 3

  group1 group2      Diff     Lower    Upper    q-value  p-value
0     No    Yes  0.249587  0.238724  0.26045  63.695285    0.001


region: 4

  group1 group2      Diff     Lower     Upper    q-value  p-value
0     No    Yes  0.313255  0.248124  0.378386  13.400629    0.001
```

#### NFI validation¶

plot level validation (1 central pixel + 4 neighbours)

In [19]:

```
# load a dataframe with predictions and NFI atributes
nfi_df = pandas.read_csv("nfi_predictions.csv")

# at least 4 pixels needed
nfi_df = nfi_df[nfi_df.pred_count > 3]

nfi_df["region"] = pandas.Categorical(nfi_df.region)
nfi_df["naturalness"] = pandas.Categorical(nfi_df.naturalness, categories=[3, 1, 2])
nfi_df.naturalness.cat.rename_categories({
    1.0: "normal", 2.0: "natural", 3.0: "plantation"
}, inplace=True)

# drop rows with NA in natura2000 column
nfi_df.dropna(subset=["natura2000"], inplace=True)
nfi_df.natura2000.replace({0: "No", 1: "Yes"}, inplace=True)
nfi_df["natura2000"] = pandas.Categorical(nfi_df.natura2000)

# show the number of validation polygons per region
display(nfi_df.region.value_counts())
print("TOTAL PLOTS N: ", nfi_df.region.value_counts().sum())
```

```
2    5397
3    4623
1    3072
4     683
Name: region, dtype: int64
```

```
TOTAL PLOTS N:  13775
```

##### Boxplot -> forest naturalness classes¶

1=normal; 2=natural forest; 3=plantation forest

In [20]:

```
plots.plot_boxplots(
    nfi_df, "naturalness", height=4, aspect=0.75, fig_title="c) Plot-level validation - Forest naturalness",
    save_name="Figure4c.png", width=0.85
)
```

In [21]:

```
plots.describe_ci(nfi_df, "pred_mean", ["region", "naturalness"])
```

Out[21]:

|  |  | mean | count | std | ci95\_lo | ci95\_hi |
| --- | --- | --- | --- | --- | --- | --- |
| region | naturalness |  |  |  |  |  |
| 1 | plantation | 0.175805 | 19 | 0.102767 | 0.129596 | 0.222015 |
|  | normal | 0.349462 | 2897 | 0.244655 | 0.340553 | 0.358371 |
|  | natural | 0.823523 | 156 | 0.171393 | 0.796627 | 0.850419 |
| 2 | plantation | 0.239409 | 74 | 0.138909 | 0.207760 | 0.271059 |
|  | normal | 0.362109 | 5200 | 0.223184 | 0.356043 | 0.368175 |
|  | natural | 0.739977 | 123 | 0.217532 | 0.701533 | 0.778421 |
| 3 | plantation | 0.289700 | 103 | 0.166305 | 0.257582 | 0.321818 |
|  | normal | 0.376630 | 4494 | 0.186557 | 0.371176 | 0.382085 |
|  | natural | 0.603615 | 26 | 0.175282 | 0.536239 | 0.670992 |
| 4 | plantation | 0.303751 | 69 | 0.131673 | 0.272682 | 0.334820 |
|  | normal | 0.381305 | 612 | 0.173916 | 0.367526 | 0.395084 |
|  | natural | 0.473450 | 2 | 0.241901 | 0.138192 | 0.808708 |

In [22]:

```
plots.describe_ci(nfi_df, "pred_mean", ["region", "naturalness"])["count"].sum()
```

Out[22]:

```
13775
```

In [23]:

```
plots.tukey_subsets(nfi_df, "pred_mean", "naturalness", "region")
```

```
region: 1

    group1      group2      Diff     Lower     Upper    q-value   p-value
0   normal     natural  0.474061  0.427635  0.520487  33.861383  0.001000
1   normal  plantation  0.173657  0.043647  0.303667   4.429416  0.004984
2  natural  plantation  0.647718  0.510467  0.784968  15.649601  0.001000


region: 2

    group1      group2      Diff     Lower     Upper    q-value  p-value
0   normal     natural  0.377868  0.330361  0.425375  26.370800    0.001
1   normal  plantation  0.122699  0.061734  0.183665   6.672623    0.001
2  natural  plantation  0.500568  0.423956  0.577180  21.662318    0.001


region: 3

       group1      group2      Diff     Lower     Upper    q-value  p-value
0      normal  plantation  0.086930  0.043456  0.130405   6.629737    0.001
1      normal     natural  0.226985  0.141183  0.312787   8.771208    0.001
2  plantation     natural  0.313915  0.218169  0.409662  10.870534    0.001


region: 4

    group1      group2      Diff     Lower     Upper   q-value   p-value
0  natural      normal  0.092145 -0.191152  0.375442  1.080472  0.706773
1  natural  plantation  0.169699 -0.117206  0.456604  1.964827  0.347866
2   normal  plantation  0.077554  0.026759  0.128349  5.071834  0.001047
```

In [24]:

```
plots.plot_boxplots(
    nfi_df, "natura2000", height=4, aspect=0.75, fig_title="d) Plot-level validation - Natura 2000 habitat quality",
    save_name="Figure4d.png", width=0.5
)
```

In [25]:

```
plots.describe_ci(nfi_df, "pred_mean", ["region", "natura2000"])
```

Out[25]:

|  |  | mean | count | std | ci95\_lo | ci95\_hi |
| --- | --- | --- | --- | --- | --- | --- |
| region | natura2000 |  |  |  |  |  |
| 1 | No | 0.338197 | 2841 | 0.233413 | 0.329614 | 0.346780 |
|  | Yes | 0.793866 | 231 | 0.237824 | 0.763196 | 0.824535 |
| 2 | No | 0.354599 | 5216 | 0.216831 | 0.348714 | 0.360483 |
|  | Yes | 0.785148 | 181 | 0.196554 | 0.756512 | 0.813783 |
| 3 | No | 0.370124 | 4528 | 0.182905 | 0.364796 | 0.375451 |
|  | Yes | 0.654614 | 95 | 0.181591 | 0.618097 | 0.691130 |
| 4 | No | 0.366167 | 660 | 0.167526 | 0.353386 | 0.378948 |
|  | Yes | 0.591048 | 23 | 0.149615 | 0.529902 | 0.652194 |

In [26]:

```
plots.tukey_subsets(nfi_df, "pred_mean", "natura2000", "region")
```

```
region: 1

  group1 group2      Diff     Lower     Upper    q-value  p-value
0     No    Yes  0.455668  0.424312  0.487025  40.294907    0.001


region: 2

  group1 group2      Diff     Lower     Upper    q-value  p-value
0     No    Yes  0.430549  0.398505  0.462592  37.251445    0.001


region: 3

  group1 group2     Diff     Lower     Upper   q-value  p-value
0     No    Yes  0.28449  0.247322  0.321658  21.22133    0.001


region: 4

  group1 group2      Diff     Lower     Upper   q-value  p-value
0     No    Yes  0.224881  0.155338  0.294424  8.979154    0.001
```

In [ ]:
