## Supplementary material for "Mapping forests with different levels of naturalness using machine learning and landscape data mining": Jupyter Notebooks: M4b_onescale_nolatlon.html

validation


In [1]:

```
%load_ext autoreload
%autoreload 2

import warnings
warnings.filterwarnings("ignore")

from matplotlib import pyplot as plt
import matplotlib as mpl
mpl.rcParams["figure.dpi"] = 150

import json
import sys
import numpy as np
import pandas
import seaborn as sns
sns.set_style("white")

sys.path.insert(0, "/app/src")

from plots import plots
from config import settings
```

In [2]:

```
with open("./config.json", "r") as _file:
    mconfig = json.load(_file)
print(json.dumps(mconfig, indent=2))

model_name = mconfig["model_metadata"]["name"]
target_var = mconfig["model_specification"]["target_variable"]
region_var = mconfig["model_specification"]["stratification_layer"]
```

```
{
  "model_metadata": {
    "name": "S1_onescale_nolatlon",
    "description": "One-scale model i.e. only 1ha scale predictors are included. Seperate model for each region. Latitude and longitude excluded from the predictors list."
  },
  "model_data": {
    "samples_layer": "SAMPLES_1"
  },
  "model_specification": {
    "target_variable": "HCVF",
    "stratification_layer": "ecoregions",
    "stratification_submodels": [
      {
        "regions": [
          1
        ],
        "variables_list": []
      },
      {
        "regions": [
          2
        ],
        "variables_list": []
      },
      {
        "regions": [
          3
        ],
        "variables_list": []
      },
      {
        "regions": [
          4
        ],
        "variables_list": []
      }
    ],
    "lat_field": "LAT",
    "lon_field": "LON",
    "target_epsg": 3006,
    "variables_list": [],
    "variables_drop_regexp": "([A-Z]?[0-9]{3})",
    "variables_keep_regexp": "",
    "variables_latlon": false,
    "balanced_rf": true
  },
  "model_crossvalidation": {
    "n_splits": 10,
    "spatial": true,
    "spatial_grid": "grid_20.geojson"
  },
  "model_hyperparameters": {
    "n_estimators": 500,
    "oob_score": true
  },
  "prediction": {
    "predict": true
  }
}
```

In [3]:

```
models = plots.load_models("./", regexp=fr"{model_name}_R[1-5]*.pickle")
models_stats = plots.get_cv_validation_stats(models)
```

```
Loading S1_onescale_nolatlon_R1.pickle...
Loading S1_onescale_nolatlon_R2.pickle...
Loading S1_onescale_nolatlon_R3.pickle...
Loading S1_onescale_nolatlon_R4.pickle...
```

In [4]:

```
print(models[0].data.shape)
models_labels=["North boreal", "South boreal", "Hemiboreal", "Nemoral"]

g = sns.FacetGrid(
    data=models[0].data, col=region_var, aspect=0.7, 
    sharey=False
)
g.map_dataframe(sns.countplot, x=target_var, palette=["black","red"])
axes = g.axes.flatten()
for i, ax in enumerate(axes):
    ax.set_title(models_labels[i], fontweight="bold", fontsize=16)
```

```
(14090, 130)
```

In [5]:

```
plots.plot_validation_curves_multimodel(
    models_stats, colors="bgrcmykw", figsize=(10, 5), plot_std=True, std_alpha=0.1,
    models_labels=["North boreal", "South boreal", "Hemiboreal", "Nemoral"],
    save_name="Figure2.png"
)
```

In [6]:

```
# performance metrics
means, stds = plots.get_metrics(models_stats)
display(means)
display(stds)
```

|  | accuracy | TSS | roc\_auc | pr\_auc | pearson\_r | briers\_score | MCC |
| --- | --- | --- | --- | --- | --- | --- | --- |
| 0 | 0.792805 | 0.586580 | 0.876023 | 0.866604 | 0.651483 | 0.145480 | 0.585209 |
| 1 | 0.798925 | 0.592101 | 0.881210 | 0.846176 | 0.658300 | 0.142251 | 0.591217 |
| 2 | 0.774585 | 0.518373 | 0.849758 | 0.774621 | 0.590361 | 0.161628 | 0.537694 |
| 3 | 0.785869 | 0.545068 | 0.861103 | 0.804252 | 0.613188 | 0.161147 | 0.573026 |

|  | accuracy | TSS | roc\_auc | pr\_auc | pearson\_r | briers\_score | MCC |
| --- | --- | --- | --- | --- | --- | --- | --- |
| 0 | 0.017182 | 0.037697 | 0.014893 | 0.022676 | 0.025179 | 0.007183 | 0.033937 |
| 1 | 0.010536 | 0.028175 | 0.008404 | 0.017131 | 0.015673 | 0.005967 | 0.022485 |
| 2 | 0.019253 | 0.030534 | 0.016944 | 0.033910 | 0.029091 | 0.009012 | 0.030234 |
| 3 | 0.073862 | 0.138241 | 0.054972 | 0.098689 | 0.108795 | 0.029910 | 0.129780 |

In [7]:

```
# number of variables per model
for i, model in enumerate(models):
    print(f"Model_{i+1}: ", len(model.cov_names))
```

```
Model_1:  26
Model_2:  26
Model_3:  26
Model_4:  26
```

In [8]:

```
# plot features Gini's importances
for i, model in enumerate(models):
    print(models_labels[i])
    _plot = plots.plot_gini_importances(model, figsize=(14, 10), n=50)
```

```
North boreal
```

```
South boreal
```

```
Hemiboreal
```

```
Nemoral
```

##### Partial Dependence Plot¶

In [9]:

```
plots.plot_partial_dependences(models, n_features=6)
```

##### Accumulated Local Effects Plot¶

In [10]:

```
plots.plot_ales(models, n_features=6, xlims=(-2, 2))
```

In [11]:

```
# average max tree depth per model
import numpy as np
for i, model in enumerate(models):
    max_depths = []
    for tree_idx, est in enumerate(model.model.estimators_):
        max_depths.append(est.tree_.max_depth)
    print(f"{models_labels[i]}: ", np.array(max_depths).mean())
```

```
North boreal:  22.464
South boreal:  21.728
Hemiboreal:  21.402
Nemoral:  12.576
```

### Validation¶

#### Sveaskog validation¶

tree stand (polygon) level validation; minimum 10 pixels with predicted value

In [12]:

```
# load a dataframe with predictions and Sveaskog atributes
svea_df = pandas.read_csv("svea_predictions.csv")
svea_df = plots.svea_preprocess(svea_df)

# filter out data not reliable for validation
# min number of 1 ha pixels to calculate the mean probability
svea_df = svea_df[svea_df.pred_count > 10]

# show the number of validation polygons per region
display(svea_df.region.value_counts())
print("TOTAL POLYGONS N: ", svea_df.region.value_counts().sum())
```

```
1    41784
2     7989
3     7519
4      256
Name: region, dtype: int64
```

```
TOTAL POLYGONS N:  57548
```

##### Boxplot -> forest management classes¶

"NF" (conservation, no management)
"NF\_NM" (conservation, no yet specified)
"NM" (conservation, management)
"PE" (production forest, increased conservation considerations)
"PG" (production forest, general conservation conciderations)

Management categories for stands. A goal for long-term management is set for each stand, in one of four categories: (i) NF = Nature conservation, free development; (ii) NM = Nature conservation-oriented management, often implying restoration measures; (iii) PG = Production with general conservation concern (green-tree and deadwood retention for biodiversity at harvest, represented by black dots). The minimum level of retention is specified, and always >15 % of the harvested area; (iv) PE = Production with enhanced conservation concern.

In [13]:

```
plots.plot_boxplots(
    svea_df, "management", height=4, aspect=0.75, fig_title="a) Stand-level validation - Forest management objectives",
    save_name="Figure4a.png", width=0.85
)
```

In [14]:

```
plots.describe_ci(svea_df, "pred_mean", ["region", "management"])
```

Out[14]:

|  |  | mean | count | std | ci95\_lo | ci95\_hi |
| --- | --- | --- | --- | --- | --- | --- |
| region | management |  |  |  |  |  |
| 1 | NF | 0.652173 | 4148 | 0.203235 | 0.645988 | 0.658358 |
|  | NF\_NM | 0.714170 | 2401 | 0.172938 | 0.707252 | 0.721087 |
|  | NM | 0.561996 | 903 | 0.205400 | 0.548599 | 0.575393 |
|  | PE | 0.405301 | 2516 | 0.194477 | 0.397702 | 0.412901 |
|  | PG | 0.267868 | 31816 | 0.159930 | 0.266110 | 0.269625 |
| 2 | NF | 0.660822 | 1245 | 0.177235 | 0.650977 | 0.670667 |
|  | NF\_NM | 0.679510 | 82 | 0.188272 | 0.638760 | 0.720261 |
|  | NM | 0.499638 | 253 | 0.200275 | 0.474959 | 0.524317 |
|  | PE | 0.369847 | 868 | 0.198730 | 0.356626 | 0.383068 |
|  | PG | 0.255937 | 5541 | 0.168137 | 0.251510 | 0.260364 |
| 3 | NF | 0.618239 | 297 | 0.156463 | 0.600444 | 0.636034 |
|  | NF\_NM | 0.641031 | 172 | 0.159917 | 0.617131 | 0.664930 |
|  | NM | 0.545287 | 303 | 0.188980 | 0.524008 | 0.566566 |
|  | PE | 0.375628 | 608 | 0.173368 | 0.361847 | 0.389409 |
|  | PG | 0.295678 | 6139 | 0.142632 | 0.292110 | 0.299246 |
| 4 | NF | 0.687365 | 5 | 0.074984 | 0.621639 | 0.753091 |
|  | NF\_NM | NaN | 0 | NaN | NaN | NaN |
|  | NM | 0.594961 | 19 | 0.176732 | 0.515493 | 0.674430 |
|  | PE | 0.485663 | 13 | 0.157698 | 0.399937 | 0.571388 |
|  | PG | 0.280798 | 219 | 0.139456 | 0.262328 | 0.299268 |

In [15]:

```
plots.tukey_subsets(svea_df, "pred_mean", "management", "region")
```

```
region: 1

  group1 group2      Diff     Lower     Upper     q-value  p-value
0     NF  NF_NM  0.061622  0.049693  0.073551   19.928674    0.001
1     NF     PG  0.386143  0.378457  0.393828  193.836962    0.001
2     NF     PE  0.231002  0.218134  0.243869   69.257434    0.001
3     NF     NM  0.091705  0.074594  0.108817   20.675228    0.001
4  NF_NM     PG  0.447765  0.437906  0.457624  175.207927    0.001
5  NF_NM     PE  0.292624  0.278351  0.306897   79.094314    0.001
6  NF_NM     NM  0.153328  0.135136  0.171519   32.515231    0.001
7     PG     PE  0.155141  0.144165  0.166117   54.528566    0.001
8     PG     NM  0.294437  0.278699  0.310176   72.171895    0.001
9     PE     NM  0.139296  0.120476  0.158117   28.552871    0.001


region: 2

  group1 group2      Diff     Lower     Upper     q-value   p-value
0     PG     PE  0.115084  0.095246  0.134921   22.384407  0.001000
1     PG     NF  0.409112  0.393951  0.424274  104.118262  0.001000
2     PG     NM  0.249098  0.218122  0.280075   31.028814  0.001000
3     PG  NF_NM  0.428238  0.374780  0.481695   30.910136  0.001000
4     PE     NF  0.294029  0.270888  0.317170   49.026547  0.001000
5     PE     NM  0.134015  0.098445  0.169584   14.537927  0.001000
6     PE  NF_NM  0.313154  0.256910  0.369398   21.483728  0.001000
7     NF     NM  0.160014  0.126825  0.193203   18.603242  0.001000
8     NF  NF_NM  0.019125 -0.035644  0.073895    1.347396  0.864622
9     NM  NF_NM  0.179139  0.118071  0.240208   11.318769  0.001000


region: 3

  group1 group2      Diff     Lower     Upper    q-value   p-value
0     PE     NF  0.240000  0.210806  0.269194  31.720945  0.001000
1     PE     PG  0.082471  0.064446  0.100496  17.654571  0.001000
2     PE  NF_NM  0.265707  0.230115  0.301299  28.805807  0.001000
3     PE     NM  0.168426  0.139102  0.197749  22.162697  0.001000
4     NF     PG  0.322470  0.298340  0.346601  51.564523  0.001000
5     NF  NF_NM  0.025707 -0.013333  0.064748   2.540778  0.376553
6     NF     NM  0.071574  0.038149  0.104999   8.262435  0.001000
7     PG  NF_NM  0.348178  0.316606  0.379749  42.552950  0.001000
8     PG     NM  0.250897  0.226609  0.275184  39.861095  0.001000
9  NF_NM     NM  0.097281  0.058144  0.136419   9.590995  0.001000


region: 4

  group1 group2      Diff     Lower     Upper    q-value   p-value
0     PG     NM  0.323151  0.234457  0.411846  13.334163  0.001000
1     PG     NF  0.396055  0.209424  0.582685   7.766591  0.001000
2     PG     PE  0.188779  0.074341  0.303218   6.037265  0.001000
3     NM     NF  0.072903 -0.130431  0.276238   1.312177  0.764807
4     NM     PE  0.134372 -0.005665  0.274409   3.511759  0.065313
5     NF     PE  0.207275 -0.008536  0.423086   3.515048  0.064931
```

##### Boxplot -> forest naturalness classes¶

In [16]:

```
plots.plot_boxplots(
    svea_df, "naturalness", height=4, aspect=0.75, fig_title="b) Stand-level validation - Forest naturalness",
    save_name="Figure4b.png", width=0.5
)
```

In [17]:

```
plots.describe_ci(svea_df, "pred_mean", ["region", "naturalness"])
```

Out[17]:

|  |  | mean | count | std | ci95\_lo | ci95\_hi |
| --- | --- | --- | --- | --- | --- | --- |
| region | naturalness |  |  |  |  |  |
| 1 | No | 0.277238 | 34130 | 0.166031 | 0.275477 | 0.279000 |
|  | Yes | 0.654231 | 7654 | 0.204526 | 0.649649 | 0.658813 |
| 2 | No | 0.270184 | 6306 | 0.175930 | 0.265842 | 0.274526 |
|  | Yes | 0.618090 | 1683 | 0.205232 | 0.608285 | 0.627896 |
| 3 | No | 0.302435 | 6602 | 0.147402 | 0.298879 | 0.305991 |
|  | Yes | 0.551767 | 917 | 0.197801 | 0.538964 | 0.564569 |
| 4 | No | 0.293802 | 233 | 0.149407 | 0.274618 | 0.312987 |
|  | Yes | 0.612763 | 23 | 0.167515 | 0.544302 | 0.681224 |

In [18]:

```
plots.tukey_subsets(svea_df, "pred_mean", "naturalness", "region")
```

```
region: 1

  group1 group2      Diff     Lower     Upper     q-value  p-value
0    Yes     No  0.379842  0.375471  0.384213  240.884157    0.001


region: 2

  group1 group2      Diff     Lower     Upper    q-value  p-value
0     No    Yes  0.355412  0.345422  0.365401  98.629763    0.001


region: 3

  group1 group2      Diff     Lower     Upper    q-value  p-value
0     No    Yes  0.250362  0.239571  0.261153  64.320667    0.001


region: 4

  group1 group2      Diff     Lower     Upper  q-value  p-value
0     No    Yes  0.322557  0.256463  0.388651  13.5975    0.001
```

#### NFI validation¶

plot level validation (1 central pixel + 4 neighbours)

In [19]:

```
# load a dataframe with predictions and NFI atributes
nfi_df = pandas.read_csv("nfi_predictions.csv")

# at least 4 pixels needed
nfi_df = nfi_df[nfi_df.pred_count > 3]

nfi_df["region"] = pandas.Categorical(nfi_df.region)
nfi_df["naturalness"] = pandas.Categorical(nfi_df.naturalness, categories=[3, 1, 2])
nfi_df.naturalness.cat.rename_categories({
    1.0: "normal", 2.0: "natural", 3.0: "plantation"
}, inplace=True)

# drop rows with NA in natura2000 column
nfi_df.dropna(subset=["natura2000"], inplace=True)
nfi_df.natura2000.replace({0: "No", 1: "Yes"}, inplace=True)
nfi_df["natura2000"] = pandas.Categorical(nfi_df.natura2000)

# show the number of validation polygons per region
display(nfi_df.region.value_counts())
print("TOTAL PLOTS N: ", nfi_df.region.value_counts().sum())
```

```
2    5397
3    4623
1    3072
4     683
Name: region, dtype: int64
```

```
TOTAL PLOTS N:  13775
```

##### Boxplot -> forest naturalness classes¶

1=normal; 2=natural forest; 3=plantation forest

In [20]:

```
plots.plot_boxplots(
    nfi_df, "naturalness", height=4, aspect=0.75, fig_title="c) Plot-level validation - Forest naturalness",
    save_name="Figure4c.png", width=0.85
)
```

In [21]:

```
plots.describe_ci(nfi_df, "pred_mean", ["region", "naturalness"])
```

Out[21]:

|  |  | mean | count | std | ci95\_lo | ci95\_hi |
| --- | --- | --- | --- | --- | --- | --- |
| region | naturalness |  |  |  |  |  |
| 1 | plantation | 0.182095 | 19 | 0.113188 | 0.131199 | 0.232990 |
|  | normal | 0.347233 | 2897 | 0.244713 | 0.338322 | 0.356145 |
|  | natural | 0.819513 | 156 | 0.175626 | 0.791953 | 0.847074 |
| 2 | plantation | 0.226612 | 74 | 0.136507 | 0.195510 | 0.257715 |
|  | normal | 0.357385 | 5200 | 0.225326 | 0.351260 | 0.363509 |
|  | natural | 0.742810 | 123 | 0.215723 | 0.704686 | 0.780934 |
| 3 | plantation | 0.293743 | 103 | 0.172514 | 0.260426 | 0.327059 |
|  | normal | 0.374429 | 4494 | 0.185231 | 0.369013 | 0.379845 |
|  | natural | 0.586481 | 26 | 0.193417 | 0.512134 | 0.660828 |
| 4 | plantation | 0.302432 | 69 | 0.136566 | 0.270208 | 0.334655 |
|  | normal | 0.381046 | 612 | 0.175226 | 0.367164 | 0.394929 |
|  | natural | 0.511650 | 2 | 0.275560 | 0.129744 | 0.893556 |

In [22]:

```
plots.describe_ci(nfi_df, "pred_mean", ["region", "naturalness"])["count"].sum()
```

Out[22]:

```
13775
```

In [23]:

```
plots.tukey_subsets(nfi_df, "pred_mean", "naturalness", "region")
```

```
region: 1

    group1      group2      Diff     Lower     Upper    q-value  p-value
0   normal     natural  0.472280  0.425808  0.518752  33.701027  0.00100
1   normal  plantation  0.165139  0.035001  0.295277   4.208004  0.00829
2  natural  plantation  0.637419  0.500033  0.774804  15.385627  0.00100


region: 2

    group1      group2      Diff     Lower     Upper    q-value  p-value
0   normal     natural  0.385425  0.337487  0.433363  26.656372    0.001
1   normal  plantation  0.130772  0.069254  0.192291   7.047720    0.001
2  natural  plantation  0.516198  0.438891  0.593505  22.137892    0.001


region: 3

       group1      group2      Diff     Lower     Upper    q-value  p-value
0      normal  plantation  0.080686  0.037462  0.123911   6.189106    0.001
1      normal     natural  0.212052  0.126743  0.297361   8.241518    0.001
2  plantation     natural  0.292738  0.197542  0.387934  10.195785    0.001


region: 4

    group1      group2      Diff     Lower     Upper   q-value   p-value
0  natural      normal  0.130604 -0.155454  0.416661  1.516649  0.530318
1  natural  plantation  0.209218 -0.080482  0.498918  2.399015  0.207531
2   normal  plantation  0.078615  0.027325  0.129905  5.091582  0.001000
```

In [24]:

```
plots.plot_boxplots(
    nfi_df, "natura2000", height=4, aspect=0.75, fig_title="d) Plot-level validation - Natura 2000 habitat quality",
    save_name="Figure4d.png", width=0.5
)
```

In [25]:

```
plots.describe_ci(nfi_df, "pred_mean", ["region", "natura2000"])
```

Out[25]:

|  |  | mean | count | std | ci95\_lo | ci95\_hi |
| --- | --- | --- | --- | --- | --- | --- |
| region | natura2000 |  |  |  |  |  |
| 1 | No | 0.336033 | 2841 | 0.233729 | 0.327438 | 0.344628 |
|  | Yes | 0.790339 | 231 | 0.236759 | 0.759807 | 0.820871 |
| 2 | No | 0.349812 | 5216 | 0.219050 | 0.343867 | 0.355756 |
|  | Yes | 0.784070 | 181 | 0.197999 | 0.755224 | 0.812915 |
| 3 | No | 0.368089 | 4528 | 0.181779 | 0.362794 | 0.373383 |
|  | Yes | 0.647181 | 95 | 0.182476 | 0.610487 | 0.683875 |
| 4 | No | 0.366143 | 660 | 0.169700 | 0.353196 | 0.379090 |
|  | Yes | 0.584213 | 23 | 0.149394 | 0.523158 | 0.645269 |

In [26]:

```
plots.tukey_subsets(nfi_df, "pred_mean", "natura2000", "region")
```

```
region: 1

  group1 group2      Diff     Lower     Upper    q-value  p-value
0     No    Yes  0.454306  0.422921  0.485691  40.138201    0.001


region: 2

  group1 group2      Diff     Lower     Upper    q-value  p-value
0     No    Yes  0.434258  0.401889  0.466627  37.194759    0.001


region: 3

  group1 group2      Diff     Lower    Upper    q-value  p-value
0     No    Yes  0.279092  0.242145  0.31604  20.942964    0.001


region: 4

  group1 group2     Diff    Lower     Upper   q-value  p-value
0     No    Yes  0.21807  0.14765  0.288489  8.598783    0.001
```

In [ ]:
