## Supplementary material for "Mapping forests with different levels of naturalness using machine learning and landscape data mining": Jupyter Notebooks: M5a_baseline.html

validation


In [5]:

```
%load_ext autoreload
%autoreload 2

import warnings
warnings.filterwarnings("ignore")

from matplotlib import pyplot as plt
import matplotlib as mpl
mpl.rcParams["figure.dpi"] = 150

import json
import sys
import numpy as np
import pandas
import seaborn as sns
sns.set_style("white")

sys.path.insert(0, "/app/src")

from plots import plots
from config import settings
```

```
The autoreload extension is already loaded. To reload it, use:
  %reload_ext autoreload
```

In [6]:

```
with open("./config.json", "r") as _file:
    mconfig = json.load(_file)
print(json.dumps(mconfig, indent=2))

model_name = mconfig["model_metadata"]["name"]
target_var = mconfig["model_specification"]["target_variable"]
region_var = mconfig["model_specification"]["stratification_layer"]
```

```
{
  "model_metadata": {
    "name": "S1_baseline",
    "description": "Baseline / reference model. Seperate model for each region."
  },
  "model_data": {
    "samples_layer": "SAMPLES_1"
  },
  "model_specification": {
    "target_variable": "HCVF",
    "stratification_layer": "ecoregions",
    "stratification_submodels": [
      {
        "regions": [
          1
        ],
        "variables_list": []
      },
      {
        "regions": [
          2
        ],
        "variables_list": []
      },
      {
        "regions": [
          3
        ],
        "variables_list": []
      },
      {
        "regions": [
          4
        ],
        "variables_list": []
      }
    ],
    "lat_field": "LAT",
    "lon_field": "LON",
    "target_epsg": 3006,
    "variables_list": [
      "DEM",
      "HEIGHTc",
      "HANSEN",
      "ROADSd"
    ],
    "variables_drop_regexp": "",
    "variables_keep_regexp": "",
    "variables_latlon": true,
    "balanced_rf": true
  },
  "model_crossvalidation": {
    "n_splits": 10,
    "spatial": true,
    "spatial_grid": "grid_20.geojson"
  },
  "model_hyperparameters": {
    "n_estimators": 500,
    "oob_score": true
  },
  "prediction": {
    "predict": true
  }
}
```

In [7]:

```
models = plots.load_models("./", regexp=fr"{model_name}_R[1-5]*.pickle")
models_stats = plots.get_cv_validation_stats(models)
```

```
Loading S1_baseline_R1.pickle...
Loading S1_baseline_R2.pickle...
Loading S1_baseline_R3.pickle...
Loading S1_baseline_R4.pickle...
```

In [8]:

```
print(models[0].data.shape)
models_labels=["North boreal", "South boreal", "Hemiboreal", "Nemoral"]

g = sns.FacetGrid(
    data=models[0].data, col=region_var, aspect=0.7, 
    sharey=False
)
g.map_dataframe(sns.countplot, x=target_var, palette=["black","red"])
axes = g.axes.flatten()
for i, ax in enumerate(axes):
    ax.set_title(models_labels[i], fontweight="bold", fontsize=16)
```

```
(14090, 130)
```

In [9]:

```
plots.plot_validation_curves_multimodel(
    models_stats, colors="bgrcmykw", figsize=(10, 5), plot_std=True, std_alpha=0.1,
    models_labels=["North boreal", "South boreal", "Hemiboreal", "Nemoral"],
    save_name="Figure2.png"
)
```

In [10]:

```
# performance metrics
means, stds = plots.get_metrics(models_stats)
display(means)
display(stds)
```

|  | accuracy | TSS | roc\_auc | pr\_auc | pearson\_r | briers\_score | MCC |
| --- | --- | --- | --- | --- | --- | --- | --- |
| 0 | 0.758733 | 0.507383 | 0.838740 | 0.831429 | 0.588221 | 0.162195 | 0.513251 |
| 1 | 0.749884 | 0.496724 | 0.829487 | 0.794658 | 0.568235 | 0.167663 | 0.492306 |
| 2 | 0.707342 | 0.403525 | 0.780570 | 0.679091 | 0.472400 | 0.191555 | 0.407270 |
| 3 | 0.656002 | 0.246195 | 0.716099 | 0.566226 | 0.353802 | 0.220035 | 0.324518 |

|  | accuracy | TSS | roc\_auc | pr\_auc | pearson\_r | briers\_score | MCC |
| --- | --- | --- | --- | --- | --- | --- | --- |
| 0 | 0.024808 | 0.043078 | 0.022603 | 0.050957 | 0.041590 | 0.012556 | 0.049505 |
| 1 | 0.020970 | 0.039207 | 0.018816 | 0.039804 | 0.035716 | 0.008836 | 0.043492 |
| 2 | 0.014840 | 0.029297 | 0.020209 | 0.047123 | 0.035534 | 0.008206 | 0.031082 |
| 3 | 0.060203 | 0.132261 | 0.092978 | 0.096870 | 0.144961 | 0.027487 | 0.139413 |

In [11]:

```
# number of variables per model
for i, model in enumerate(models):
    print(f"Model_{i+1}: ", len(model.cov_names))
```

```
Model_1:  6
Model_2:  6
Model_3:  6
Model_4:  6
```

In [31]:

```
# plot features Gini's importances
for i, model in enumerate(models):
    print(models_labels[i])
    _plot = plots.plot_gini_importances(model, figsize=(14, 4), n=50)
```

```
North boreal
```

```
South boreal
```

```
Hemiboreal
```

```
Nemoral
```

##### Partial Dependence Plot¶

In [13]:

```
plots.plot_partial_dependences(models, n_features=6)
```

##### Accumulated Local Effects Plot¶

In [14]:

```
plots.plot_ales(models, n_features=6, xlims=(-2, 2))
```

In [15]:

```
# average max tree depth per model
import numpy as np
for i, model in enumerate(models):
    max_depths = []
    for tree_idx, est in enumerate(model.model.estimators_):
        max_depths.append(est.tree_.max_depth)
    print(f"{models_labels[i]}: ", np.array(max_depths).mean())
```

```
North boreal:  24.914
South boreal:  24.262
Hemiboreal:  22.936
Nemoral:  15.96
```

### Validation¶

#### Sveaskog validation¶

tree stand (polygon) level validation; minimum 10 pixels with predicted value

In [16]:

```
# load a dataframe with predictions and Sveaskog atributes
svea_df = pandas.read_csv("svea_predictions.csv")
svea_df = plots.svea_preprocess(svea_df)

# filter out data not reliable for validation
# min number of 1 ha pixels to calculate the mean probability
svea_df = svea_df[svea_df.pred_count > 10]

# show the number of validation polygons per region
display(svea_df.region.value_counts())
print("TOTAL POLYGONS N: ", svea_df.region.value_counts().sum())
```

```
1    41790
2     7989
3     7523
4      257
Name: region, dtype: int64
```

```
TOTAL POLYGONS N:  57559
```

##### Boxplot -> forest management classes¶

"NF" (conservation, no management)
"NF\_NM" (conservation, no yet specified)
"NM" (conservation, management)
"PE" (production forest, increased conservation considerations)
"PG" (production forest, general conservation conciderations)

Management categories for stands. A goal for long-term management is set for each stand, in one of four categories: (i) NF = Nature conservation, free development; (ii) NM = Nature conservation-oriented management, often implying restoration measures; (iii) PG = Production with general conservation concern (green-tree and deadwood retention for biodiversity at harvest, represented by black dots). The minimum level of retention is specified, and always >15 % of the harvested area; (iv) PE = Production with enhanced conservation concern.

In [17]:

```
plots.plot_boxplots(
    svea_df, "management", height=4, aspect=0.75, fig_title="a) Stand-level validation - Forest management objectives",
    save_name="Figure4a.png", width=0.85
)
```

In [18]:

```
plots.describe_ci(svea_df, "pred_mean", ["region", "management"])
```

Out[18]:

|  |  | mean | count | std | ci95\_lo | ci95\_hi |
| --- | --- | --- | --- | --- | --- | --- |
| region | management |  |  |  |  |  |
| 1 | NF | 0.603532 | 4148 | 0.209488 | 0.597156 | 0.609907 |
|  | NF\_NM | 0.656876 | 2401 | 0.188957 | 0.649317 | 0.664434 |
|  | NM | 0.513098 | 903 | 0.193634 | 0.500469 | 0.525728 |
|  | PE | 0.381215 | 2516 | 0.204412 | 0.373228 | 0.389203 |
|  | PG | 0.293689 | 31822 | 0.174339 | 0.291774 | 0.295605 |
| 2 | NF | 0.671565 | 1245 | 0.195849 | 0.660686 | 0.682444 |
|  | NF\_NM | 0.635770 | 82 | 0.218305 | 0.588519 | 0.683021 |
|  | NM | 0.523783 | 253 | 0.221347 | 0.496508 | 0.551059 |
|  | PE | 0.392012 | 868 | 0.211582 | 0.377936 | 0.406088 |
|  | PG | 0.288242 | 5541 | 0.181623 | 0.283460 | 0.293025 |
| 3 | NF | 0.622095 | 297 | 0.162554 | 0.603607 | 0.640582 |
|  | NF\_NM | 0.630105 | 172 | 0.166870 | 0.605167 | 0.655044 |
|  | NM | 0.509995 | 303 | 0.202630 | 0.487179 | 0.532811 |
|  | PE | 0.392014 | 608 | 0.174248 | 0.378163 | 0.405865 |
|  | PG | 0.314794 | 6143 | 0.149836 | 0.311047 | 0.318541 |
| 4 | NF | 0.534114 | 5 | 0.208575 | 0.351290 | 0.716938 |
|  | NF\_NM | NaN | 0 | NaN | NaN | NaN |
|  | NM | 0.415651 | 19 | 0.152342 | 0.347150 | 0.484152 |
|  | PE | 0.458247 | 13 | 0.212342 | 0.342816 | 0.573677 |
|  | PG | 0.287514 | 220 | 0.142700 | 0.268657 | 0.306371 |

In [19]:

```
plots.tukey_subsets(svea_df, "pred_mean", "management", "region")
```

```
region: 1

  group1 group2      Diff     Lower     Upper     q-value  p-value
0     NF  NF_NM  0.053101  0.040309  0.065894   16.013633    0.001
1     NF     PG  0.309306  0.301064  0.317547  144.786103    0.001
2     NF     PE  0.190706  0.176907  0.204504   53.316354    0.001
3     NF     NM  0.092891  0.074541  0.111241   19.528689    0.001
4  NF_NM     PG  0.362407  0.351834  0.372980  132.235626    0.001
5  NF_NM     PE  0.243807  0.228501  0.259113   61.450688    0.001
6  NF_NM     NM  0.145992  0.126483  0.165501   28.869567    0.001
7     PG     PE  0.118600  0.106830  0.130371   38.871472    0.001
8     PG     NM  0.216415  0.199537  0.233293   49.466352    0.001
9     PE     NM  0.097815  0.077632  0.117998   18.696542    0.001


region: 2

  group1 group2      Diff     Lower     Upper    q-value  p-value
0     PG     PE  0.120004  0.098482  0.141526  21.515094  0.00100
1     PG     NF  0.384749  0.368301  0.401198  90.256048  0.00100
2     PG     NM  0.237543  0.203937  0.271149  27.274118  0.00100
3     PG  NF_NM  0.349112  0.291116  0.407108  23.227104  0.00100
4     PE     NF  0.264745  0.239640  0.289851  40.689596  0.00100
5     PE     NM  0.117539  0.078950  0.156128  11.752894  0.00100
6     PE  NF_NM  0.229108  0.168089  0.290126  14.487903  0.00100
7     NF     NM  0.147206  0.111200  0.183213  15.775067  0.00100
8     NF  NF_NM  0.035637 -0.023782  0.095056   2.314227  0.47495
9     NM  NF_NM  0.111569  0.045316  0.177822   6.497797  0.00100


region: 3

  group1 group2      Diff     Lower     Upper    q-value  p-value
0     PE     NF  0.221616  0.191197  0.252035  28.111526    0.001
1     PE     PG  0.084223  0.065443  0.103004  17.304129    0.001
2     PE  NF_NM  0.233187  0.196102  0.270273  24.262214    0.001
3     PE     NM  0.110501  0.079947  0.141054  13.954912    0.001
4     NF     PG  0.305839  0.280696  0.330982  46.936330    0.001
5     NF  NF_NM  0.011572 -0.029107  0.052250   1.097618    0.900
6     NF     NM  0.111115  0.076287  0.145943  12.310474    0.001
7     PG  NF_NM  0.317411  0.284514  0.350307  37.230807    0.001
8     PG     NM  0.194724  0.169418  0.220030  29.691279    0.001
9  NF_NM     NM  0.122687  0.081907  0.163467  11.608624    0.001


region: 4

  group1 group2      Diff     Lower     Upper   q-value   p-value
0     PG     NM  0.136730  0.045734  0.227725  5.499056  0.001000
1     PG     NF  0.301879  0.110377  0.493381  5.769028  0.001000
2     PG     PE  0.142188  0.024771  0.259605  4.431772  0.010437
3     NM     NF  0.165149 -0.043503  0.373802  2.896653  0.173703
4     NM     PE  0.005458 -0.138241  0.149158  0.139010  0.900000
5     NF     PE  0.159691 -0.061764  0.381146  2.638991  0.245622
```

##### Boxplot -> forest naturalness classes¶

In [20]:

```
plots.plot_boxplots(
    svea_df, "naturalness", height=4, aspect=0.75, fig_title="b) Stand-level validation - Forest naturalness",
    save_name="Figure4b.png", width=0.5
)
```

In [21]:

```
plots.describe_ci(svea_df, "pred_mean", ["region", "naturalness"])
```

Out[21]:

|  |  | mean | count | std | ci95\_lo | ci95\_hi |
| --- | --- | --- | --- | --- | --- | --- |
| region | naturalness |  |  |  |  |  |
| 1 | No | 0.299400 | 34136 | 0.177667 | 0.297515 | 0.301285 |
|  | Yes | 0.604720 | 7654 | 0.208397 | 0.600052 | 0.609389 |
| 2 | No | 0.300892 | 6306 | 0.188413 | 0.296242 | 0.305543 |
|  | Yes | 0.630267 | 1683 | 0.218076 | 0.619848 | 0.640686 |
| 3 | No | 0.320647 | 6606 | 0.153904 | 0.316935 | 0.324358 |
|  | Yes | 0.546999 | 917 | 0.196876 | 0.534257 | 0.559742 |
| 4 | No | 0.299522 | 234 | 0.156333 | 0.279491 | 0.319553 |
|  | Yes | 0.421302 | 23 | 0.142610 | 0.363019 | 0.479585 |

In [22]:

```
plots.tukey_subsets(svea_df, "pred_mean", "naturalness", "region")
```

```
region: 1

  group1 group2      Diff     Lower     Upper     q-value  p-value
0    Yes     No  0.304099  0.299473  0.308726  182.195517    0.001


region: 2

  group1 group2      Diff     Lower     Upper    q-value  p-value
0     No    Yes  0.332042  0.321315  0.342769  85.811609    0.001


region: 3

  group1 group2      Diff     Lower     Upper    q-value  p-value
0     No    Yes  0.225955  0.214827  0.237083  56.292869    0.001


region: 4

  group1 group2     Diff    Lower    Upper   q-value  p-value
0     No    Yes  0.13568  0.06847  0.20289  5.624558    0.001
```

#### NFI validation¶

plot level validation (1 central pixel + 4 neighbours)

In [23]:

```
# load a dataframe with predictions and NFI atributes
nfi_df = pandas.read_csv("nfi_predictions.csv")

# at least 4 pixels needed
nfi_df = nfi_df[nfi_df.pred_count > 3]

nfi_df["region"] = pandas.Categorical(nfi_df.region)
nfi_df["naturalness"] = pandas.Categorical(nfi_df.naturalness, categories=[3, 1, 2])
nfi_df.naturalness.cat.rename_categories({
    1.0: "normal", 2.0: "natural", 3.0: "plantation"
}, inplace=True)

# drop rows with NA in natura2000 column
nfi_df.dropna(subset=["natura2000"], inplace=True)
nfi_df.natura2000.replace({0: "No", 1: "Yes"}, inplace=True)
nfi_df["natura2000"] = pandas.Categorical(nfi_df.natura2000)

# show the number of validation polygons per region
display(nfi_df.region.value_counts())
print("TOTAL PLOTS N: ", nfi_df.region.value_counts().sum())
```

```
2    5397
3    4627
1    3073
4     683
Name: region, dtype: int64
```

```
TOTAL PLOTS N:  13780
```

##### Boxplot -> forest naturalness classes¶

1=normal; 2=natural forest; 3=plantation forest

In [24]:

```
plots.plot_boxplots(
    nfi_df, "naturalness", height=4, aspect=0.75, fig_title="c) Plot-level validation - Forest naturalness",
    save_name="Figure4c.png", width=0.85
)
```

In [25]:

```
plots.describe_ci(nfi_df, "pred_mean", ["region", "naturalness"])
```

Out[25]:

|  |  | mean | count | std | ci95\_lo | ci95\_hi |
| --- | --- | --- | --- | --- | --- | --- |
| region | naturalness |  |  |  |  |  |
| 1 | plantation | 0.175405 | 19 | 0.106305 | 0.127604 | 0.223206 |
|  | normal | 0.367780 | 2898 | 0.247688 | 0.358762 | 0.376798 |
|  | natural | 0.813938 | 156 | 0.184962 | 0.784913 | 0.842963 |
| 2 | plantation | 0.284786 | 74 | 0.169639 | 0.246135 | 0.323438 |
|  | normal | 0.377408 | 5200 | 0.229607 | 0.371167 | 0.383649 |
|  | natural | 0.720022 | 123 | 0.249374 | 0.675951 | 0.764093 |
| 3 | plantation | 0.312041 | 103 | 0.173627 | 0.278509 | 0.345572 |
|  | normal | 0.408213 | 4498 | 0.190566 | 0.402644 | 0.413782 |
|  | natural | 0.566004 | 26 | 0.178829 | 0.497264 | 0.634743 |
| 4 | plantation | 0.360988 | 69 | 0.171694 | 0.320476 | 0.401501 |
|  | normal | 0.416712 | 612 | 0.166273 | 0.403538 | 0.429885 |
|  | natural | 0.556400 | 2 | 0.237022 | 0.227904 | 0.884896 |

In [26]:

```
plots.describe_ci(nfi_df, "pred_mean", ["region", "naturalness"])["count"].sum()
```

Out[26]:

```
13780
```

In [27]:

```
plots.tukey_subsets(nfi_df, "pred_mean", "naturalness", "region")
```

```
region: 1

    group1      group2      Diff     Lower     Upper    q-value   p-value
0   normal     natural  0.446157  0.399073  0.493242  31.422843  0.001000
1   normal  plantation  0.192375  0.060521  0.324229   4.838235  0.001828
2  natural  plantation  0.638533  0.499335  0.777730  15.211900  0.001000


region: 2

    group1      group2      Diff     Lower     Upper    q-value   p-value
0   normal     natural  0.342614  0.293559  0.391669  23.156000  0.001000
1   normal  plantation  0.092621  0.029669  0.155573   4.877997  0.001642
2  natural  plantation  0.435235  0.356127  0.514344  18.240722  0.001000


region: 3

       group1      group2      Diff     Lower     Upper   q-value  p-value
0      normal  plantation  0.096172  0.051747  0.140597  7.177559    0.001
1      normal     natural  0.157791  0.070112  0.245471  5.966835    0.001
2  plantation     natural  0.253963  0.156122  0.351805  8.606118    0.001


region: 4

    group1      group2      Diff     Lower     Upper   q-value   p-value
0  natural      normal  0.139688 -0.138055  0.417432  1.670700  0.466533
1  natural  plantation  0.195412 -0.085869  0.476692  2.307772  0.233239
2   normal  plantation  0.055723  0.005924  0.105523  3.717031  0.023809
```

In [28]:

```
plots.plot_boxplots(
    nfi_df, "natura2000", height=4, aspect=0.75, fig_title="d) Plot-level validation - Natura 2000 habitat quality",
    save_name="Figure4d.png", width=0.5
)
```

In [29]:

```
plots.describe_ci(nfi_df, "pred_mean", ["region", "natura2000"])
```

Out[29]:

|  |  | mean | count | std | ci95\_lo | ci95\_hi |
| --- | --- | --- | --- | --- | --- | --- |
| region | natura2000 |  |  |  |  |  |
| 1 | No | 0.356223 | 2842 | 0.236784 | 0.347517 | 0.364928 |
|  | Yes | 0.795454 | 231 | 0.238426 | 0.764707 | 0.826201 |
| 2 | No | 0.369342 | 5216 | 0.222473 | 0.363304 | 0.375379 |
|  | Yes | 0.804819 | 181 | 0.201824 | 0.775416 | 0.834222 |
| 3 | No | 0.401651 | 4531 | 0.187275 | 0.396198 | 0.407104 |
|  | Yes | 0.657455 | 96 | 0.198668 | 0.617713 | 0.697197 |
| 4 | No | 0.404318 | 660 | 0.163395 | 0.391852 | 0.416784 |
|  | Yes | 0.617326 | 23 | 0.163076 | 0.550679 | 0.683973 |

In [30]:

```
plots.tukey_subsets(nfi_df, "pred_mean", "natura2000", "region")
```

```
region: 1

  group1 group2      Diff     Lower     Upper    q-value  p-value
0     No    Yes  0.439232  0.407451  0.471012  38.323656    0.001


region: 2

  group1 group2      Diff     Lower     Upper   q-value  p-value
0     No    Yes  0.435477  0.402599  0.468355  36.72153    0.001


region: 3

  group1 group2      Diff     Lower    Upper  q-value  p-value
0     No    Yes  0.255804  0.217889  0.29372  18.7054    0.001


region: 4

  group1 group2      Diff     Lower     Upper   q-value  p-value
0     No    Yes  0.213008  0.144961  0.281055  8.692073    0.001
```

In [ ]:
