## Supplementary material for "Mapping forests with different levels of naturalness using machine learning and landscape data mining": Jupyter Notebooks: M5b_baseline_nolatlon.html

validation


In [1]:

```
%load_ext autoreload
%autoreload 2

import warnings
warnings.filterwarnings("ignore")

from matplotlib import pyplot as plt
import matplotlib as mpl
mpl.rcParams["figure.dpi"] = 150

import json
import sys
import numpy as np
import pandas
import seaborn as sns
sns.set_style("white")

sys.path.insert(0, "/app/src")

from plots import plots
from config import settings
```

In [2]:

```
with open("./config.json", "r") as _file:
    mconfig = json.load(_file)
print(json.dumps(mconfig, indent=2))

model_name = mconfig["model_metadata"]["name"]
target_var = mconfig["model_specification"]["target_variable"]
region_var = mconfig["model_specification"]["stratification_layer"]
```

```
{
  "model_metadata": {
    "name": "S1_baseline_nolatlon",
    "description": "Baseline / reference model. Seperate model for each region. Latitude and longitude excluded from the predictors list."
  },
  "model_data": {
    "samples_layer": "SAMPLES_1"
  },
  "model_specification": {
    "target_variable": "HCVF",
    "stratification_layer": "ecoregions",
    "stratification_submodels": [
      {
        "regions": [
          1
        ],
        "variables_list": []
      },
      {
        "regions": [
          2
        ],
        "variables_list": []
      },
      {
        "regions": [
          3
        ],
        "variables_list": []
      },
      {
        "regions": [
          4
        ],
        "variables_list": []
      }
    ],
    "lat_field": "LAT",
    "lon_field": "LON",
    "target_epsg": 3006,
    "variables_list": [
      "DEM",
      "HEIGHTc",
      "HANSEN",
      "ROADSd"
    ],
    "variables_drop_regexp": "",
    "variables_keep_regexp": "",
    "variables_latlon": false,
    "balanced_rf": true
  },
  "model_crossvalidation": {
    "n_splits": 10,
    "spatial": true,
    "spatial_grid": "grid_20.geojson"
  },
  "model_hyperparameters": {
    "n_estimators": 500,
    "oob_score": true
  },
  "prediction": {
    "predict": true
  }
}
```

In [3]:

```
models = plots.load_models("./", regexp=fr"{model_name}_R[1-5]*.pickle")
models_stats = plots.get_cv_validation_stats(models)
```

```
Loading S1_baseline_nolatlon_R1.pickle...
Loading S1_baseline_nolatlon_R2.pickle...
Loading S1_baseline_nolatlon_R3.pickle...
Loading S1_baseline_nolatlon_R4.pickle...
```

In [4]:

```
print(models[0].data.shape)
models_labels=["North boreal", "South boreal", "Hemiboreal", "Nemoral"]

g = sns.FacetGrid(
    data=models[0].data, col=region_var, aspect=0.7, 
    sharey=False
)
g.map_dataframe(sns.countplot, x=target_var, palette=["black","red"])
axes = g.axes.flatten()
for i, ax in enumerate(axes):
    ax.set_title(models_labels[i], fontweight="bold", fontsize=16)
```

```
(14090, 130)
```

In [5]:

```
plots.plot_validation_curves_multimodel(
    models_stats, colors="bgrcmykw", figsize=(10, 5), plot_std=True, std_alpha=0.1,
    models_labels=["North boreal", "South boreal", "Hemiboreal", "Nemoral"],
    save_name="Figure2.png"
)
```

In [6]:

```
# performance metrics
means, stds = plots.get_metrics(models_stats)
display(means)
display(stds)
```

|  | accuracy | TSS | roc\_auc | pr\_auc | pearson\_r | briers\_score | MCC |
| --- | --- | --- | --- | --- | --- | --- | --- |
| 0 | 0.719342 | 0.424074 | 0.784234 | 0.751526 | 0.492242 | 0.191159 | 0.436606 |
| 1 | 0.730658 | 0.453347 | 0.808004 | 0.768267 | 0.529964 | 0.180363 | 0.456175 |
| 2 | 0.676426 | 0.357611 | 0.739874 | 0.627357 | 0.404748 | 0.211267 | 0.347248 |
| 3 | 0.628921 | 0.187592 | 0.685096 | 0.575033 | 0.319815 | 0.229578 | 0.258398 |

|  | accuracy | TSS | roc\_auc | pr\_auc | pearson\_r | briers\_score | MCC |
| --- | --- | --- | --- | --- | --- | --- | --- |
| 0 | 0.019820 | 0.030378 | 0.016093 | 0.032976 | 0.028107 | 0.008301 | 0.040276 |
| 1 | 0.026597 | 0.049623 | 0.027361 | 0.037897 | 0.047913 | 0.013855 | 0.054207 |
| 2 | 0.026006 | 0.056795 | 0.027466 | 0.028351 | 0.044690 | 0.013076 | 0.050459 |
| 3 | 0.069362 | 0.133284 | 0.070467 | 0.085808 | 0.112087 | 0.031562 | 0.161465 |

In [7]:

```
# number of variables per model
for i, model in enumerate(models):
    print(f"Model_{i+1}: ", len(model.cov_names))
```

```
Model_1:  4
Model_2:  4
Model_3:  4
Model_4:  4
```

In [8]:

```
# plot features Gini's importances
for i, model in enumerate(models):
    print(models_labels[i])
    _plot = plots.plot_gini_importances(model, figsize=(14, 4), n=50)
```

```
North boreal
```

```
South boreal
```

```
Hemiboreal
```

```
Nemoral
```

##### Partial Dependence Plot¶

In [10]:

```
plots.plot_partial_dependences(models, n_features=4)
```

##### Accumulated Local Effects Plot¶

In [11]:

```
plots.plot_ales(models, n_features=4, xlims=(-2, 2))
```

In [12]:

```
# average max tree depth per model
import numpy as np
for i, model in enumerate(models):
    max_depths = []
    for tree_idx, est in enumerate(model.model.estimators_):
        max_depths.append(est.tree_.max_depth)
    print(f"{models_labels[i]}: ", np.array(max_depths).mean())
```

```
North boreal:  27.182
South boreal:  25.366
Hemiboreal:  24.526
Nemoral:  17.08
```

### Validation¶

#### Sveaskog validation¶

tree stand (polygon) level validation; minimum 10 pixels with predicted value

In [13]:

```
# load a dataframe with predictions and Sveaskog atributes
svea_df = pandas.read_csv("svea_predictions.csv")
svea_df = plots.svea_preprocess(svea_df)

# filter out data not reliable for validation
# min number of 1 ha pixels to calculate the mean probability
svea_df = svea_df[svea_df.pred_count > 10]

# show the number of validation polygons per region
display(svea_df.region.value_counts())
print("TOTAL POLYGONS N: ", svea_df.region.value_counts().sum())
```

```
1    41790
2     7989
3     7523
4      257
Name: region, dtype: int64
```

```
TOTAL POLYGONS N:  57559
```

##### Boxplot -> forest management classes¶

"NF" (conservation, no management)
"NF\_NM" (conservation, no yet specified)
"NM" (conservation, management)
"PE" (production forest, increased conservation considerations)
"PG" (production forest, general conservation conciderations)

Management categories for stands. A goal for long-term management is set for each stand, in one of four categories: (i) NF = Nature conservation, free development; (ii) NM = Nature conservation-oriented management, often implying restoration measures; (iii) PG = Production with general conservation concern (green-tree and deadwood retention for biodiversity at harvest, represented by black dots). The minimum level of retention is specified, and always >15 % of the harvested area; (iv) PE = Production with enhanced conservation concern.

In [14]:

```
plots.plot_boxplots(
    svea_df, "management", height=4, aspect=0.75, fig_title="a) Stand-level validation - Forest management objectives",
    save_name="Figure4a.png", width=0.85
)
```

In [15]:

```
plots.describe_ci(svea_df, "pred_mean", ["region", "management"])
```

Out[15]:

|  |  | mean | count | std | ci95\_lo | ci95\_hi |
| --- | --- | --- | --- | --- | --- | --- |
| region | management |  |  |  |  |  |
| 1 | NF | 0.567207 | 4148 | 0.204781 | 0.560975 | 0.573439 |
|  | NF\_NM | 0.626731 | 2401 | 0.187706 | 0.619223 | 0.634240 |
|  | NM | 0.491432 | 903 | 0.191207 | 0.478961 | 0.503904 |
|  | PE | 0.374847 | 2516 | 0.203651 | 0.366889 | 0.382804 |
|  | PG | 0.294014 | 31822 | 0.173160 | 0.292111 | 0.295916 |
| 2 | NF | 0.683950 | 1245 | 0.200954 | 0.672788 | 0.695113 |
|  | NF\_NM | 0.634497 | 82 | 0.224056 | 0.586001 | 0.682993 |
|  | NM | 0.519978 | 253 | 0.224147 | 0.492358 | 0.547599 |
|  | PE | 0.383584 | 868 | 0.221763 | 0.368831 | 0.398337 |
|  | PG | 0.288584 | 5541 | 0.193275 | 0.283495 | 0.293673 |
| 3 | NF | 0.593149 | 297 | 0.167216 | 0.574131 | 0.612166 |
|  | NF\_NM | 0.597059 | 172 | 0.180644 | 0.570062 | 0.624056 |
|  | NM | 0.474110 | 303 | 0.207479 | 0.450748 | 0.497472 |
|  | PE | 0.364983 | 608 | 0.182120 | 0.350506 | 0.379459 |
|  | PG | 0.296265 | 6143 | 0.148875 | 0.292542 | 0.299988 |
| 4 | NF | 0.559931 | 5 | 0.151996 | 0.426701 | 0.693161 |
|  | NF\_NM | NaN | 0 | NaN | NaN | NaN |
|  | NM | 0.421919 | 19 | 0.155889 | 0.351823 | 0.492016 |
|  | PE | 0.446371 | 13 | 0.195824 | 0.339920 | 0.552822 |
|  | PG | 0.298379 | 220 | 0.152890 | 0.278176 | 0.318583 |

In [16]:

```
plots.tukey_subsets(svea_df, "pred_mean", "management", "region")
```

```
region: 1

  group1 group2      Diff     Lower     Upper     q-value  p-value
0     NF  NF_NM  0.059080  0.046432  0.071727   18.020629    0.001
1     NF     PG  0.271992  0.263844  0.280140  128.777218    0.001
2     NF     PE  0.157393  0.143750  0.171035   44.506809    0.001
3     NF     NM  0.077416  0.059273  0.095558   16.461702    0.001
4  NF_NM     PG  0.331072  0.320618  0.341525  122.185144    0.001
5  NF_NM     PE  0.216473  0.201340  0.231605   55.185968    0.001
6  NF_NM     NM  0.136496  0.117208  0.155783   27.300771    0.001
7     PG     PE  0.114599  0.102962  0.126236   37.990110    0.001
8     PG     NM  0.194576  0.177889  0.211263   44.983792    0.001
9     PE     NM  0.079977  0.060023  0.099932   15.462023    0.001


region: 2

  group1 group2      Diff     Lower     Upper    q-value   p-value
0     PG     PE  0.112784  0.090186  0.135382  19.257935  0.001000
1     PG     NF  0.394821  0.377550  0.412091  88.209208  0.001000
2     PG     NM  0.231964  0.196678  0.267250  25.365611  0.001000
3     PG  NF_NM  0.345717  0.284822  0.406612  21.906186  0.001000
4     PE     NF  0.282037  0.255676  0.308397  41.283515  0.001000
5     PE     NM  0.119180  0.078663  0.159698  11.349691  0.001000
6     PE  NF_NM  0.232933  0.168865  0.297002  14.028543  0.001000
7     NF     NM  0.162856  0.125050  0.200663  16.621299  0.001000
8     NF  NF_NM  0.049103 -0.013286  0.111493   3.036894  0.200204
9     NM  NF_NM  0.113753  0.044188  0.183317   6.309576  0.001000


region: 3

  group1 group2      Diff     Lower     Upper    q-value  p-value
0     PE     NF  0.220098  0.189555  0.250641  27.805684    0.001
1     PE     PG  0.075298  0.056441  0.094156  15.407659    0.001
2     PE  NF_NM  0.227175  0.189939  0.264412  23.540711    0.001
3     PE     NM  0.102774  0.072096  0.133453  12.926478    0.001
4     NF     PG  0.295397  0.270151  0.320642  45.149714    0.001
5     NF  NF_NM  0.007077 -0.033768  0.047922   0.668555    0.900
6     NF     NM  0.117324  0.082354  0.152294  12.945575    0.001
7     PG  NF_NM  0.302474  0.269443  0.335504  35.334723    0.001
8     PG     NM  0.178073  0.152664  0.203482  27.042092    0.001
9  NF_NM     NM  0.124401  0.083455  0.165347  11.723028    0.001


region: 4

  group1 group2      Diff     Lower     Upper   q-value   p-value
0     PG     NM  0.134485  0.039138  0.229833  5.161890  0.001828
1     PG     NF  0.309217  0.108554  0.509879  5.639508  0.001000
2     PG     PE  0.129170  0.006137  0.252203  3.842236  0.035468
3     NM     NF  0.174731 -0.043902  0.393364  2.924820  0.166830
4     NM     PE  0.005315 -0.145257  0.155888  0.129194  0.900000
5     NF     PE  0.180047 -0.052001  0.412095  2.839564  0.188005
```

##### Boxplot -> forest naturalness classes¶

In [17]:

```
plots.plot_boxplots(
    svea_df, "naturalness", height=4, aspect=0.75, fig_title="b) Stand-level validation - Forest naturalness",
    save_name="Figure4b.png", width=0.5
)
```

In [18]:

```
plots.describe_ci(svea_df, "pred_mean", ["region", "naturalness"])
```

Out[18]:

|  |  | mean | count | std | ci95\_lo | ci95\_hi |
| --- | --- | --- | --- | --- | --- | --- |
| region | naturalness |  |  |  |  |  |
| 1 | No | 0.299558 | 34136 | 0.176607 | 0.297685 | 0.301432 |
|  | Yes | 0.571572 | 7654 | 0.204969 | 0.566980 | 0.576164 |
| 2 | No | 0.300205 | 6306 | 0.199128 | 0.295290 | 0.305120 |
|  | Yes | 0.638149 | 1683 | 0.225907 | 0.627356 | 0.648942 |
| 3 | No | 0.301697 | 6606 | 0.153444 | 0.297997 | 0.305397 |
|  | Yes | 0.514033 | 917 | 0.203989 | 0.500830 | 0.527236 |
| 4 | No | 0.308738 | 234 | 0.161656 | 0.288025 | 0.329451 |
|  | Yes | 0.435557 | 23 | 0.147576 | 0.375245 | 0.495870 |

In [19]:

```
plots.tukey_subsets(svea_df, "pred_mean", "naturalness", "region")
```

```
region: 1

  group1 group2      Diff     Lower    Upper     q-value  p-value
0    Yes     No  0.270034  0.265457  0.27461  163.557264    0.001


region: 2

  group1 group2     Diff     Lower     Upper    q-value  p-value
0     No    Yes  0.33828  0.327028  0.349533  83.339123    0.001


region: 3

  group1 group2      Diff     Lower     Upper    q-value  p-value
0     No    Yes  0.212149  0.200986  0.223312  52.684495    0.001


region: 4

  group1 group2      Diff     Lower     Upper   q-value  p-value
0     No    Yes  0.140161  0.070294  0.210028  5.589368    0.001
```

#### NFI validation¶

plot level validation (1 central pixel + 4 neighbours)

In [20]:

```
# load a dataframe with predictions and NFI atributes
nfi_df = pandas.read_csv("nfi_predictions.csv")

# at least 4 pixels needed
nfi_df = nfi_df[nfi_df.pred_count > 3]

nfi_df["region"] = pandas.Categorical(nfi_df.region)
nfi_df["naturalness"] = pandas.Categorical(nfi_df.naturalness, categories=[3, 1, 2])
nfi_df.naturalness.cat.rename_categories({
    1.0: "normal", 2.0: "natural", 3.0: "plantation"
}, inplace=True)

# drop rows with NA in natura2000 column
nfi_df.dropna(subset=["natura2000"], inplace=True)
nfi_df.natura2000.replace({0: "No", 1: "Yes"}, inplace=True)
nfi_df["natura2000"] = pandas.Categorical(nfi_df.natura2000)

# show the number of validation polygons per region
display(nfi_df.region.value_counts())
print("TOTAL PLOTS N: ", nfi_df.region.value_counts().sum())
```

```
2    5397
3    4627
1    3073
4     683
Name: region, dtype: int64
```

```
TOTAL PLOTS N:  13780
```

##### Boxplot -> forest naturalness classes¶

1=normal; 2=natural forest; 3=plantation forest

In [21]:

```
plots.plot_boxplots(
    nfi_df, "naturalness", height=4, aspect=0.75, fig_title="c) Plot-level validation - Forest naturalness",
    save_name="Figure4c.png", width=0.85
)
```

In [22]:

```
plots.describe_ci(nfi_df, "pred_mean", ["region", "naturalness"])
```

Out[22]:

|  |  | mean | count | std | ci95\_lo | ci95\_hi |
| --- | --- | --- | --- | --- | --- | --- |
| region | naturalness |  |  |  |  |  |
| 1 | plantation | 0.192474 | 19 | 0.143214 | 0.128077 | 0.256871 |
|  | normal | 0.367899 | 2898 | 0.246074 | 0.358940 | 0.376858 |
|  | natural | 0.771919 | 156 | 0.184178 | 0.743017 | 0.800822 |
| 2 | plantation | 0.265434 | 74 | 0.177943 | 0.224890 | 0.305977 |
|  | normal | 0.375867 | 5200 | 0.236511 | 0.369438 | 0.382295 |
|  | natural | 0.717572 | 123 | 0.240153 | 0.675131 | 0.760014 |
| 3 | plantation | 0.317090 | 103 | 0.182212 | 0.281901 | 0.352280 |
|  | normal | 0.402548 | 4498 | 0.195255 | 0.396842 | 0.408254 |
|  | natural | 0.557435 | 26 | 0.197770 | 0.481414 | 0.633455 |
| 4 | plantation | 0.346907 | 69 | 0.163826 | 0.308252 | 0.385563 |
|  | normal | 0.410985 | 612 | 0.169508 | 0.397555 | 0.424415 |
|  | natural | 0.534300 | 2 | 0.278176 | 0.148768 | 0.919832 |

In [23]:

```
plots.describe_ci(nfi_df, "pred_mean", ["region", "naturalness"])["count"].sum()
```

Out[23]:

```
13780
```

In [24]:

```
plots.tukey_subsets(nfi_df, "pred_mean", "naturalness", "region")
```

```
region: 1

    group1      group2      Diff     Lower     Upper    q-value   p-value
0   normal     natural  0.404020  0.357218  0.450823  28.626485  0.001000
1   normal  plantation  0.175425  0.044360  0.306490   4.438510  0.004881
2  natural  plantation  0.579446  0.441081  0.717810  13.887383  0.001000


region: 2

    group1      group2      Diff     Lower     Upper    q-value  p-value
0   normal     natural  0.341705  0.291254  0.392157  22.455176    0.001
1   normal  plantation  0.110433  0.045689  0.175178   5.655038    0.001
2  natural  plantation  0.452139  0.370778  0.533499  18.424489    0.001


region: 3

       group1      group2      Diff     Lower     Upper   q-value  p-value
0      normal  plantation  0.085458  0.039901  0.131015  6.219512    0.001
1      normal     natural  0.154886  0.064973  0.244799  5.711495    0.001
2  plantation     natural  0.240344  0.140010  0.340678  7.942292    0.001


region: 4

    group1      group2      Diff     Lower     Upper   q-value   p-value
0  natural      normal  0.123315 -0.158108  0.404738  1.455588  0.555019
1  natural  plantation  0.187393 -0.097614  0.472400  2.184135  0.271289
2   normal  plantation  0.064078  0.013619  0.114537  4.218434  0.008309
```

In [25]:

```
plots.plot_boxplots(
    nfi_df, "natura2000", height=4, aspect=0.75, fig_title="d) Plot-level validation - Natura 2000 habitat quality",
    save_name="Figure4d.png", width=0.5
)
```

In [26]:

```
plots.describe_ci(nfi_df, "pred_mean", ["region", "natura2000"])
```

Out[26]:

|  |  | mean | count | std | ci95\_lo | ci95\_hi |
| --- | --- | --- | --- | --- | --- | --- |
| region | natura2000 |  |  |  |  |  |
| 1 | No | 0.358022 | 2842 | 0.237456 | 0.349292 | 0.366753 |
|  | Yes | 0.747826 | 231 | 0.240334 | 0.716832 | 0.778819 |
| 2 | No | 0.367891 | 5216 | 0.229821 | 0.361654 | 0.374128 |
|  | Yes | 0.792772 | 181 | 0.213388 | 0.761684 | 0.823859 |
| 3 | No | 0.396336 | 4531 | 0.191968 | 0.390747 | 0.401926 |
|  | Yes | 0.646000 | 96 | 0.214814 | 0.603028 | 0.688972 |
| 4 | No | 0.399152 | 660 | 0.168239 | 0.386316 | 0.411987 |
|  | Yes | 0.569035 | 23 | 0.143409 | 0.510425 | 0.627644 |

In [27]:

```
plots.tukey_subsets(nfi_df, "pred_mean", "natura2000", "region")
```

```
region: 1

  group1 group2      Diff    Lower     Upper    q-value  p-value
0     No    Yes  0.389803  0.35792  0.421686  33.901392    0.001


region: 2

  group1 group2      Diff     Lower     Upper   q-value  p-value
0     No    Yes  0.424881  0.390895  0.458867  34.65976    0.001


region: 3

  group1 group2      Diff     Lower    Upper    q-value  p-value
0     No    Yes  0.249664  0.210748  0.28858  17.787008    0.001


region: 4

  group1 group2      Diff     Lower     Upper   q-value  p-value
0     No    Yes  0.169883  0.100124  0.239641  6.762208    0.001
```

In [ ]:
